# A *GFP* splicing reporter in a *coilin* mutant background reveals links between alternative splicing, siRNAs and coilin function in *Arabidopsis thaliana*

**DOI:** 10.1101/2022.12.18.520953

**Authors:** Tatsuo Kanno, Phebe Chiou, Ming-Tsung Wu, Wen-Dar Lin, Antonius Matzke, Marjori Matzke

**Affiliations:** Institute of Plant and Microbial Biology, Academia Sinica, Taipei, Taiwan

**Keywords:** alternative splicing, *Arabidopsis thaliana*, Cajal bodies, coilin, GFP, MEM1, posttranscriptional gene silencing, small RNAs, SMU2, WRAP53, ZC3HC1

## Abstract

Coilin is a scaffold protein essential for the structure of Cajal bodies, which are nucleolar-associated, nonmembranous organelles that coordinate the assembly of nuclear ribonucleoproteins (RNPs) including spliceosomal snRNPs. To learn more about coilin functions and pathways in plants, we conducted a genetic suppressor screen using a *coilin (coi1)* mutant in *Arabidopsis thaliana* and performed an immunoprecipitation-mass spectrometry analysis on coilin protein. The *coi1* mutations modify alternative splicing of a *GFP* reporter gene, resulting in a ‘hyper-GFP’ phenotype in young *coi1* seedlings relative to the intermediate wild-type level. As shown here, this hyper-GFP phenotype is extinguished in older *coi1* seedlings by posttranscriptional gene silencing triggered by siRNAs derived from aberrant splice variants of *GFP* pre-mRNA. In the *coi1* suppressor screen, we identified suppressor mutations in WRAP53, a putative coilin-interacting protein; SMU2, a predicted splicing factor; and ZC3HC1, an incompletely characterized zinc finger protein. These suppressor mutations return the hyper-GFP fluorescence of young *coi1* seedlings to the intermediate wild-type level. Additionally, *zc3hc1 coi1* mutants display more extensive *GFP* silencing and elevated levels of *GFP* siRNAs, suggesting the involvement of wild-type ZC3HC1 in siRNA biogenesis or stability. The immunoprecipitation-mass spectrometry analysis reinforced coilin’s roles in pre-mRNA splicing, nucleolar chromatin structure, and rRNA processing. Coilin’s participation in these processes, at least some of which incorporate small RNAs, supports the hypothesis that coilin acts as a chaperone for small noncoding RNAs. Our study demonstrates the usefulness of the *GFP* splicing reporter for investigating alternative splicing, ribosome biogenesis, and siRNA-mediated silencing in the context of coilin function.

## Introduction

Coilin is an intriguing protein that is present in most multicellular eukaryotes. Even after decades of research, the full range of coilin’s functions and mode of action are not fully understood (Machyna et al., 2015). Coilin is best known as an essential structural constituent and prominent marker of Cajal bodies (CBs). CBs are nonmembranous nuclear organelles that are usually situated adjacent to the nucleolus, a second membrane-free nuclear organelle that also contains detectable amounts of coilin (Hebert 2013; Trinkle-Mulcahy and Sleeman, 2017). CBs are a major site for maturation, processing, and quality control of small nuclear RNAs (snRNAs), including those that are present in small nuclear ribonucleoprotein particles (snRNPs) required for pre-mRNA splicing, and small nucleolar RNAs (snoRNAs), which guide processing and maturation of rRNAs and tRNAs in the nucleolus (Machyna et al., 2015).

Even though coilin appears to be particularly enriched in CBs and is required for their formation and maintenance, a large fraction of coilin protein in animal cells is distributed throughout the nucleoplasm (Lam et al., 2002) where its roles are less clear. Through dynamic associations with chromatin, nucleoplasmic coilin in animals is thought to contribute to a number of processes including not only pre-mRNA splicing but also chromatin organization, DNA repair and telomere maintenance (Machyna et al., 2015). Whether coilin also performs these multiple functions in plants is not yet clear.

The requirement for coilin to maintain CB structural integrity is illustrated by the ability of coilin mutations to disrupt CBs in animal cells (Liu et al., 2009; Carrero et al., 2011: Machyna et al., 2015) and in *Arabidopsis thaliana* (Arabidopsis) (Collier et al., 2006: Kanno et al., 2016). Mutations in coilin lead to embryonic lethality or semi-lethality in vertebrates (Strzelecka et al., 2010; Walker *et al*., 2009) whereas they have minimal effects on development in Drosophila (Liu et al., 2009) and Arabidopsis (Collier et al., 2006; Kanno et al., 2016; Abulfaraj et al., 2022). In both animals and plants, splicing defects have been observed in coilin-deficient mutants, consistent with a role for coilin in pre-mRNA splicing (Walker et al., 2009; Strzelecka et al., 2010: Kanno et al., 2016; Abulfaraj et al., 2022). Expression of stress- and plant immunity-related genes is often altered in coilin mutants in Arabidopsis (Kanno et al., 2016; Abulfaraj et al., 2022), indicating the importance of coilin for plant protective functions.

The ability of coilin to facilitate CB assembly depends on self-associations and interactions with RNA and other CB proteins (Makarov et al., 2013; Machyna et al., 2015). Conserved regions of coilin protein in plants and animals include an N-terminal self-association domain; a C-terminal atypical tudor domain that mediates interactions with several snRNPs (Herbert et al., 2001; Xu et al., 2005); and a central disordered domain (Makarov et al., 2013). Although coilin lacks conventional RNA binding motifs, analysis of the secondary structure of Arabidopsis coilin revealed several degenerate RNA recognition motifs in the N-terminal region and central disordered domain (Makarov et al., 2013; Machyna et al., 2015). The reported ability of coilin to bind RNAs, particularly small noncoding RNAs such as snRNAs and snoRNAs (Machyna et al., 2014), led to the proposal that coilin may serve as a chaperone for trafficking of nuclear noncoding small RNAs (Machyna et al., 2015).

The possible involvement of CBs and coilin in small interfering RNA (siRNA)-induced gene silencing (RNA silencing) has been explored in plants, where 21-24-nt siRNAs generated by dicer cleavage of double-stranded RNA (dsRNA) precursors mediate both posttranscriptional and transcriptional gene silencing (PTGS and TGS, respectively). DsRNAs can be produced by RNA-dependent RNA polymerases (RDR) acting on single stranded RNAs that are perceived by the cell as aberrant in some way. Aberrant RNAs can include ‘foreign’ RNAs derived from viruses and transgenes as well as endogenous RNAs that are untranslatable due to premature termination codons resulting from single nucleotide mutations, missplicing, or lack of a poly-A tail. PTGS involves sequence-specific degradation of target RNAs triggered by complementary 21-22-nt siRNAs, which are produced in Arabidopsis by DICER-LIKE4 (DCL4) and DCL2, respectively. TGS is associated with DNA methylation of promoter segments guided by DCL3-dependent 24-nt siRNAs in a process referred to as RNA-directed DNA methylation (RdDM) (Uslu and Wassenegger, 2020; El-Sappah et al., 2021; Kryovrysanaki et al, 2022)

In Arabidopsis, factors required for PTGS and TGS, such as DCL enzymes and ARGONAUTE (AGO) silencing effector proteins, have been detected in CBs by immunofluorescence microscopy (Li et al., 2006; Pontes and Pikaard, 2008; Pontes et al., 2013). However, as shown by experiments using an Arabidopsis *ncb* (*no cajal body*) mutant (Collier et al., 2006), intact CBs do not seem to be required for siRNA-mediated gene silencing to occur. Neither RdDM nor production of siRNAs at several tested loci was substantially affected by the loss of CB integrity in *ncb* mutants, although the efficiency of RdDM at these sites may have been somewhat reduced (Li et al., 2008).

Our interest in coilin was stimulated by the retrieval of multiple *coilin* (*coi1*) mutants in a forward genetic screen based on an alternatively-spliced *GFP* reporter gene in Arabidopsis (Kanno et al., 2016; Kanno et al., 2020). In this screen, *coi1* mutants were initially identified in young seedlings (shortly after germination) by their hyper-GFP phenotype relative to wild-type seedlings, which display an intermediate level of GFP fluorescence (Kanno et al., 2020). Although the basis of the hyper-GFP phenotype in young *coi1* seedlings is not yet completely understood, it can be attributed at least in part to modified splicing of *GFP* pre-mRNA, leading to elevated amounts of the only *GFP* splice variant that can be translated into protein (Kanno et al., 2016; Kanno et al., 2020).

After publishing the identification of the hyper-GFP *coi1* mutants (Kanno et al., 2016), we later observed that in many older *coi1* seedlings (second to fourth true leaf stage), that abrupt silencing of the *GFP* reporter gene occurs. The silencing pattern, featuring dark red leaves emerging from the shoot apex of a hyper-GFP plantlet, superficially resembled virus recovery, in which asymptomatic leaves appear during growth of an infected plant. Recovery from virus infection is known to involve PTGS that is provoked by virus-derived siRNAs (Goshal and Sanfacon H, 2015: Uslu and Wassenegger, 2020).

Here we report the results of experiments designed to investigate the silencing/’recovery’ phenomenon in older *coi1* mutants. We also describe findings from genetic and biochemical approaches that were carried out to identify proteins that may be functionally linked to coilin and potentially illuminate new coilin-dependent pathways.

## MATERIALS AND METHODS

### Plant material and growth conditions

Wild-type and mutant plants used in this study were in the ecotype Col-0 background and cultivated under long-day conditions (22-23°C, 16 hours light, 8 hours dark). Gene names are indicated in italics and proteins in upright letters. Mutant names are indicated in lower case letters; wild-type names in upper case. To test for the occurrence of PTGS, *rdr6-14* and *dcl4-12* alleles were introduced into the *WT T* line and into various mutants by intercrossing. For assessments of GFP fluorescence intensity in seedlings, seeds of the desired line were surface sterilized, germinated on solid Murashige and Skoog (MS) medium in plastic Petri dishes and cultivated in a growth incubator under a 16-hour light/8-hour dark cycle at 24° C. Seedlings were observed daily for fluorescence levels under a Leica fluorescence stereo-microscope over a period of 4-6 weeks.

Seeds of the *WT T* line and coilin suppressor mutants as well as other mutants used in this study are available from the Arabidopsis Biological Research Center (ABRC) under the ABRC seed stock numbers listed in **Table S1**. Accession numbers for RNAseq, small RNAseq and DNAseq data presented in this manuscript are also listed in **Table S1**.

### Forward genetic screen and identification of coilin suppressor mutants

The protocol for forward genetic screens based on an alternatively-spliced, intron-containing *GFP* reporter gene in Arabidopsis [*T* line; referred to herein as ‘wild-type’ (*WT T*)] has been described previously (Kanno et al, 2016; 2017a, b; 2018). In the present coilin suppressor screen, approximately 40,000 Arabidopsis seeds of a homozygous *coi1-8* mutant (also homozygous for the *T* locus) were treated with ethyl methane sulfonate (EMS) and sown on soil (M1 generation). From approximately 20,000 M1 plants that grew to maturity and produced self-fertilized seeds, 52 batches of M2 seeds (the first generation when a recessive mutation can be homozygous) were harvested. M2 seeds were surface sterilized, germinated on solid MS medium in plastic Petri dishes and cultivated in a growth incubator as described above.

Putative suppressor mutants were identified by examining M2 seedlings under a Lecia fluorescence stereo-microscope approximately seven days after germination for reduced GFP fluorescence relative to the hyper-GFP phenotype of the *coi1-8* single mutant. Around 100 putative suppressor mutants were recovered and grouped into two main categories: GFP-weak (GFP fluorescence reduced from hyper-GFP level but still microscopically visible) and GFP-negative (GFP-neg) (microscopically visible GFP fluorescence eliminated; seedlings appear deep red, owing to autofluorescence of chlorophyll at the excitation wavelength of GFP). Sequencing of the *GFP* reporter gene in the GFP-neg mutants revealed that five harbored a previously unidentified loss-of-function mutation in the GFP protein coding region (**named in Table S1; described in Figure S1**; Fu et al., 2015).

GFP-weak mutants that contained a wild-type *GFP* nucleotide sequence were tested to confirm: (1) stable meiotic inheritance of the GFP-weak phenotype; (2) the presence of the expected *coi1-8* mutation in the endogenous *coi1* gene; and (3) the absence of any second-site intragenic suppressor mutations induced by the EMS treatment into the *coi1-8* gene sequence. GFP-weak mutants that passed these additional tests were then subjected to next generation mapping (NGM) (James et al. 2013; Kanno et al., 2020) and/or candidate gene sequencing to identify the respective causative suppressor mutations. Mutations were confirmed by the identification of multiple alleles and/or complementation analyses. All mutations reported here are recessive.

### Western blotting using a GFP antibody

Western blotting to determine the GFP protein levels in the *coi1 wrap53, coi1 smu2* and *coi1 zc3hc1* double mutants compared to the *coi1* single mutant and *WT T* line was performed as described in previous publications (Fu *et al*., 2015). For a loading control, a duplicate gel containing the same samples was run and stained with Coomassie Brilliant blue.

### RNA and small RNA sequencing (RNAseq and sRNAseq)

Total RNA was prepared using a Plant Total RNA Miniprep kit (GMbiolab, Taiwan) from two-week-old seedlings of the *WT T* line and the *coi1 wrap53, coi1 smu2* and *coi1 zc3hc1* double mutants cultivated on solid MS medium as described above. RNA concentrations were assessed by NanoDrop (ND-1000 spectrophotometer). Library preparation, RNAseq and sRNAseq were performed (biological triplicates for each sample) as described previously by an in-house Genomic Technology Core facility (Kanno *et al*., 2016). Whole genome re-sequencing of the *coi1-8 wrap53, coi1-8 smu2* and *coi1-8 zc3hc1* mutants was carried out to identify any remaining EMS-induced second-site mutations that alter splice sites. These mutations were then excluded from the analysis of alternative splicing.

### Analysis of RNAseq and sRNAseq data

Pair-ended RNAseq reads were mapped to the Arabidopsis TAIR10 genome in two steps. The first step was to map reads to a combined transcriptome database of Araport11 (Cheng *et al*., 2017) and Atrtd2 (Zhang *et al*., 2017) using Bowtie2 (Langmead and Salzberg, 2012), where only ungapped alignments of read pairs been mapped to the same transcripts were accepted. Coordinates of these transcriptome-mapping alignments were transferred to the genome. The second step was to map rest reads to the TAIR10 genome using BLAT (Kent, 2002) directly.

To gain alignments of small RNA reads to the GFP-containing vector pAJM-EPRV (GenBank accession: HE582394.1) as precise as possible, the following prioritized approach was adopted: (i) reads were mapped to the canonical GFP transcript, only perfect matches were accepted, (ii) for splicing junctions supported by RNAseq data, reads were mapped to junction-spanning fragments and only perfect matches were accepted, (iii) repeat above two steps by allowing one mismatch, and (iv) rest reads were mapped to pAJM-EPRV using BLAT with a set of fine scanning parameters (-minMatch=1 -minScore=14 -tileSize=9) and allowing one mismatch.

Alternative splicing (AS) detection for intron-retention (IR), exon-skipping (ES), and alternative donor/acceptor (altDA) was done using a similar method as described in our previous publications (Kanno *et al*., 2017a, b; 2018). For each kind of AS event, read-based counts that are supporting the AS event from the control and treatment samples were considered as signals, where read-based counts that are related but not supporting the event were considered as backgrounds. In practice, intron read depths, counts of splicing reads skipping exon(s), and counts of reads exactly supporting one donor-accepter pair were taken as the signals for IR, ES, and altDA events, respectively. For backgrounds, read depths of neighboring exons, counts of splicing reads involving one skipped exon, and counts of reads not supporting the donor-accepter pair were taken for IR, ES, and altDA events, respectively. Signals from the control and treatment samples were then compared to the backgrounds using chi-squared test for goodness-of-fit for discovering differential preference of AS events in the two samples. In this study, an AS event was considered as significant if its P-value is below 0.05.

### Protein immunoprecipitation (IP)of epitope-tagged coilin proteins

Total proteins were extracted from two-week-old seedlings of coilin-mRed, FLAG-coilin and coilin-FLAG transgenic plants. Protein extraction and immunoprecipitation were carried out according to a published protocol (Huang et al., 2016). Briefly, five grams of seedlings were ground to a powder in liquid nitrogen. After grinding, 14 ml of SPII+ buffer [100 mM Na-phosphate, pH 8.0, 150 mM NaCl, 5mM EDTA, 5mM EGTA, 0.1% NP-40, 1mM PMSF, 5μM MG132, 1x Phosphatase inhibitor cocktail 2 (Sigma-Aldrich), 1x Phosphatase cocktail 3 (Sigma-Aldrich) and a protease inhibitor (cOmplet EDTA-free Protease Inhibitor Cocktail, Roche) were added to the ground sample. The solution was sonicated followed by centrifugation (two times at 20,000 g for 10 min at 4^°^C) to remove insoluble cell debris. The supernatant constituted the protein extract that was used for immunoprecipitation.

One hundred microliters of Anti-FLAG M2 Magnetic Beads (Millipore) or RFP-trap magnetic beads (ChromoTek) were washed three times with 1 ml of SII buffer and then added to the protein extract. The whole solution was incubated at 4^°^C for one hour with 20 rpm agitation. Following this step, the beads were collected using a magnetic stand and washed three times with 10 ml of SPII buffer. The bound proteins were eluted by adding elution buffer [100mM Na-phosphate, pH 8.0, 150mM NaCl, 0.05% Triton X-100, 500μg/ml 3x FLAG peptide (Millipore)] for FLAG-IP samples or, for mRed-IP sample, according to the manufacturer’s instruction, incubating the beads with 0.2 M glycine pH 2.5 solution followed by add 1M Tris base pH 10.4 for neutralization.

### LC-MS/MS analysis and data processing

Digested peptides were injected into a Thermo Scientific Ultimate 3000 coupled with a Thermo Scientific Q-Exactive Mass Spectrometer. The samples were run in a linear 90 min gradient at a flow rate of 300 nL/min. The raw files were searched against the Araport_11 database using Proteome Discoverer (version 2.2). The peptide precursor mass tolerance was set at 10 ppm, and MS/MS tolerance was set at 0.02 Da. The false discovery rate (FDR) at protein and peptide level was set at 1%. The variable modification was set as oxidation on Methionine resides, and the cysteine carbamidomethylation was set as a static modification. The mass spectrometry proteomics data have been deposited to the ProteomeXchange Consortium via the PRIDE [1] partner repository with the dataset identifier PXD027146”.

## RESULTS AND DISCUSSION

### 1. Investigation of GFP silencing/recovery in older coi1 seedlings

#### Characteristics of *GFP* silencing/recovery

The *coi1* mutants were identified in a previous forward genetic screen using a ‘wild-type’ *T* line (*WT T*) containing an alternatively-spliced *GFP* reporter gene in Arabidopsis (**Figure 1**) (Kanno et al., 2016; Kanno et al., 2020). *WT T* seedlings display an intermediate level of GFP fluorescence relative to two possible extremes: GFP-weak and hyper-GFP (Kanno et al., 2020). The *coi1* mutants can be distinguished from wild-type in young seedlings by their hyper-GFP phenotype (**Figure 2A, B, C; 8 DAG**) (Kanno et al., 2016).

**Figure 1:**
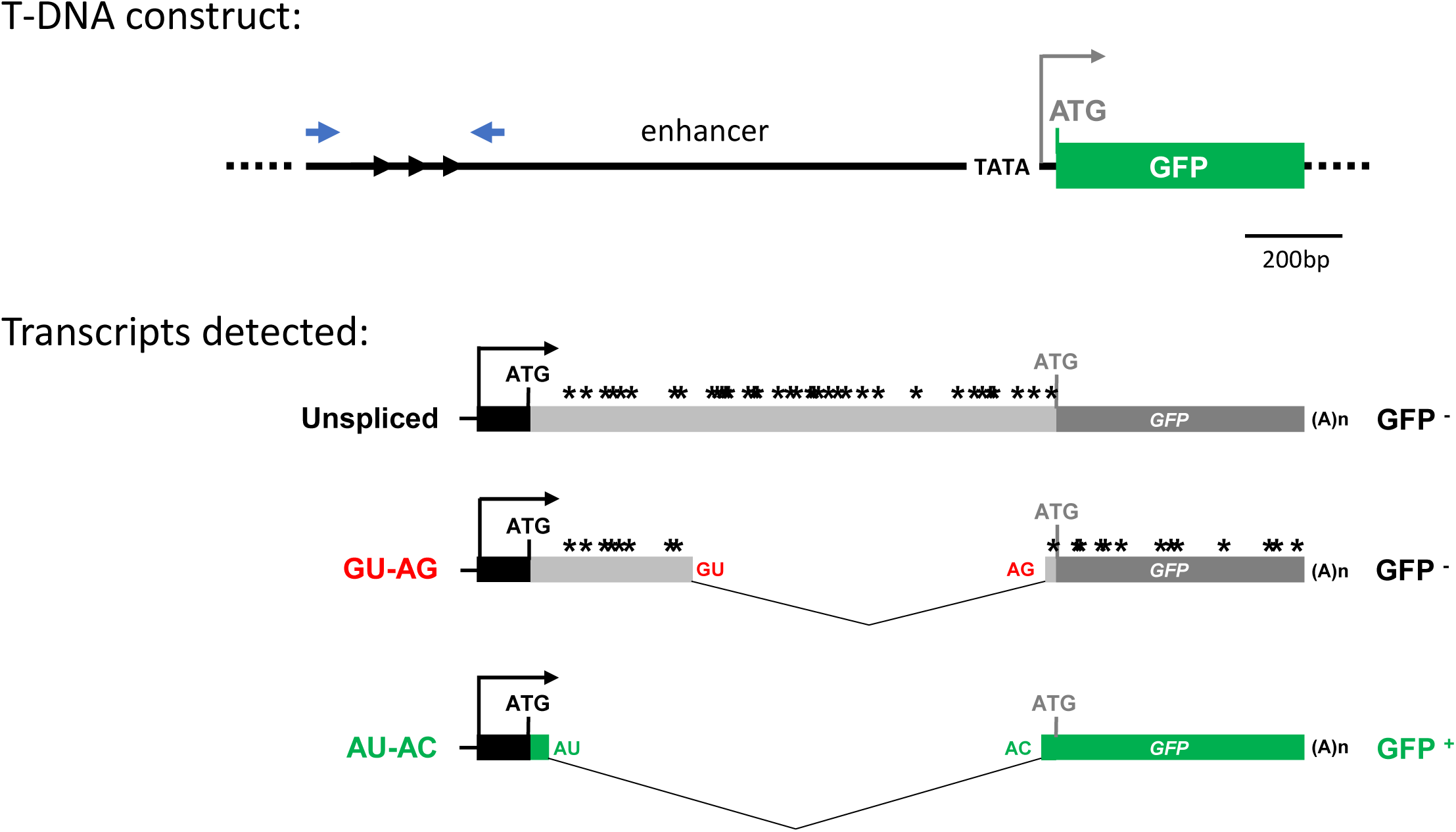
Alternative splicing of *GFP* reporter gene. <u>T-DNA construct:</u> One complete copy of the indicated T-DNA construct is integrated into the genome of *WT T* plants. An endogenous pararetrovirus enhancer is upstream of a minimal 35S promoter (TATA) and the anticipated GFP protein coding sequence (CDS; **long green bar**) The three black arrowheads at the 5’ end of the enhancer represent a short tandem repeat, which becomes methylated in the presence of homologous siRNAs (Kanno et al., 2008). Opposing blue arrowheads denote positions of primers used to detect DNA methylation (**Table S3**). Enhancer region, 1277 bp; minimal promoter, 91 bp; expected GFP coding region, 720 bp, Total length, 2088 bp. <u>Transcripts detected:</u> Although transcription was predicted to start downstream of the TATA minimal promoter, this promoter and the downstream ATG start codon remain unused (**gray letters**). Instead, transcription initiates at a cryptic promoter upstream in the tandem repeats (**black arrow**). Three polyadenylated *GFP* transcripts can be detected: an unspliced form (*GFP* pre-RNA) and two spliced variants, which contain canonical (GU-AG) and noncanonical (AU-AC; *GFP* mRNA) splice sites, respectively (Kanno et al., 2008). Owing to the presence of stop codons (asterisks) in the unspliced and GU-AG transcripts, only the AU-AC transcript is translated into GFP protein (**green bar**). Variations in the proportions of the three splice variants by alternative splicing modify the level of GPF fluorescence (Kanno et al., 2020). Translation initiates from a new upstream in-frame ATG start codon (**black letters**), which results in a 27 amino acid N-terminal extension to standard GFP protein (**short green bar**) (Fu et al., 2013).

**Figure 2:**
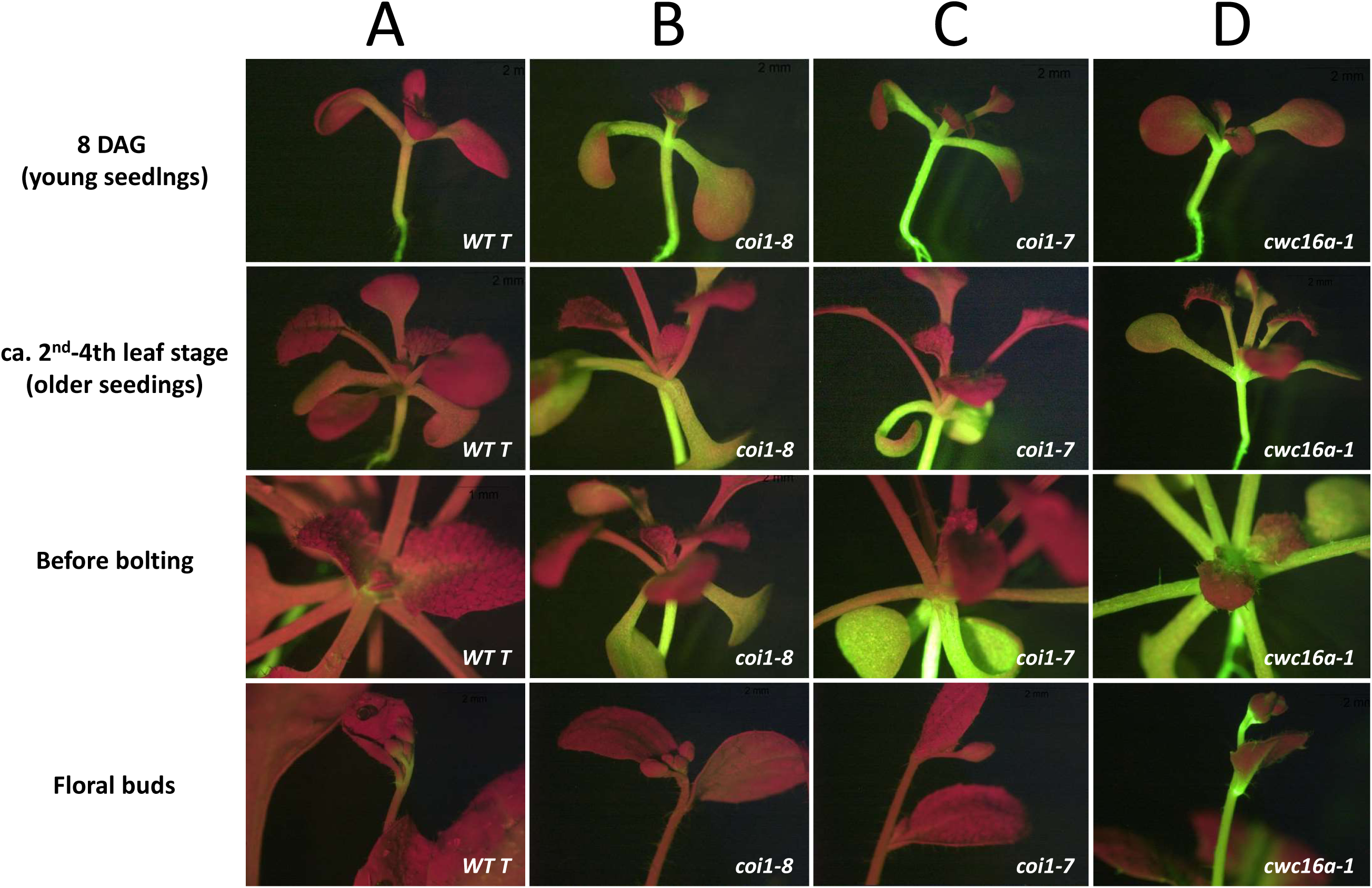
Age-dependent *GFP* silencing/recovery in *coi1* single mutants. **(A-C) Row 1:** ‘Young’ seedlings (8 days after germination, DAG) of *coi1* single mutants [***coi1-7*** and ***coi1-8***; **Figure S5**] display hyper-GFP phenotypes relative to the *WT T* line, which has an intermediate level of GFP fluorescence. **Rows 2, 3**: In ‘older’ *coi1* seedlings (2^nd^ to 4th true leaf stage), abrupt silencing/recovery, typified by dark red leaves emerging from a hyper-GFP stem, can occur. **Row 4:** Silencing persists into the floral stage. **A, all rows:** Most *WT T* plants also continue to show faint but visible accumulation of GFP fluorescence into the floral stage. **(A, all rows). (D, all rows):** The hyper-GFP *cwc16a* mutant (Kanno et al., 2017a; 2020) does not display silencing/recovery but remains hyper-GFP during the entire lifetime of the plant.

After publication of these results (Kanno et al., 2016), we subsequently found that later in development - typically the second to fourth true leaf stage (referred to here as ‘older’ seedlings, possibly coincident with the juvenile-to-adult transition) – 40-90% of *coi1* seedlings spontaneously undergoes a *GFP* silencing phenomenon characterized by the abrupt emergence of uniformly dark red leaves from a hyper-GFP shoot (**Figure 2B, C, older seedlings**). To distinguish this sudden, non-spreading silencing pattern from other silencing phenomena, we refer to it herein as ‘silencing/recovery’ (**Figure 2B, C, before bolting**). Silencing/recovery in older *coi1* seedlings persists in adult plants through the reproductive stage (**Figure 2B, C, floral buds**).

#### *coi1* mutants and *WT T* seedlings contain small RNAs likely derived from aberrant but not translatable *GPF* transcripts

To understand the silencing/recovery phenomenon, we tested whether it was associated with *GFP* siRNAs. We sequenced small RNAs in *coi1* mutant seedlings and as a control, in *WT T* seedlings, which we assumed would lack *GFP* small RNAs because there was no prior evidence of PTGS in this line. *GFP* small RNAs of both sense and antisense orientations were indeed detected, not only in the *coi1* single mutant (**Figure 3A**) but also, unexpectedly, in *WT T* plants displaying intermediate levels of GFP fluorescence (**Figure 4A**).

**Figure 3:**
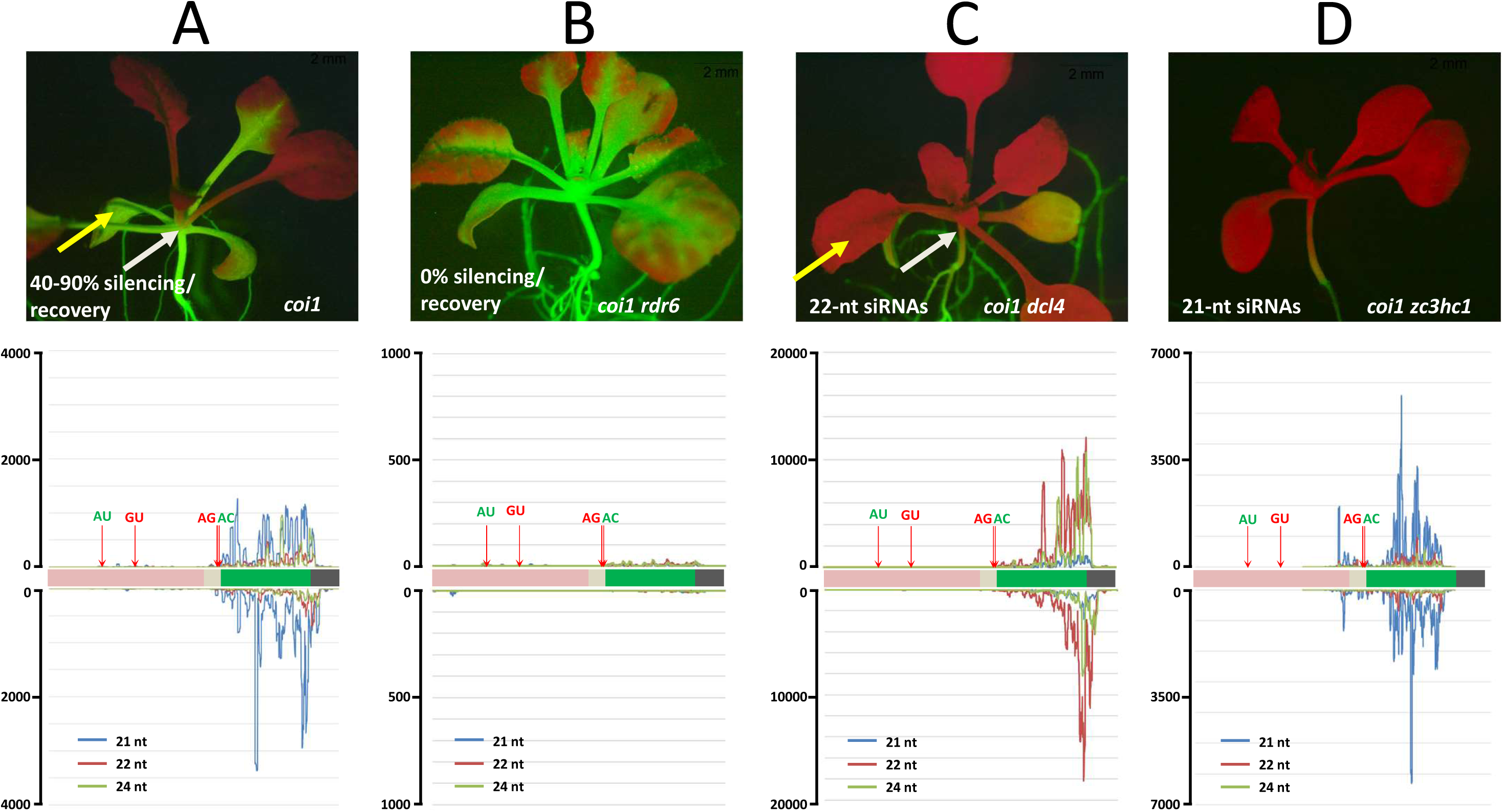
Silencing/recovery in *coi1* mutants is due to siRNA-mediated PTGS. **(A)** <u>Top</u>: silencing/recovery in *coi1* single mutant. Hyper-GFP fluorescence is extinguished at the 2^nd^-4^th^ true leaf stage and does not spread below the site of initiation (**white arrow**). <u>Bottom</u>: *GFP* siRNAs, predominantly 21-nt in length, are distributed throughout the GFP CDS (**green bar**) and more sparsely, in the upstream region flanked by AU and GU splice sites. **(B)** <u>Top</u>: *coi1 rdr6* double mutants remain hyper-GFP throughout the plant’s lifetime and *GFP* siRNAs are essentially eliminated (<u>Bottom</u>). **(C)** <u>Top:</u> *coi1 dcl4* double mutants display premature silencing that is already visible in primary true leaves (**yellow arrow**), which remain mostly hyper-GFP in the single *coi1* mutant (**A, yellow arrow**). Unlike the non-spreading silencing/recovery in a *coi1* single mutant (**A, white arrow**), *GFP* silencing in *coi1 dcl4* double mutants extends into primary leaves, shoot apex and the top of the hypocotyl (**C, white arrow**) coincident with a large increase of 22-nt *GFP* siRNAs (<u>Bottom</u>). (**D**) *coi1 zc3hc1* double mutants exhibit a silencing phenotype similar to that observed in *coi1 dcl4* but the size class of the predominant *GFP* siRNA differs: 22 nt in *coi1 dcl4* vs. 21 nt in *coi1 zc3hc1*. Seedlings were photographed at approximately 21 DAG. Length and distribution of *GFP* siRNAs along the GFP CDS and upstream region are indicated. Y axes show read number (scale differs in each graph). Quantitative comparative differences are shown in **Figure S3**.

**Figure 4:**
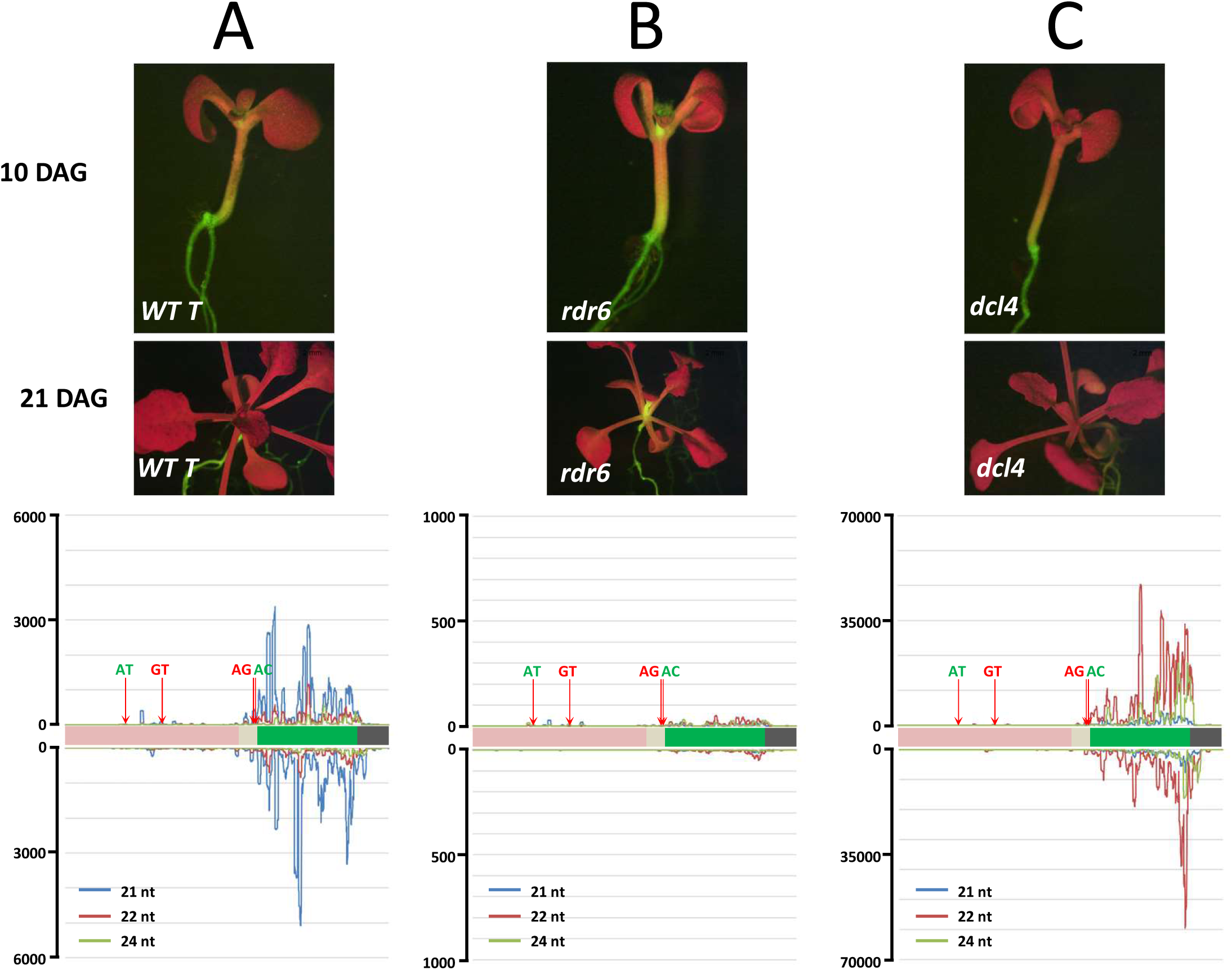
Pre-existing intermediate PTGS of *GFP* in *WT T* line. **(A)** <u>Top</u>: The *WT T* line shows intermediate GFP fluorescence that is highest in the shoot apex, hypocotyl and portions of the root (Daxinger et al., 2009). <u>Bottom</u>: *GFP* siRNAs in *WT T* are predominantly 21-nt in length (blue lines) and likely originate from the spliced, untranslatable GU-AG transcript (**Figure S2**). **(B)** <u>Top</u>: an *rdr6* mutation introduced into the *WT T* line increases GFP fluorescence in the shoot apex and hypocotyl and eliminates nearly all *GFP* siRNAs (<u>Bottom</u>) **(C)** <u>Top:</u> a *dcl4* mutation results in diminished GFP fluorescence that gradually spreads into the shoot apex and hypocotyl. <u>Bottom</u>: 22-nt siRNAs (red lines) are substantially increased in a *dcl4* mutant background while 21-nt siRNAs are greatly reduced. Seedlings were photographed at 10 and 21 DAG. Length and distribution of *GFP* siRNAs along the GFP CDS and upstream region are depicted. Y axes show approximate read number (scale differs in each graph). Quantitative comparative differences are shown in **Figure S2**.

In both the *coi1* mutant (**Figure 3A**) and *WT T* plants (**Figure 4A**), the *GFP* small RNAs were predominantly 21-nt in length, with lesser amounts of 22 and 24-nt small RNAs. Although the siRNAs were primarily derived from the GFP protein coding sequence, much lower amounts of mainly 21-nt *GFP* small RNAs could be reproducibly detected upstream of the GFP coding region, particularly from an exon sequence present in the spliced GU-AG and unspliced transcripts but not in the translatable AC-AU variant (**Figure S2, blue box**). Neither of the former two transcripts is translatable owing to the presence of numerous stop codons (**Figure 1**), which could potentially channel them into a pathway of siRNA production and PTGS (Liu and Chen, 2016). However, the relative paucity of siRNAs from the unique central region of the unspliced transcript (**Figure S2, red dotted box**) suggests that the spliced GU-AG transcript is the primary source of upstream *GFP* small RNAs. The substantial quantitative reduction in *GFP* small RNAs in the upstream sequence compared to the GFP coding region may reflect decreased activity of RDR6 as it progresses 3’ to 5’ along the substrate RNA (Moissiard et al., 2007; **Figure S2**).

Strong evidence that the abundant small RNAs originating from the GFP protein coding region also probably originate from aberrant *GFP* transcripts and not the translatable AU-AC RNA is provided by the *cwc16a* mutant (**Figure 2D**), which was identified by its hyper-GFP phenotype in the same splicing screen as the *coi1* mutant (Kanno et al., 2017a, 2020). The *cwc16a* mutant, which is defective in a putative step 2 splicing factor, produces almost exclusively the spliced, translatable AU-AC transcript (Kanno et al., 2017a), and in agreement with the hypothesis that *GFP* small RNAs are not derived from this transcript, *cwc16a* is essentially devoid of *GFP* small RNAs in the GFP protein coding and upstream regions (**Figure 5A; Figure S3).** Consistent with the lack of *GFP* siRNAs, the *cwc16a* mutant has not been observed to undergo *GFP* silencing/recovery (**Figure 2D**).

**Figure 5:**
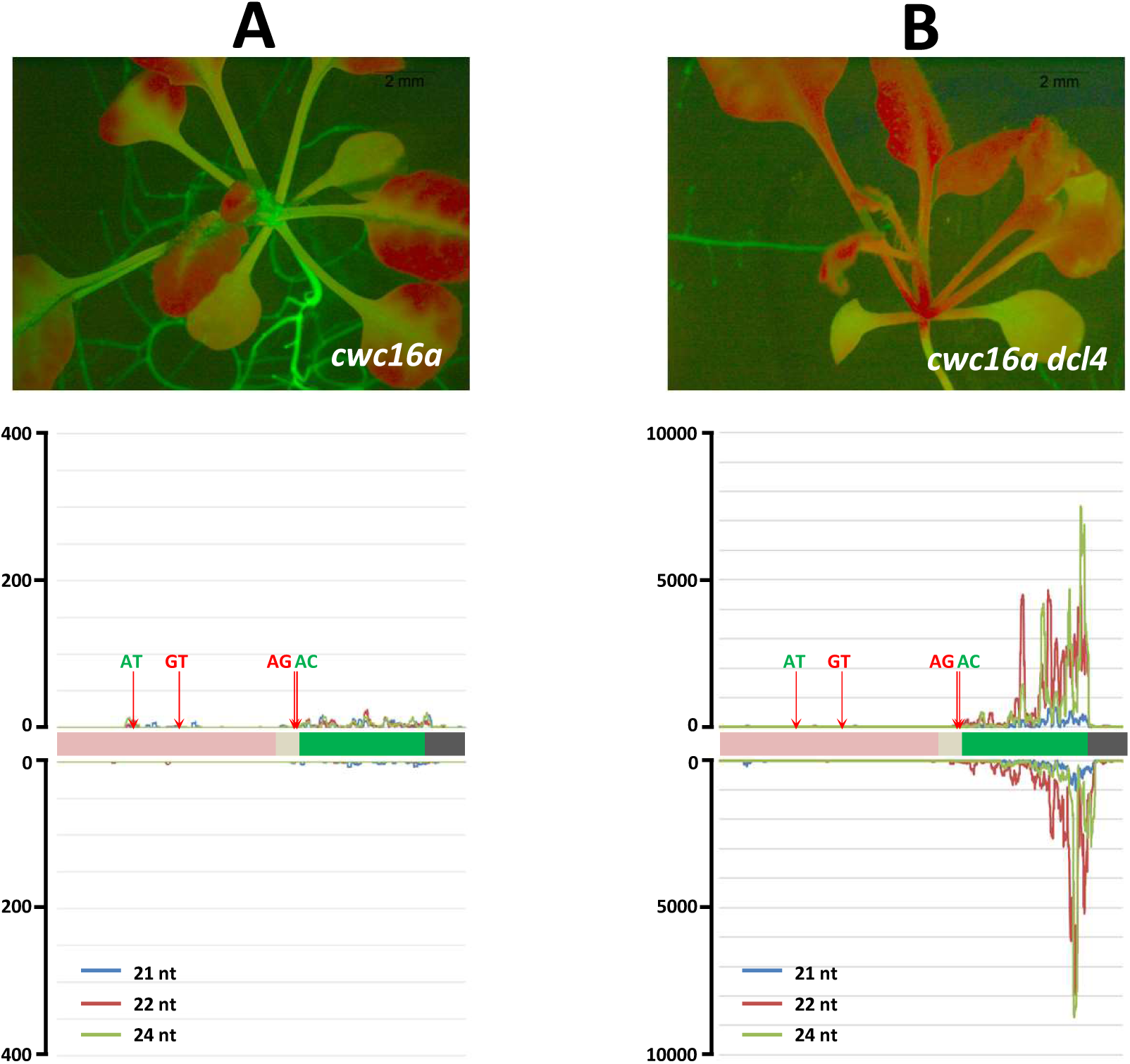
Silencing/recovery do not occur in a single *cwc16a* mutant, which lacks *GFP* siRNAs. **(A)** <u>Top:</u> *cwc16* mutants do not undergo silencing/recovery, remaining hyper-GFP throughout the plant’s lifetime, and (<u>Bottom</u>) contain only negligible amounts of *GFP* siRNAs. **(B)** <u>Top:</u> *cwc16a dcl4* double mutants undergo silencing that extends into the shoot apex and top of hypocotyl, and they produce large quantities of 22-nt *GFP* siRNAs (<u>Bottom</u>) owing to strong DCL2 activity in the absence of DCL4. Seedlings were photographed at approximately 21 DAG. Length and distribution of *GFP* siRNAs along the GFP CDS and upstream region are depicted. Y axes show approximate read number (scale differs in each graph). Quantitative comparative differences are shown in **Figure S3**.

The collective findings support the idea that the translatable AU-AC transcript is largely excluded from the small RNA biogenesis pathway. Among the three *GFP* splice variants detected (**Figure 1**), the untranslatable GU-AG transcript, and perhaps to a lesser extent, the unspliced transcript, thus represents the most credible sources of all *GFP* small RNAs in this system (**Figure S2**).

#### *GFP* small RNAs are bona fide siRNAs that trigger *GFP* silencing/recovery

To determine whether the *GFP* small RNAs are authentic siRNAs capable of triggering PTGS-of the *GFP* reporter gene, we crossed a *coi1* mutant and *WT T* plants with mutants defective in the PTGS factors RNA-DEPENDENT RNA POLYMERASE 6 (RDR6), which copies aberrant RNAs to produce dsRNAs, and DICER-LIKE4 (DCL4), which cleaves dsRNAs into 21-nt siRNAs (Lopez-Gomollon and Baulcombe, 2022). As described below, these experiments not only implicated classical PTGS in *GFP* silencing/recovery in older *coi1* mutants, but they also revealed a previously unsuspected, moderate level of PTGS in *WT T* seedlings.

Relative to the single *coi1* mutant at the silencing/recovery stage (**Figure 3A**), *coi1 rdr6* seedlings displayed increased GFP fluorescence and the loss of virtually all size classes and orientations of *GFP* small RNAs (**Figure 3B).** Notably, silencing/recovery similar to that observed in older *coi1* single mutants (**Figure 3A; Figure 2B, C**) was never observed in *coi1 rdr6* seedlings, which remained hyper-GFP throughout growth and development (**Figure 3B**).

Conversely, *coi1 dcl4* seedlings showed substantially lower *GFP* expression compared to the *coi1* single mutant (**Figure 3C**). However, unlike silencing/recovery in older *coi1* seedlings, which does not spread below the site of initiation (**Figure 2B, C; Figure 3A**), diminished *GPF* expression in *coi1 dcl4* seedlings occurs early and spreads into the shoot apex and the top part of hypocotyl (compare **Figure 3A and Figure 3C, white arrows**). The premature and more extensive *GFP* silencing in *coi1 dcl4* seedings was associated with a substantial reduction in 21-nt *GFP* small RNAs and a massive increase in 22-nt *GFP* small RNAs (**Figure 3C; Figure S3C**). Previous work has shown that a DCL4 deficiency results in enhanced DCL2 activity and increased production of 22-nt siRNAs, which can facilitate the spread and intensification of silencing (Mlotshwa et al., 2008; Chen et al., 2010; Parent et al., 2015; Taochy et al., 2017).

Similar changes were seen when *rdr6* and *dcl4* mutations were introduced into the *WT T* line: *rdr6* mutations enhanced GFP fluorescence and eliminated *GFP* small RNAs; *dcl4* mutations diminished GFP fluorescence and increased the accumulation of 22-nt small RNAs (**Figure 4B and C, respectively**).

Greatly elevated levels of 22-nt *GFP* siRNAs and extensive *GFP* silencing were also observed in *cwc16a dcl4* double mutants (**Figure 5C**). This finding indicates that the *cwc16a* mutant, which normally does not produce *GFP* siRNAs or exhibit PTGS, is nevertheless capable of undergoing PTGS provided sufficient numbers of siRNAs, such as the elevated level of 22-nt siRNAs in a *cwc16a dcl4* double mutant, are available.

#### Summary and limitations of our analysis of silencing/recovery in *coi1* mutants and *GFP* siRNAs

The results of experiments with the *rdr6* and *dcl4* mutants demonstrated that the abrupt, non-spreading silencing/recovery phenotype observed in older *coi1* single mutants is due to canonical PTGS triggered by 21-22-nt *GFP* siRNAs, which likely originate largely from the aberrant GU-AG *GFP* splice variant. What remains unclear is why silencing/recovery is provoked suddenly at a relatively late period of seedling development and not earlier in young *coi1* seedlings. Despite the presence of detectable amounts of 21-22-nt *GFP* siRNAs in young *coi1* seedlings, they consistently display a hyper-GFP phenotype prior to the sudden emergence of silencing/recovery in older *coi1* seedlings (**Figure 2**).

One possibility is that the threshold number of siRNAs required to effectively induce PTGS is only surpassed in older *coi1* seedlings, perhaps due to enhanced transcription of aberrant *GFP* splice variants at that time. It is currently difficult to assess this hypothesis because RNA-seq data are currently available only from two-week-old seedlings. Although the proportions of the three *GFP* splice variants and 21-22-nt *GFP* siRNAs differ to some extent between the *WT T* line and the *coi1* mutant at two weeks (**Figures S3 and S4)**, these quantitative differences affect primarily the translatable AU-AC splice variant **(Figure S4)**, which is not a major source of *GFP* siRNAs (**Figure S2)**. Whether these quantitative differences in levels of *GFP* splice variants and *GFP* siRNAs in two-week-old seedlings increase during development and trigger the sharp onset of PTGS in older *coi1* seedlings is not yet known.

#### Implications of pre-existing *GFP* siRNAs in the *WT T* line

Small RNA sequencing and experiments with *rdr6* and *dcl4* mutants revealed that the *WT T* line contains an endogenous and formerly unsuspected population of 21-24-nt *GFP* siRNAs that trigger a modest level of PTGS, resulting in intermediate *GFP* expression. The intermediate level of *GFP* expression in the *WT T* line can be shifted to either a GFP-weak or a hyper-GFP phenotype by mutations in factors required for PTGS (DCL4 and RDR6, respectively; this study), which change the proportions and abundance of different size classes of *GFP* siRNAs (F**igures 3 and 4)**, or by mutations in specific splicing-related factors (such as COI1, CWC16A PRP8A, PRP18A), which alter the proportions of the three *GFP* splice variants (Sasaki et al., 2015; Kanno et al., 2016, 2017a, b, 2018, 2020).

Given the new finding that the *WT T* line contains an endogenous pool of *GFP* siRNAs that can provoke a moderate degree of PTGS, it is conceivable that some of the mutants we identified in the previous screen for splicing factors (Kanno et al., 2020) could also modulate RNA silencing. Indeed, a recent study reported that PRP39 (identified in our splicing screen by its hyper-GFP phenotype; Kanno et al., 2017b), has distinct roles in not only splicing but also PTGS (Bazin et al., 2022).

#### Future perspective

An analysis of the time course of *GFP* pre-mRNA synthesis and alternative splicing patterns, as well as *GFP* siRNA accumulation, starting with young *coi1* and *WT T* seedlings and extending into later stages of development (including separation of hyper-GFP regions and silenced/recovered tissue in older *coi1* seedlings), would help to address the remaining uncertainties about *GFP* silencing/recovery in older *coi1* seedlings. Investigation of these questions in our well-defined system could reveal more general principles relevant for developmentally-associated changes in alternative splicing patterns that can potentially affect siRNA production and subsequently expression of endogenous genes.

### 2. Identifying proteins that interact with coilin

To learn more about components of coilin-dependent pathways in plants and how coilin mutations promote a hyper-GFP phenotype in young *coi1* seedlings, we performed a genetic suppressor screen using a *coi1* mutant, and conducted an immunoprecipitation-mass spectrometry (IP-MS) analysis using epitope-tagged coilin proteins.

#### Genetic suppressor screen

EMS mutagenesis and initial tests for putative suppressor mutants were performed as described in the Methods section using seeds of the hyper-GFP *coi1-8* mutant (**Figure S5**). Of the GFP-weak suppressor mutants that passed the initial tests and were subsequently carried on for further analysis (Methods), five contained a mutation in a gene retrieved in the former forward screen using the *WT T* line to identify splicing factors: *prp4k*, *sac3a*, *cbp80*, *prp8a*, and *prp18a* (**Table S1**: Kanno et al., 2020). The finding of these five genes in both the splicing screen and the *coi1-8* suppressor screen indicates that the involvement of the respective factors in *GFP* pre-mRNA splicing is independent of coilin function. However, five GFP-weak suppressor mutants contained mutations in novel factors that were identified only in the present *coi1-8* suppressor screen: *wrap53, smu2, zc3hc1*, *mad1* and *sde2-like* (**Table S1**). The failure to identify these five factors in the former splicing screen suggests their detection in our system is related in some way to the coilin deficiency in the *coi1-8* mutant. Here we discuss *wrap53*, *smu2*, and *zc3hc1*. In a *coi1-8* mutant background, these mutations reduce the intensity of GFP fluorescence and the levels of GFP protein to approximately the *WT T* intermediate state (**Figure 6 A-E, 8 DAG; Figure S6**). In addition, *coi1 zc3hc1* mutants display enhanced *GFP* silencing that extends into the shoot apex and top of hypocotyl (**Figure 6E**).

**Figure 6:**
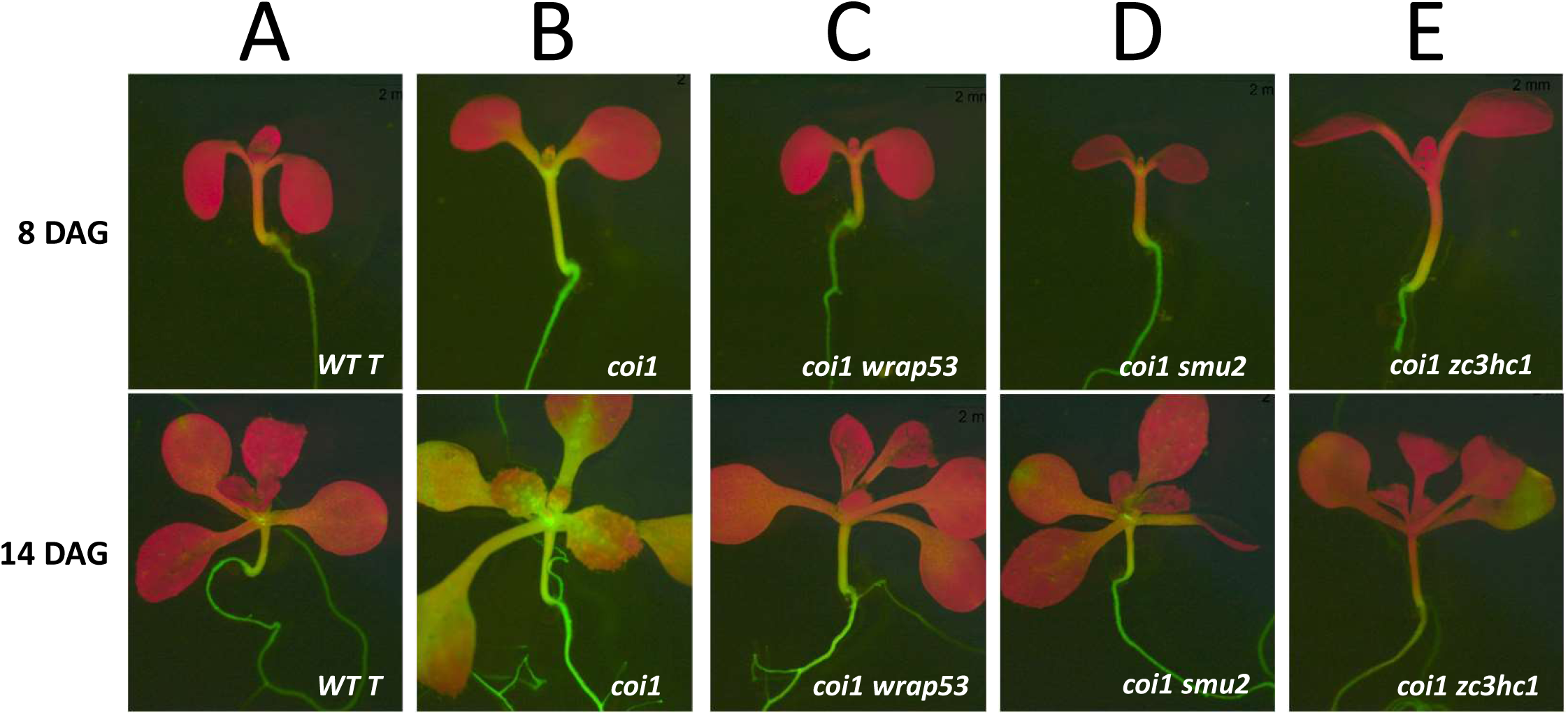
GFP phenotypes in coilin suppressor screen mutants. **(A, B)** Intermediate GFP fluorescence in the *WT T* line compared to the hyper-GFP phenotype in a non-silenced/recovered *coi1* single mutant, which was derived from the *WT T* line (Kanno et al., 2016), at 8 and 14 DAG. The *coi1* suppressor screen identified mutations that attenuate the hyper-GFP phenotype of the single *coi1* mutant. **(C-E)** Reduced *GFP* expression in the *coi1 wrap53, coi1 smu2* and *coi1 zc3hc1* double mutants at 8 and 14 DAG. The *coi1 zc3hc1* double mutant seedlings **(E)** show shortly after germination a strong reduction of GFP fluorescence that eventually extends into the shoot apex and hypocotyl. This resembles the silencing pattern observed when *dcl4* mutations are introduced into the *WT T* line, although the size class of siRNA that accumulates differs (21-nt in *coi1 zc3hc1* vs. 22-nt in *dcl4* (Figure 3C, D). Silencing observed in *coi1 zc3hc1* mutants is also distinct from the *coi1* silencing/recovery phenotype, which occurs later during seedling growth and does not spread below the site of initiation (Figure 3A). The premature and extended silencing occurs in 100% of *coi1 zc3hc1* seedlings whereas silencing/recovery occurs in only 40-90% of *coi1* seedlings. Despite the differences in the frequency, timing and pattern of *coi1 zc3hc1* silencing versus *coi1* silencing/recovery, both are associated with 21-nt *GFP* siRNAs derived from the GFP CDS (Figure 3A, D).

##### *WRAP53* (AT4g21520)

WRAP53/TCAB1 (WD40 encoding antisense to P53/telomerase Cajal body protein 1) is an evolutionarily conserved WD40 repeat protein that is widely distributed in the plant, fungal and animal kingdoms (**Figure 7A**; **Figure S7A**). To our knowledge, WRAP53 has not yet been investigated in detail in plants nor have *wrap53* mutations been recovered in any prior genetic screen.

**Figure 7:**
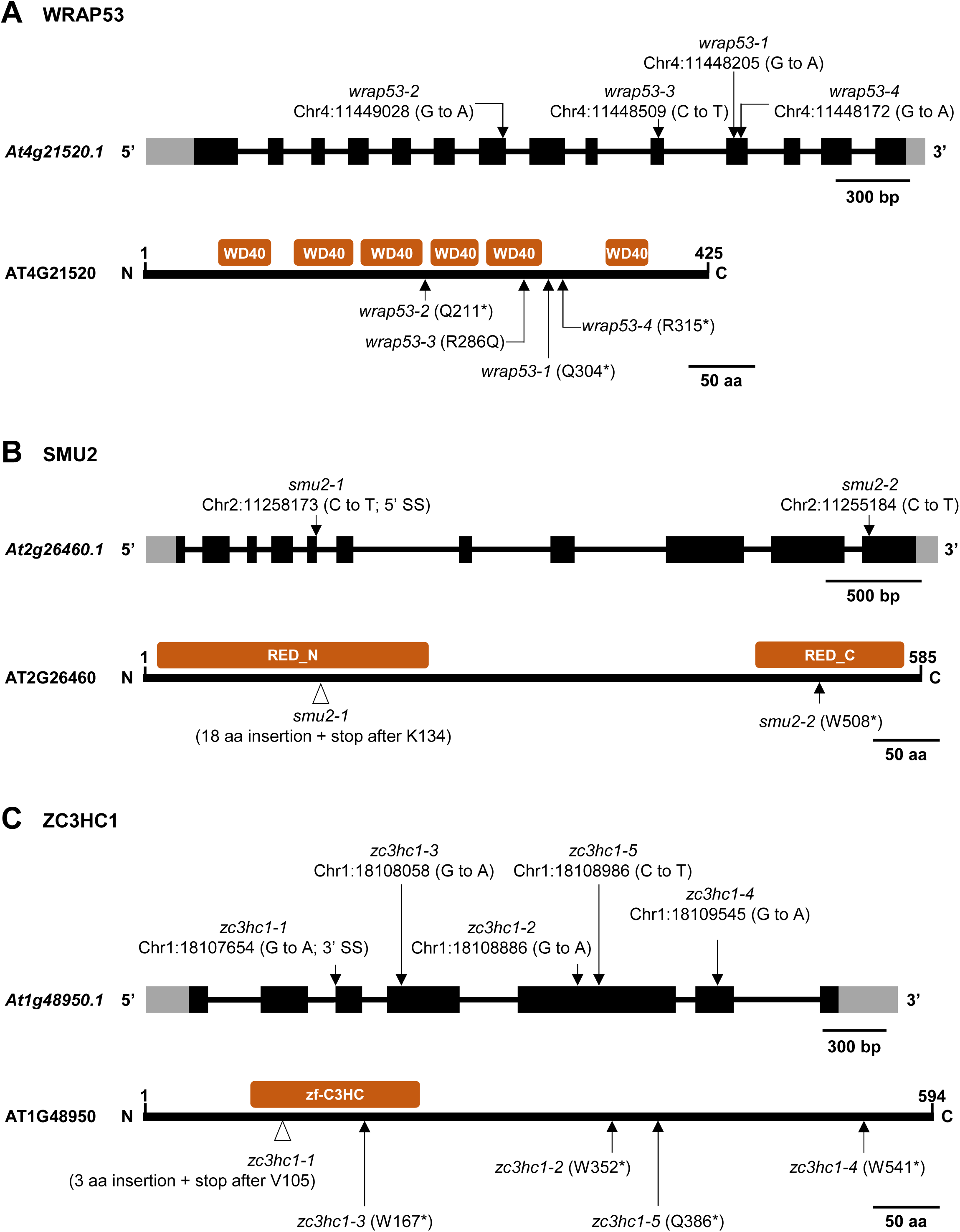
*WRAP53, SMU2* and *ZC3HC1* gene structures, protein domains and positions of mutations. Intron-exon structures of the indicated genes and nucleotide sequence changes are shown at the top of each section; changes in amino acid sequences at the bottom (asterisks indicate premature termination codons). Recognizable protein domains are indicated by red boxes. Amino acid sequence alignments of ATWRAP53, ATSMU2 and ATZC3HC1 proteins with the corresponding orthologs in other plant species and model organisms are shown in **Figure S7A-C**. **(A)** ATWRAP53 (At2g21520) is 425 amino acids in length and encoded by a single copy gene in Arabidopsis. ATWRAP53 protein contains six WD40 repeat domains. In our screen, we recovered four *wrap53* alleles, all of which contain a mutation in or around one of the six WD40 domains. **(B)** ATSMU2 (At2g26460) is 585 amino acids in length and encoded by a single copy gene in Arabidopsis. In metazoans, SMU2 is referred to as RED, named after a region rich in Arg (R)/Glu (E) or Asp/ Asp (D) repeats, which can also be identified in ATSMU2 close to the N- and C-termini. Our screen recovered two *smu2* alleles, both of which contain mutations that are located in a recognizable RED domain. **(C)** ATZC3HC1 (At1g48950) is a zinc finger protein that is 594 amino acids in length and encoded by a single copy gene in Arabidopsis. ATZC3HC1 contains one zf-C3Hc domain. Our screen recovered five independent *zc3hc1* alleles.

In mammalian cells, WRAP53 has been shown to interact with coilin (Machyna et al., 2015) and to be essential for CB cohesion (Henriksson and Farnebo, 2015). By contrast, we found that a *wrap53* mutation - unlike *coi1* mutations - does not disrupt CB structure in Arabidopsis, at least in the cell type tested (trichomes) (**Figure S8**). Furthermore, we did not identify ATWRAP53 as a coilin-interacting protein in a comprehensive IP-MS analysis (**Table S2**). These results suggest that WRAP 53 may act differently in plants than in mammals and not partner directly with coilin to support CB structure. Additional work in the future is needed to confirm this conjecture and to determine the mechanism by which *wrap53* mutations attenuate the hyper-GFP phenotype in young *coi1 wrap53* seedlings.

##### SMU2 (*suppressors of mec8 and unc52*) (At2g26460)

SMU2 is a splicing factor that is conserved in most plant and animal species that carry out alternative splicing (**Figure 7B**; **Figure S7B**). Previous studies in plants found that SMU2 is important for constitutive and alternative splicing in maize (Chung et al., 2009) and for magnesium homeostasis in Arabidopsis (Feng et al., 2020).

In mammalian cells, the SMU2 ortholog, termed RED (**Figure 7B**), interacts with the protein SMU1 to form a complex required for spliceosome activation (Keiper et al., 2019). Although interactions between SMU1-SMU2 proteins have been reported in plants (Chung et al., 2009; Feng et al, 2020), we were unable to demonstrate such an interaction using yeast two-hybrid assays, possibly because the SMU1 protein was unstable in yeast cells under our experimental conditions. Interestingly, however, we did retrieve *smu1* mutants in the previous screen using the *GFP* splicing reporter to identify splicing factors (Kanno et al, 2017a, 2020). Whereas a *smu2* mutant (in a *coi1* background) exhibits a GFP-weak phenotype resembling that of the *WT T* line (**Figure 6D**), a *smu1* mutant (in a wild-type *COI1* background) displays a hyper-GFP phenotype (Kanno et al, 2017a). The opposing effects of *smu1* and *smu2* mutations on the same *GFP* splicing reporter gene probably reflect differing impacts on splicing of the two mutations in specific genetic backgrounds. In *coi1 smu2*, the unspliced, untranslatable *GFP* pre-mRNA predominates (**Figure S3**), whereas in *COI1 smu1*, the translatable AU-AG transcript is the major splice variant (Kanno et al., 2017a). (The impact of the *coi1* mutation on SMU1 activity and the activity of the *smu2* mutation in a *COI* background have not yet been assessed.) A further interesting but currently unexplained feature of the splicing defects in *coi1 smu2* mutants is the exceptionally high number of intron retention (IR) events, and to a lesser extent exon skipping (ES) and alternative 5’-splice site events compared to *coi1 wrap53* and *coi1 zc3hc1* mutants (**Table S4**).

##### *ZC3HC1* (At1G48950)

ZC3HC1 is a zinc finger protein containing a zf-C3HC domain (**Figure 7C**). Although this domain is distributed throughout the eukaryotic kingdom, the *ZC3HC1* gene is specific to plants (**Figure S7C**). We identified five *zc3hc1* alleles in our screen (**Figure 7C**), all of which confer a typical GFP-weak phenotype in a *coi1* mutant background (**Figure 6E, 8 DAG**). Around one to two weeks after germination, however, the initially GFP-weak phenotype *coi1 zc3hc1* mutants intensifies and spreads downward into the hypocotyl (**Figure 6E**). Although this silencing pattern is reminiscent of that seen in *dcl4* backgrounds (**compare Figure 3D with Figures 3C and 4C**), the predominant size class of *GFP* siRNA involved in silencing differs in each case: 22-nt in the case of *dcl4* and 21-nt in the case of *coi1 zc3hc1* mutants (**Figure 3C; Figure S2**). The increased level of 21-nt siRNAs in *coi1 zc3hc1* relative to a *coi1* single mutant and other *coi1* suppressor mutants (**Figure S2**) hints that the wild-type ZC3HC1 protein may promote biogenesis and/or accumulation of *GFP* 21-nt siRNAs.

During the course of our study, ZC3HC1 was also identified as MEM1 (*methylation elevated mutant1*) in Arabidopsis using a reverse genetics approach to identify new proteins involved in DNA demethylation (Lu et al., 2020). MEM1 was found to act similarly to ROS3 in a ROS1-mediated DNA demethylation pathway to remove methylation from transposons and transgene promoters (Lu et al., 2020). MEM1 has also been reported to safeguard against DNA damage (Wang et al., 2022). In our system, we did not detect changes in DNA methylation in the upstream enhancer of the *GFP* reporter gene in a *coi1 zc3hc1* mutant (**Table S3; Figure 1**). Therefore, the *zc3hc1* mutations we identified are not likely to reduce *GFP* expression in our system through a mechanism that involves DNA methylation/demethylation of transcriptional regulatory regions. Conceivably, MEM1 could play roles in multiple processes including PTGS, DNA demethylation, and DNA repair depending on the physiological context in which it acts.

#### Immunoprecipitation-mass spectrometry analysis (IP-MS)

To identify putative coilin interacting proteins, we performed an IP-MS analysis using three epitope-tagged coilin proteins: C-terminal mRED (coilin-mRed); N-terminal FLAG (FLAG-coilin); and C-terminal FLAG (coilin-FLAG) (**Table S2**). This analysis identified several major functional categories of putative coilin-interacting proteins: (1) splicing factors, including many Sm proteins that constitute the SMN complex needed for biogenesis of spliceosomal snRNPs (Hebert et al., 2001: Xu et al., 2005); (2) histone proteins and histone acetylases/deacetylases, many of which are located in the nucleolus; and (3) nucleolar proteins involved primarily in rRNA processing or modification (**Table S2**). These results are consistent with prominent roles for plant coilin in splicing, nucleolar chromatin structure, and ribosome function (Machyna et al, 2015). Moreover, they are compatible with the observation that coilin is present in both CBs and nucleoli (Trinkle-Mulcahy and Sleeman, RNA, 2017).

## CONCLUSIONS

In this study, we used a *GFP* splicing reporter gene in a *coi1* mutant background to carry out experiments designed to investigate an unusual gene silencing phenomenon appearing in older *coi1* seedlings, and to learn more about coilin functions and components of coilin pathways in plants. Our results indicate that *GFP* silencing/recovery in older *coi1* seedlings follows a canonical PTGS pathway that requires 21-22-nt siRNAs emanating largely from an aberrant splice variant of *GFP* pre-mRNA. The extent to which *coi1* mutations enhance silencing/recovery is not yet known. So far, *coi1* is the only hyper-GFP mutant we have identified (Kanno et al., 2020) that frequently displays such a striking silencing/recovery phenotype, suggesting that the *coi1* mutation sensitizes the *GFP* reporter gene to PTGS. We also uncovered the existence of a previously unknown population of endogenous *GFP* siRNAs in the *WT T* line. These siRNAs are able to induce a moderate level of PTGS leading to intermediate *GFP* fluorescence midway between GFP-weak and hyper-GFP phenotypes. The trans-generational stability of intermediate GFP fluorescence in the *WT T* line renders it a valuable tool for identifying mutations that either enhance or suppress GFP expression, and are thus likely to impair factors acting in alternative splicing, RNA silencing and, in a *coi1* mutant background, general coilin functions in plants.

In addition to TGS/RdDM (Kanno et al., 2008; Eun et al., 2012) and alternative splicing (Kanno et al., 2020), the new findings of PTGS involvement add a third dimension to processes that can influence the activity of the *GFP* splicing reporter. Our detailed analysis of this system has revealed its versatility and usefulness for studying the contributions of different expression mechanisms and their coordinated influence on the activity of a complex genetic locus.

The *coi1* suppressor screen identified several new proteins that may functionally interact with coilin. We provide foundational information for three of these proteins - WRAP53, SMU2 and ZC3HC1 – all of which remain to be fully characterized in plants. Future study of these proteins is not only of inherent interest but is also potentially valuable for discovering expanded functions of coilin in plants. Seeds of the *WT T* line and the mutants as well as the high-throughput sequencing data we generated are publicly available and provide many resources for the plant scientific community (**Table S1**).

An IP-MS analysis substantiated major roles for plant coilin in splicing, nucleolar chromatin structure and rRNA processing. These processes require in some cases small RNAs. (Wang et al., 2015; Kishore et al., 2010; Park et al., 2019; Wang and Chekanova, 2016; Preuss et al., 2008; Huang et al., 2022). The identification of *zc3hc1* mutations in the *coi1* suppressor screen and their enhancement of *GFP* siRNA accumulation provide at least an indirect link between coilin function and siRNA biogenesis and/or stability.

Our cumulative results highlight the involvement of coilin in multiple processes, some of which can be unified through a requirement for small RNAs. These findings lend support to previous proposals suggesting that coilin may facilitate RNP biogenesis by acting as a chaperone for small nuclear noncoding RNAs (Machyna et al., 2015). Regarding coilin’s participation in alternative splicing, our prior forward genetic screen (Kanno et al., 2020) represents, to our knowledge, the first in any organism based on an alternatively-spliced gene and it is the only forward screen, apart from ‘*no cajal bodies’* (*ncb*), which identified *coi1* mutants (Collier et al., 2006). These findings suggest that coilin protein may carry out a special role in specifically alternative splicing in a manner that is worthy of further study.

## DATA AVAILABILITY

Seed stocks and nucleic acid sequences discussed in this paper are available under the Accession Numbers listed in Table S1.

## ACKNOWLEDGEMENTS

We are grateful to the following collegues: Shou-Jen Chou and the Genomic Technology Core facility at IPMB for preparation of DNA, RNA and sRNA libraries and DNA and RNA sequencing; Chuan-Chih Hsu and Tuan-Nan Wen for assistance in analyzing and processing LC-MS/MS data: Chia-Liang Chang for technical assistance; and Maria Kalyna, Hervé Vaucheret, Olivier Voinnet, Lucia Clemens-Daxinger and Florian Mette for helpful discussions.

## Legends for Supplementary information

### Supplementary Figures

**Figure S1:**
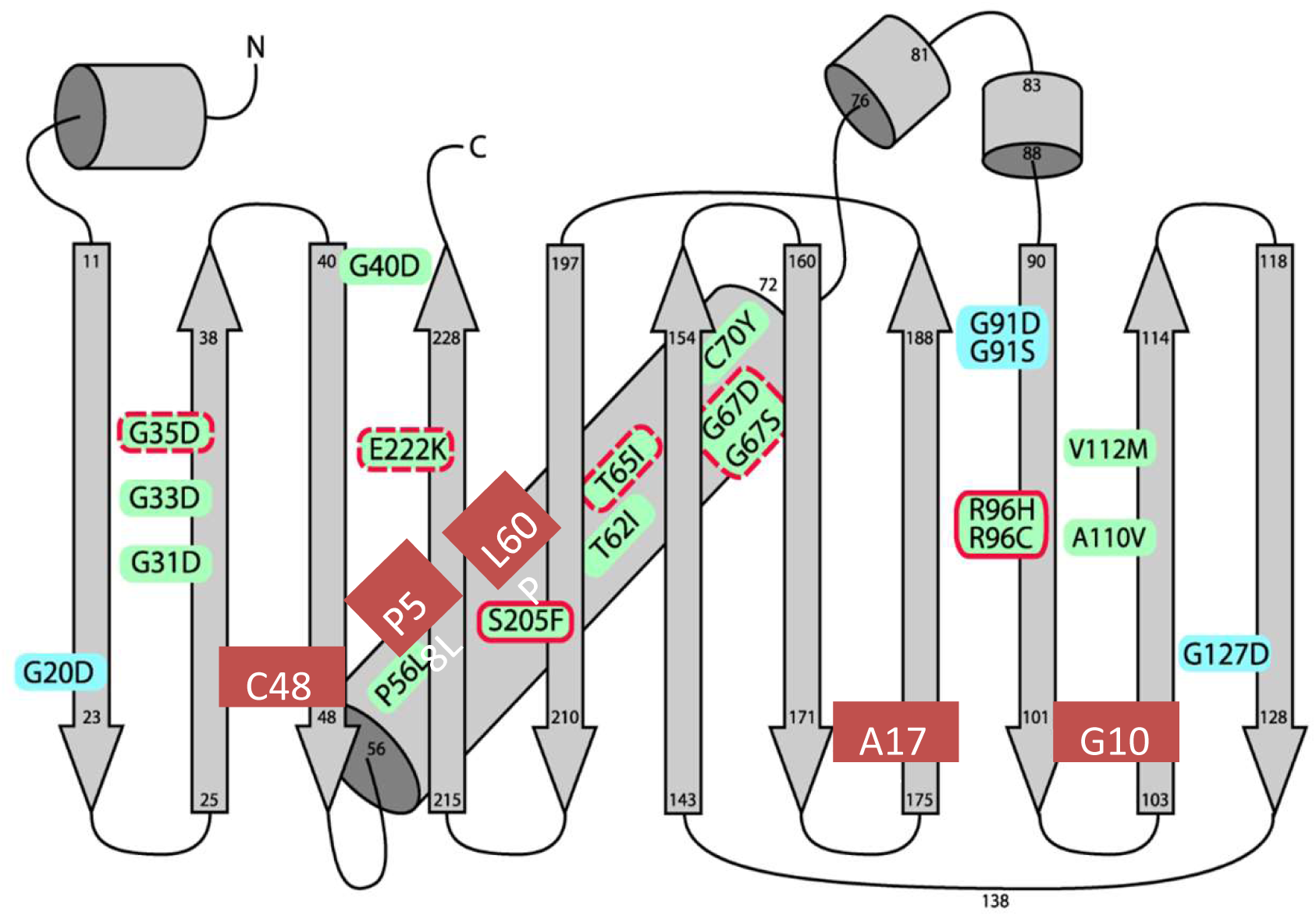
Overall GFP protein fold and positions of amino acid substitutions leading to loss of fluorescence. Figure and text are adapted from Figure 1 in Fu et al. (2015). The figure shows a schematic depiction of the overall fold of Thr65-GFP protein, the GFP variant that was used in the T-DNA construct (Figure 1 in Fu et al. 2015). Vertical arrows indicate the eleven β strands of the β-barrel structure. Amino acid residue numbers at the base and tips of the arrows denote the beginning and ends of secondary structural elements. The chromogenic tripeptide (Thr65-Tyr66-Gly67) is located on an internal α-helix (diagonal cylinder) extending from amino acids 56 to 72. Amino acid substitutions identified in a previous screen (Fu et al., 2015) that lead to losses of fluorescence are shown in blue, gray or green backgrounds. Solid red outlines indicate substitutions causing defects in chromophore formation without substantial reductions in GFP protein accumulation (Fu et al., 2015). Dotted red outlines designate substitutions resulting in lowered levels of GFP protein accumulation relative to wild-type. For the remaining substitutions, no GFP protein was detected by Western blotting (Fu et al., 2015). Lid residues at the N and C termini (G91 and G127) and the opposite side (G20), which is referred to as the ‘top’ of the barrel (Zimmer et al., 2014), are highlighted in blue. Noteworthy new mutations identified in the current study (**Table S1**) are shown on red backgrounds. These include G140D, which is present at a hinge region of the GFP secondary structure and thus is likely to be required for GFP folding and stability. Accordingly, G140 is one of 23 highly conserved amino acids in GFP-related proteins (Ong et al, 2011; Fu et al., 2015). A second notable mutation is C58L, which is present in stretch of several proline residues, including C56 that was identified in a previous screen (Fu et al., 2015). These cysteine residues are thought to be important for maintaining the alpha-helical structure necessary for chromophore formation (Fu et al. 2015). The remaining new mutations - L60P. A179T and C48Y - lead to loss of fluorescence by unknown means. The finding of the these new *gfp* mutations add to the growing list of *gfp* loss-of-function mutations, which are proving valuable in various studies for determining amino acid residues important for GFP fluorescence and protein stability, and for using GFP protein as a sensor (Pauly et al., 2017; Birnbaum et al., 2020; Decaestecker et al., 2019).

**Figure S2:**
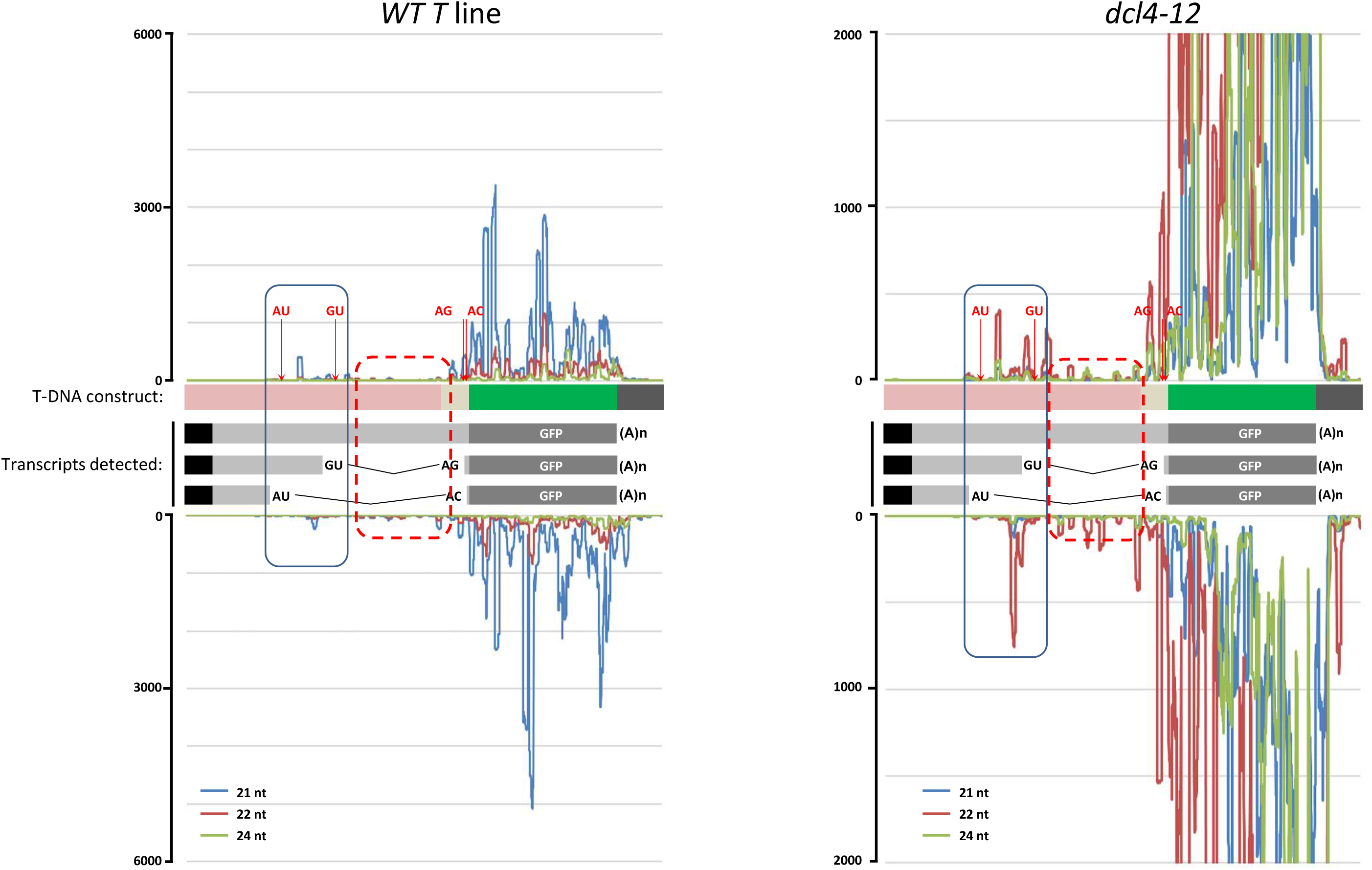
Probable source of pre-existing *GFP* siRNAs in the *WT T* line. We argue here that most of the *GFP* siRNAs in the *WT T* line and its mutant derivatives, which contain varying levels of all three alternatively spliced *GFP* transcripts (**Figure S4**), may be originating mainly from the spliced, untranslatable GU-AG transcript, with negligible contributions from the unspliced *GFP* pre-mRNA and the spliced, translatable AU-AC variant (*GFP* mRNA) (Figure 1). Evidence for this claim can be discerned by inspection of the siRNA distribution profile along the GFP protein coding sequence and upstream region. To illustrate the point, such results are shown here for the *WT T* line and a *dcl4-12* mutant. Blue, red and green vertical lines represent 21-, 22- and 24-nt siRNAs, respectively. The *GFP* coding region is indicated by the dark gray bars and the upstream region by light gray bars. Positions of splice sites in this region are indicated (also shown in Figure 1). Spliced introns are represented by thin black lines. *Negligible contributions to GFP siRNAs from the translatable AU-AC transcript and unspliced pre-mRNA* Although *GFP* siRNAs are most abundant in the GFP protein coding region, the translatable AU-AC transcript (*GFP* mRNA) does not appear to contribute significantly to the *GFP* siRNA pool. The best evidence for claim is provided by the *cwc16a* single mutant, which accumulates primarily the translatable AU-AC transcript and only minor amounts of the two untranslatable transcripts, and does not produce siRNAs (**Figure S3A and B; Figure 5A**) (Kanno et al., 2017a, 2020). The unspliced *GFP* pre-mRNA can also be ruled out as a prominent source of siRNAs as evidenced by the near absence of siRNAs from its central region (**red dotted box;** *WT T* and *dcl4-12*), which is spliced out in GU-AG and AU-AC transcripts. This demonstrates that the unspliced variant, which accumulates to an appreciable level in all genotypes (**Figure S4**), is not a major source of siRNAs. *Convincing contributions from the GU-AG splice variant to the GFP siRNA pool* The presence of siRNAs derived from the extended first exon of the untranslatable GU-AG transcript **(solid blue boxes; *WT T* and *dcl4-12***), which is not present in the translatable AU-AC transcript, supports the idea that the GU-AG transcript provides the primary substrate for RDR6 synthesis of dsRNA and subsequent processing to siRNAs by DCL activities. The accumulation of siRNAs from the unique first exon of the untranslatable GU-AU transcript can be seen most clearly in a *dcl4* mutant (*dcl4-12*; **blue box**), which produces large amounts of primarily 22-nt *GFP* siRNAs owing to strong activity of DCL2 in the absence of DCL4. The lower levels of siRNAs from the unique first exon of the GU-AU transcript compared to the *GFP* coding region probably reflect the reduced RDR6 activity as it progresses 3’ to 5’ along the single stranded aberrant RNA substrate. RDR6 has been reported to read at least 750 nt, with enzyme activity decreasing progressively toward the 5‘end of the substrate RNA (Moissiard et al., 2007). In our T-DNA construct, the enhancer region extends for 1277 bp upstream of the minimal 35S promoter (91bp) and GFP coding region (720 bp) (Figure 1). Hence the unique exon giving rise to siRNAs from the GU-AU transcript is well beyond the optimal range of RDR6, which likely results in less double stranded (ds) RNA from this region and a lower quantity of siRNAs. We assume the *GFP* 21-nt siRNAs (and the 22-nt siRNAs in a *dcl4* mutant background) represent primary siRNAs derived from cleavage of RDR6-dependent synthesis of dsRNAs using the GU-AG transcript as a substrate, although we cannot rule out more complicated scenarios.

**Figure S3:**
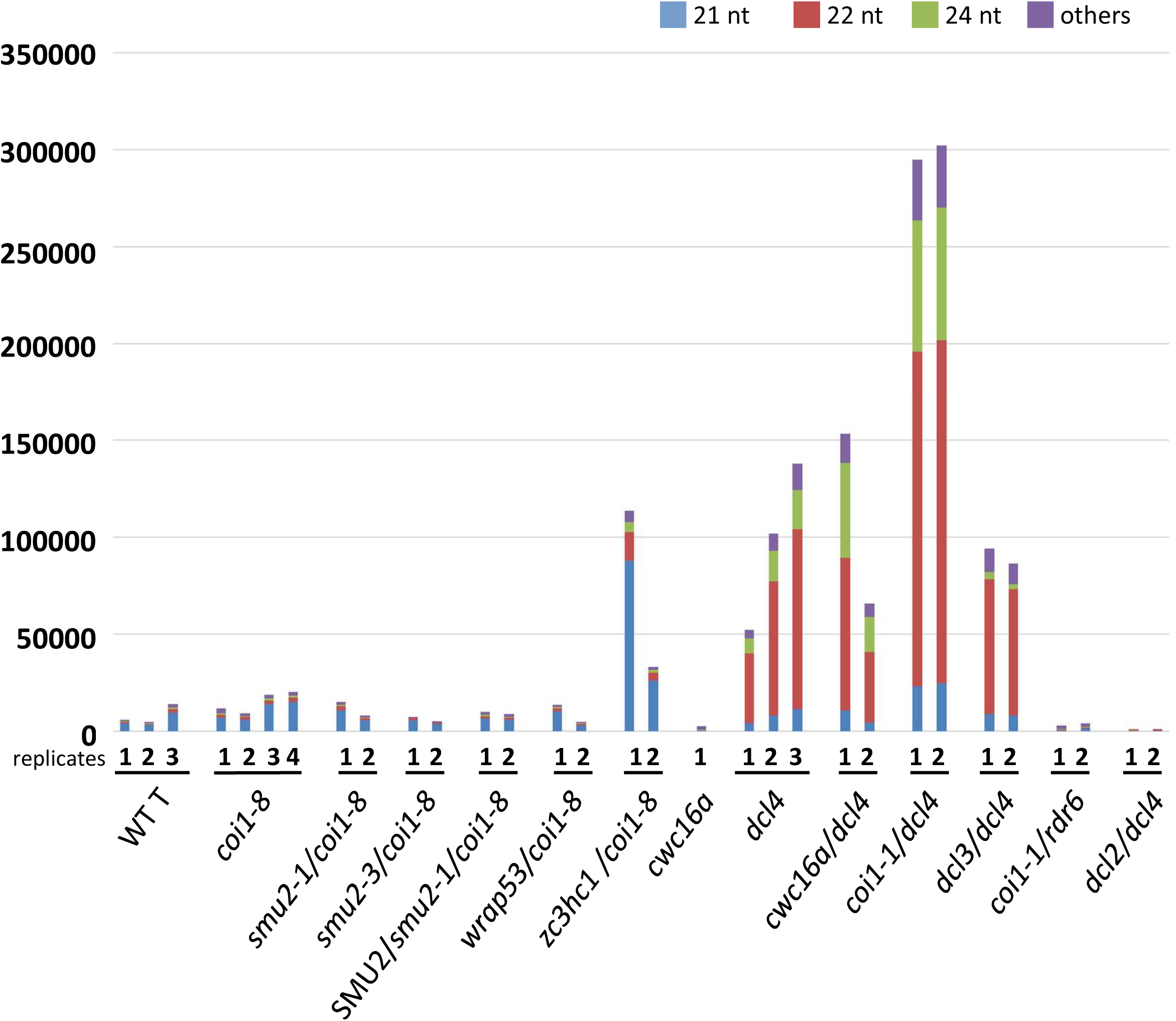
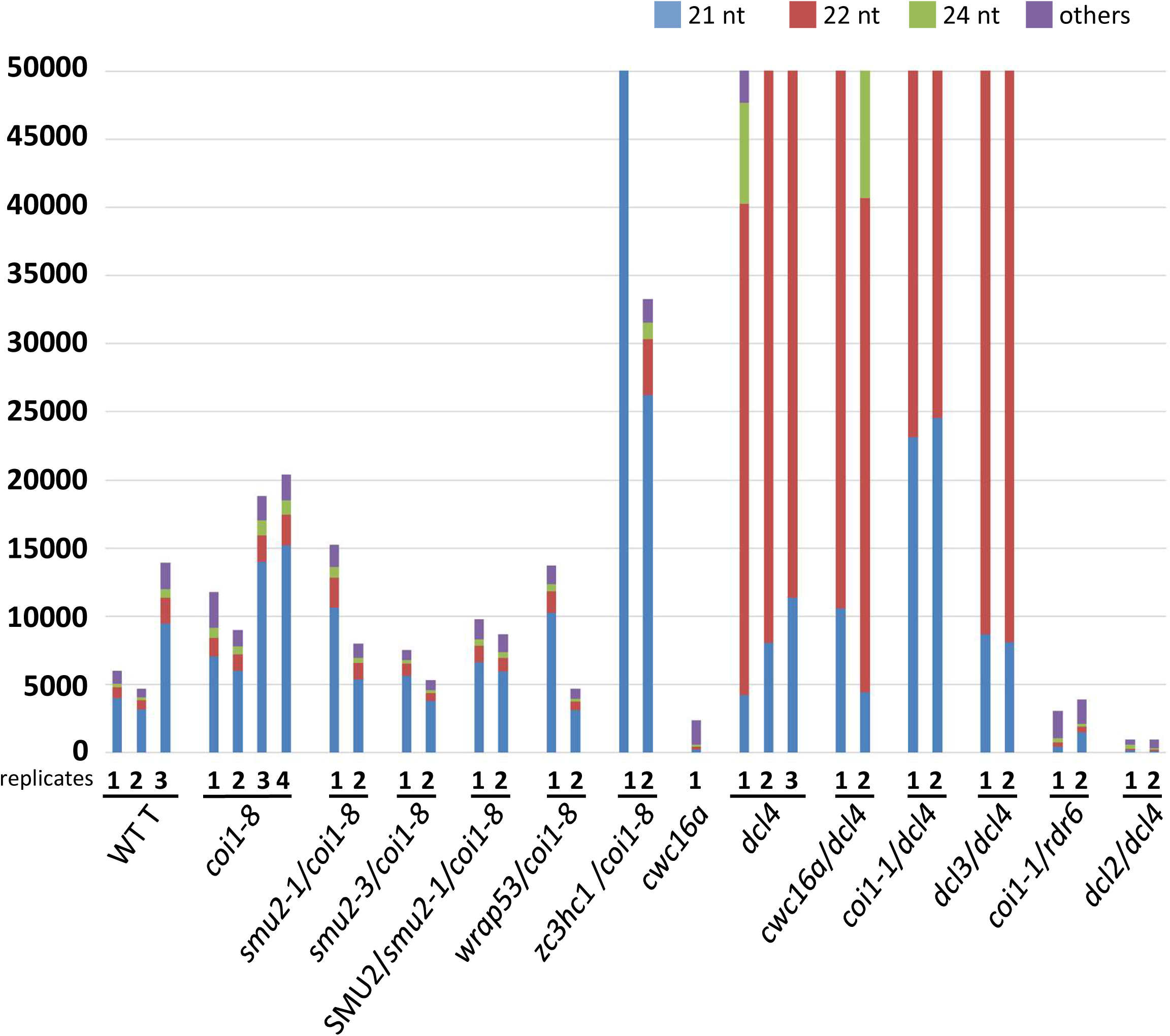
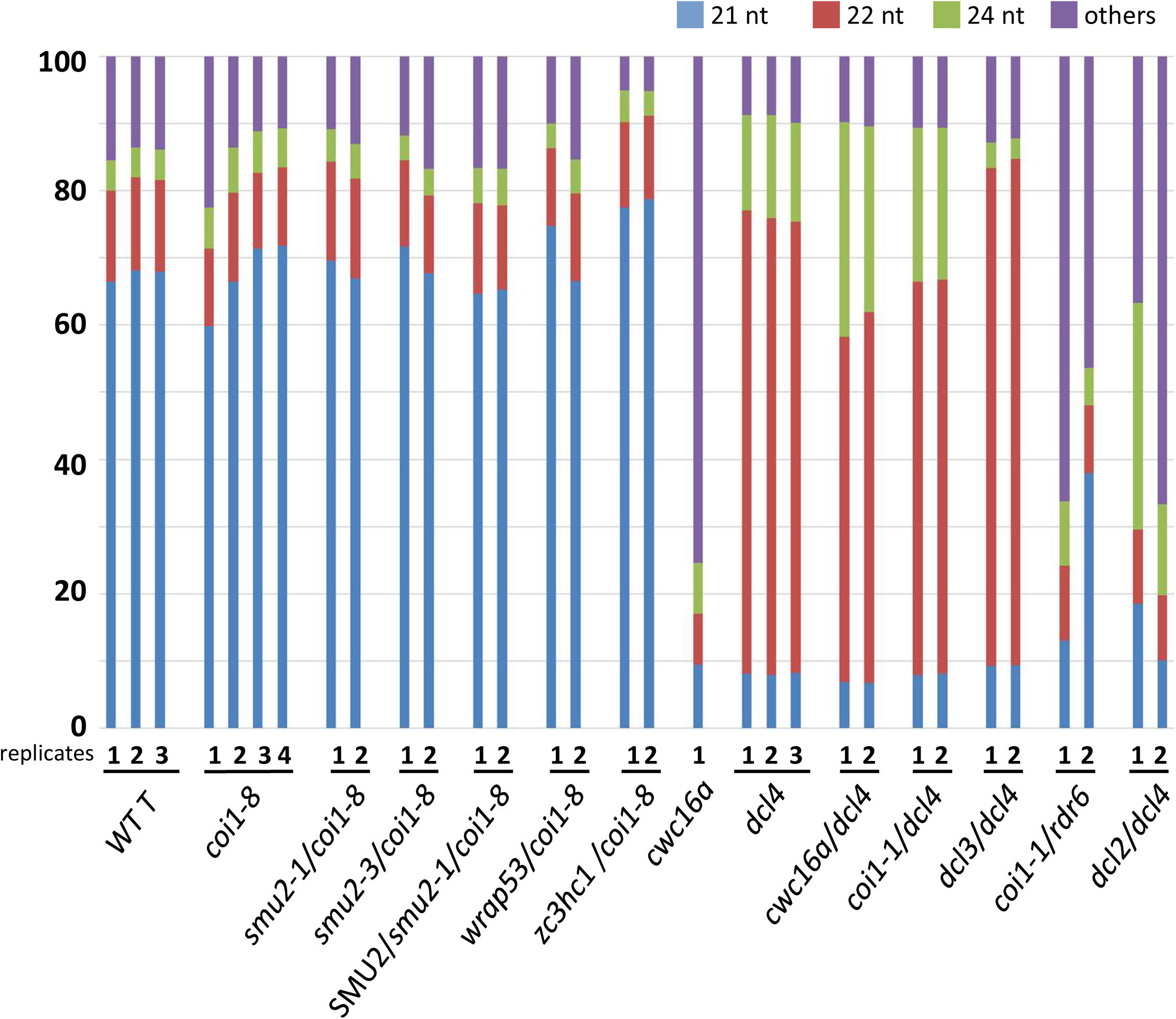
Comparison of abundances and size classes of *GFP* siRNAs in coilin suppressor screen mutants and other mutants used in this study. The y-axis shows the number of reads of *GFP* siRNAs in the indicated mutants. Blue, red and green vertical lines represent 21, 22, and 24-nt siRNAs. respectively. The number of ‘replicates’ along the x-axis refers to the number of biological replicates. **Parts A** and **B** show the same read data on different scales of maximum reads. The y-axis in **Part C** shows the ratio (percentage) of each size class of siRNA in each mutant. The suppressor mutants are all homozygous for the indicated suppressor mutation (*smu2-1*, *wrap53-1*, *zc3hc1*) and the *coi1-8* mutation.

**Figure S4:**
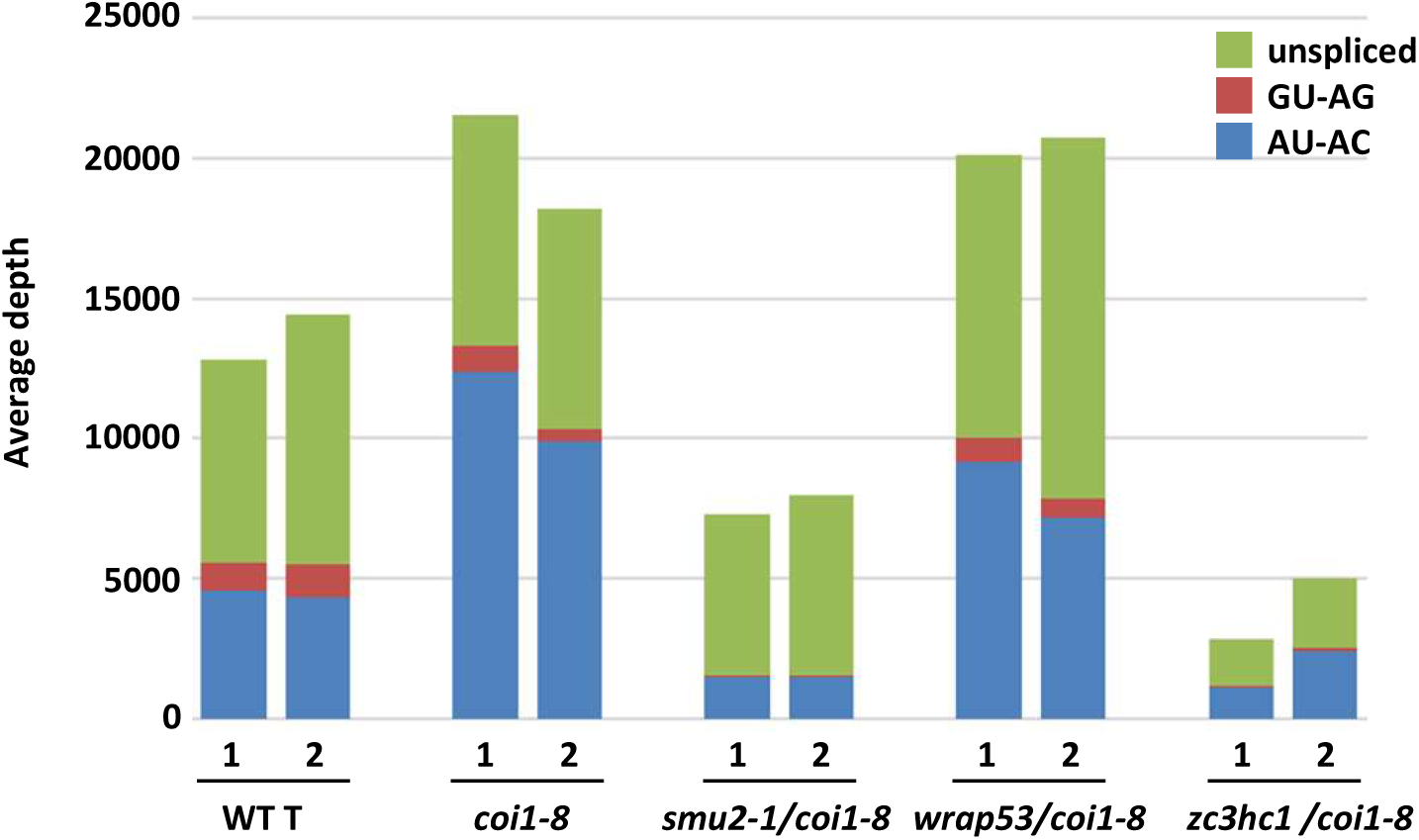
Proportions of three *GFP* splice variants in *coi1-8* and three coilin suppressor mutants. The y-axis shows the average depth (inferred from the average number of reads covering different regions involving different combinations of *GFP* splice variants) in each of the *GFP* splice variants in the indicated mutants. The x-axis shows the number of biological replicates. Blue bars: AU-AG transcript (spliced, *GFP* mRNA): red bars (GU-AG spliced, untranslatable variant, proposed main source of *GFP* siRNAs); green bars (unspliced *GFP* pre-mRNA). The suppressor mutants are all homozygous for the indicated suppressor mutation (*smu2-1*, *wrap53-1*, *zc3hc1*) and the *coi1-8* mutation.

**Figure S5:**
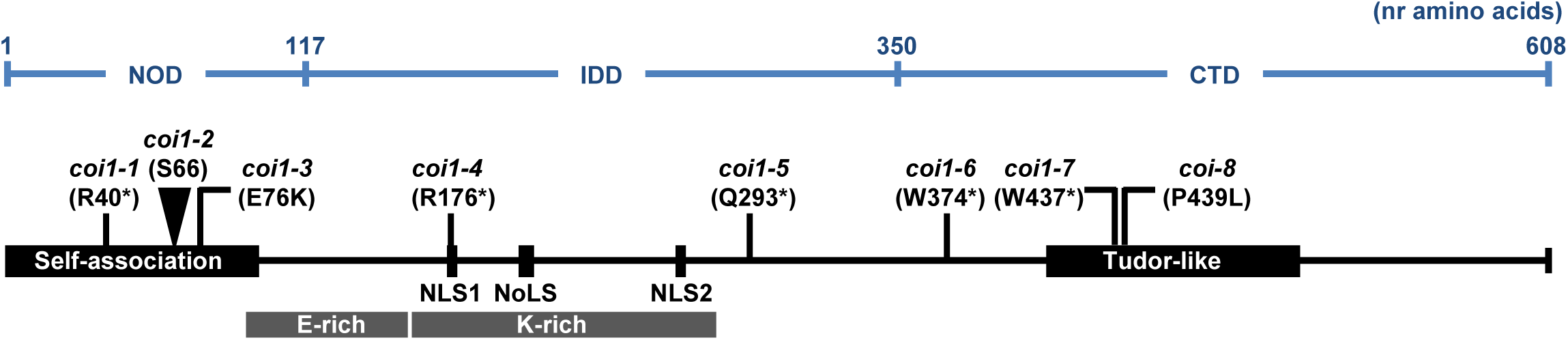
Coilin domains and positions of amino acid changes in *coi1* mutants. Figure and text are modified from Figure 2 in Kanno et al. (2016). Recognizable domains and motifs in coilin (At1G13030) include a self-association domain at the N-terminus; an atypical tudor domain at the C-terminus; two nuclear localization signals (NLS); and a nucleolar localization signal NoLS). Analysis of the secondary structure of Arabidopsis coilin revealed three structural domains: NOD, N-terminal globular domain; IDD, internal disordered domain; CTD, C-terminal domain (Makarov et al., 2013). The indicated *coi1* mutations were originally identified as <u>h</u>yper-<u>GF</u>P (*hgf*) mutants in a previous forward screen for factors affecting alternative splicing and expression of the *GFP* reporter gene (Kanno et al., 2016). The *coi1-8* mutant (P439L) was used for the present coilin suppressor screen.

**Figure S6:**
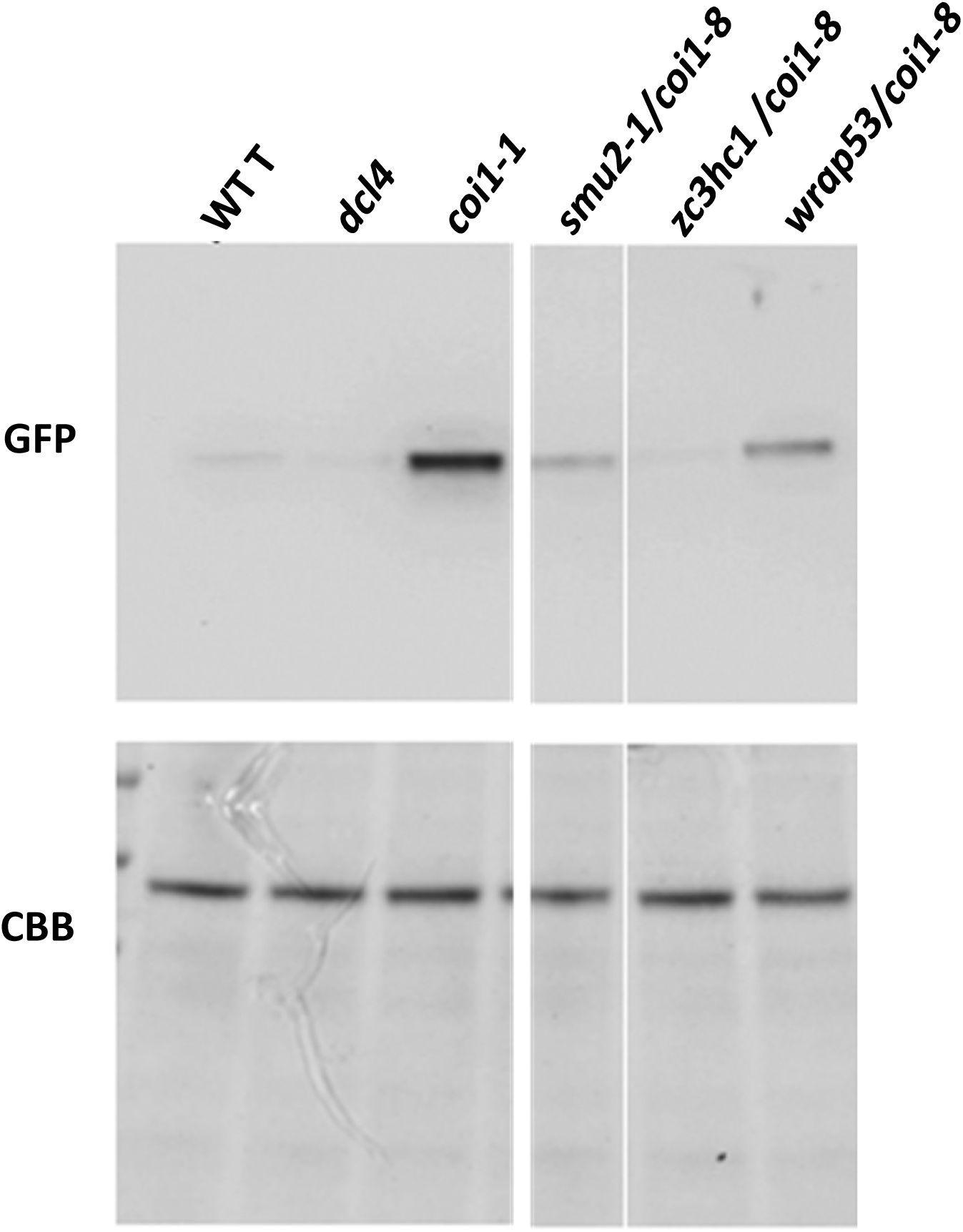
Reduced accumulation of GFP protein in coilin suppressor mutants. Western blots were used to detect GFP protein levels in the GFP-weak coilin suppressor mutants (which are homozygous for the *coi1-8* mutation) compared to the *WT T* line and the *coi1* hyper-GFP single mutant. Proteins were isolated from two-week-old seedlings with the indicated genotypes and further processed for Western blotting as described in previous publications (Fu et al., 2015; Kanno et al., 2020). The Western blot was probed with monoclonal antibodies to GFP (Roche, catalog nr. 11814 460001). For a loading control, a duplicate gel containing the same samples was run and stained with Coomassie Brilliant Blue (CBB). The coilin suppressor mutants all contain less GFP protein than the single *coi1-8* mutant and approach the level observed in *WT T* seedlings.

**Figure S7:**
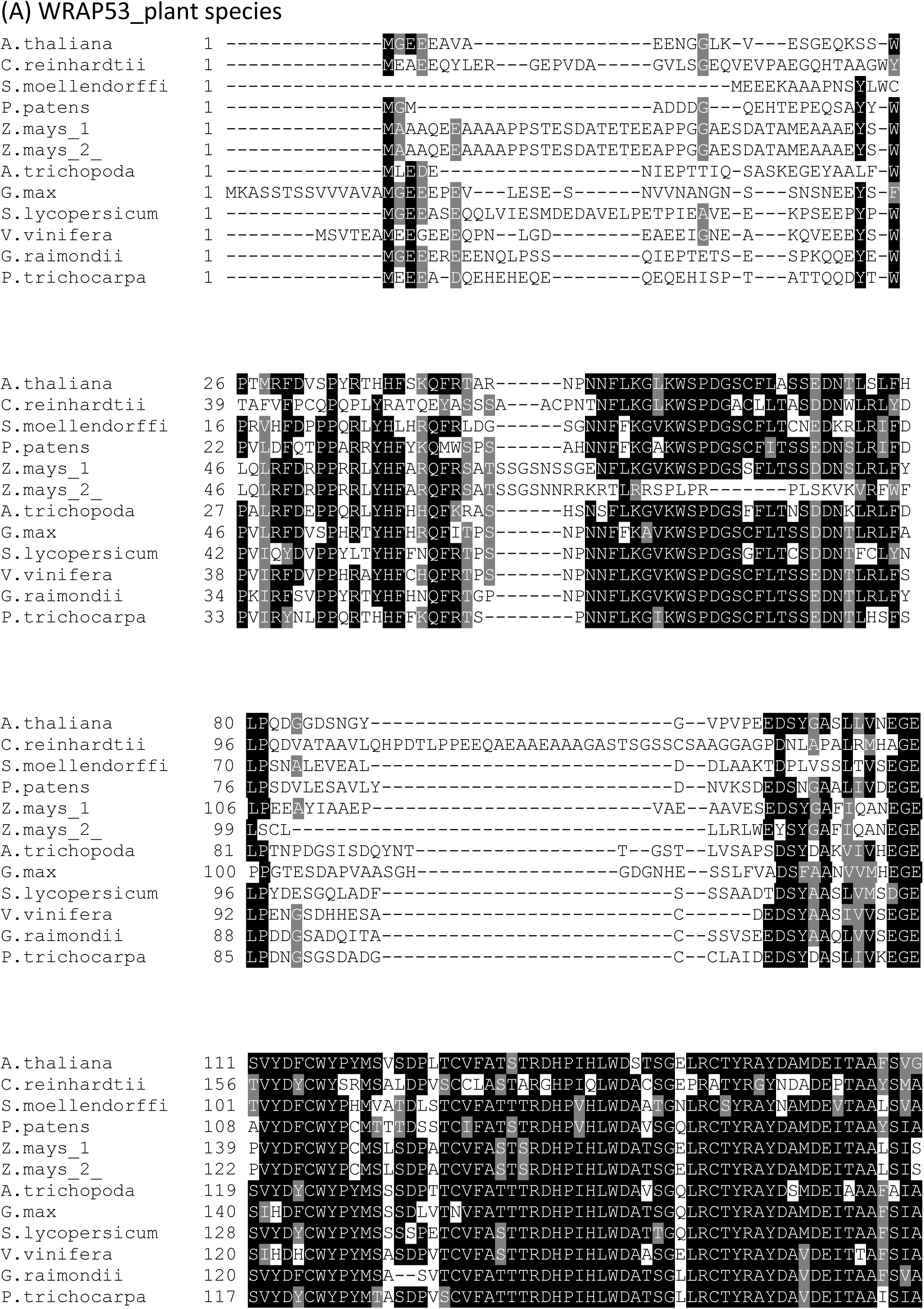
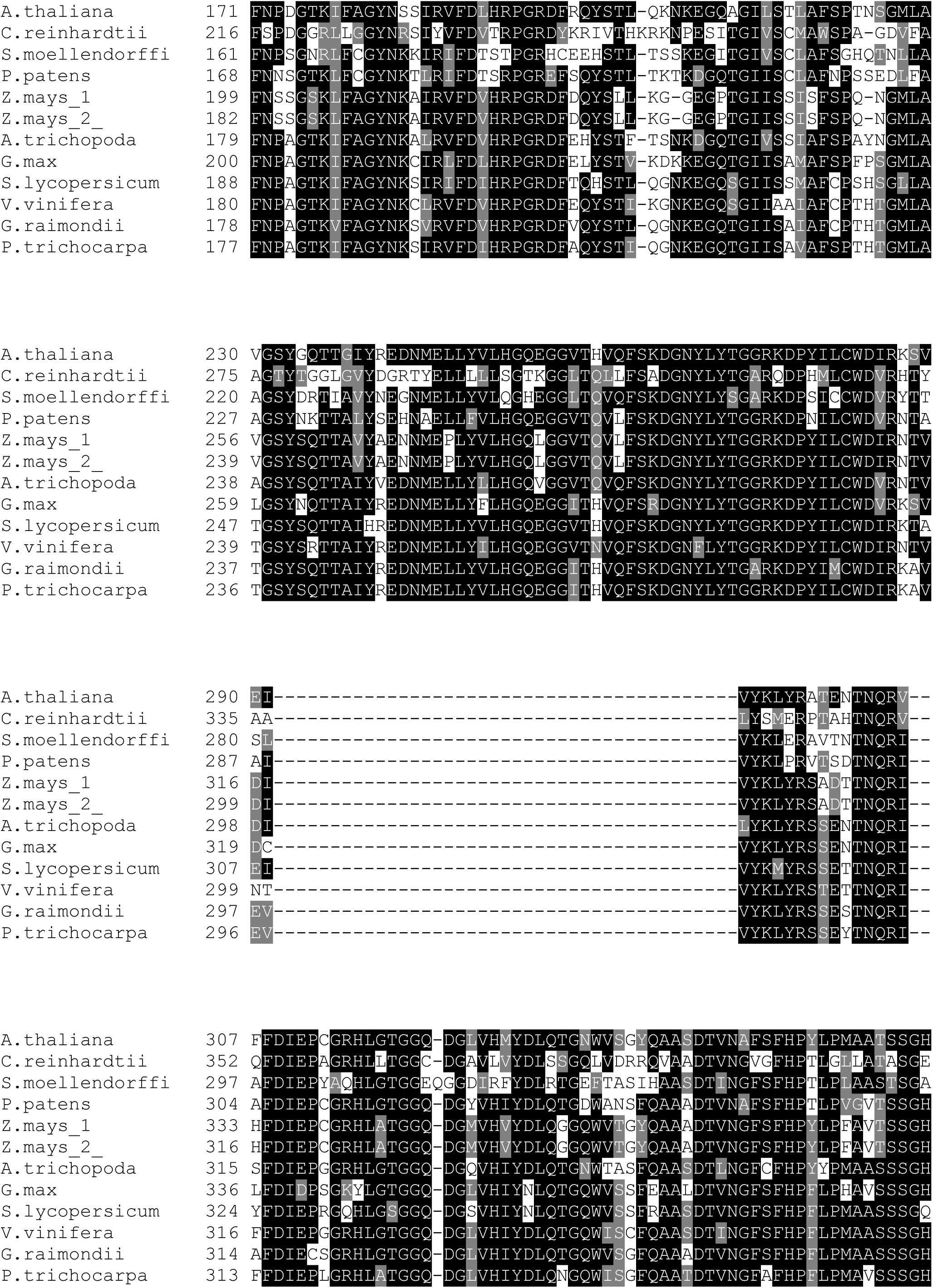
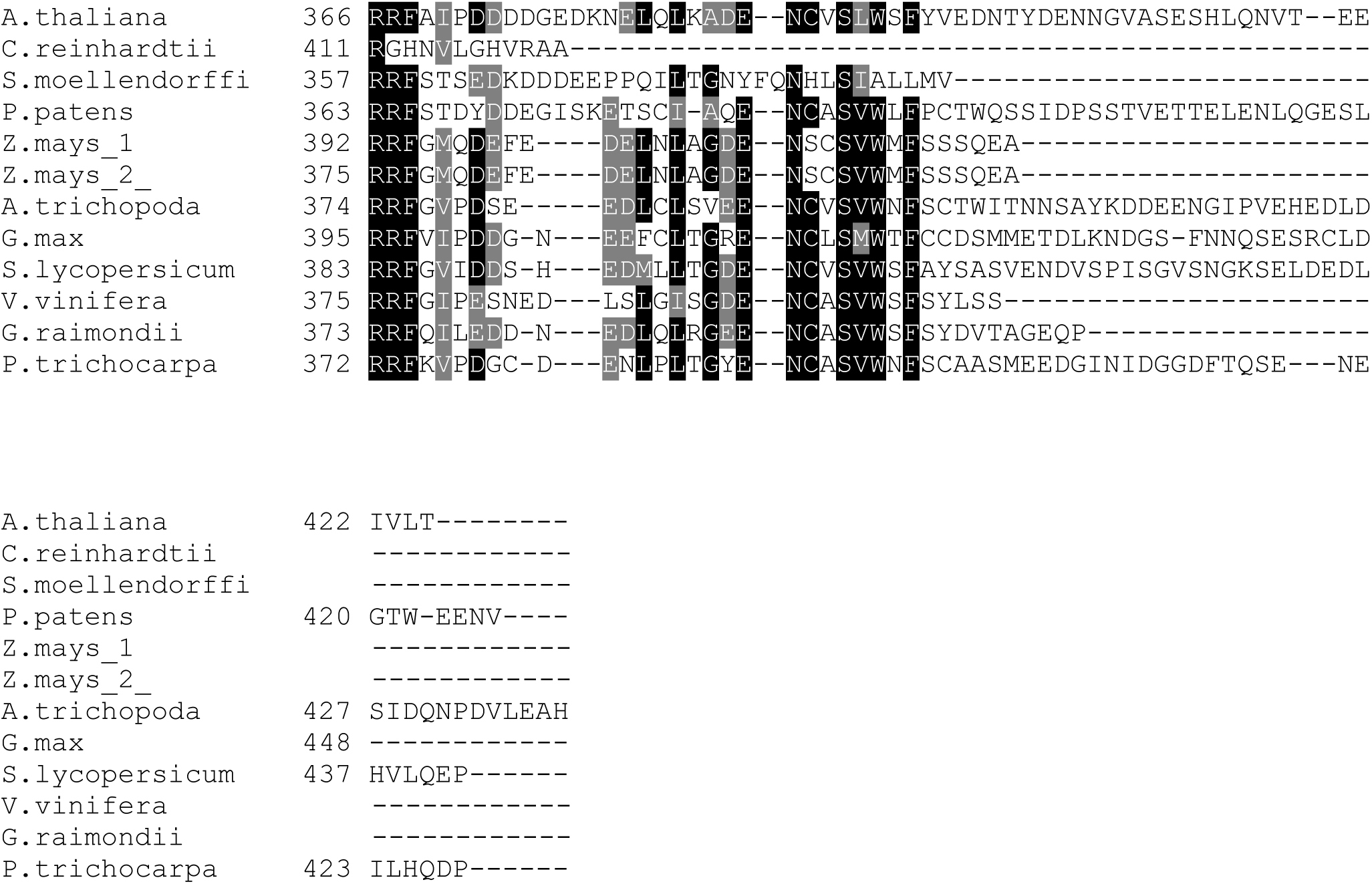
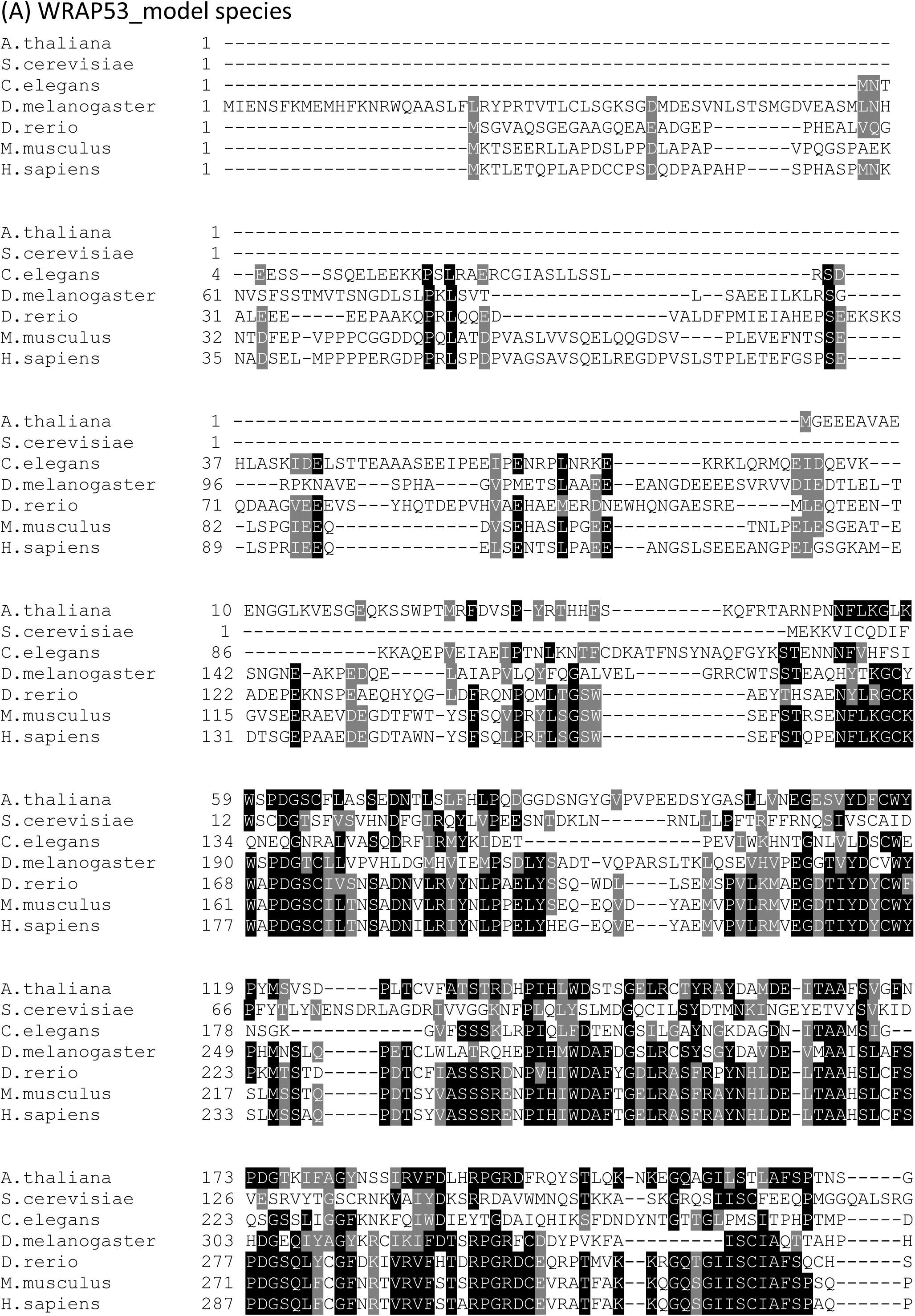
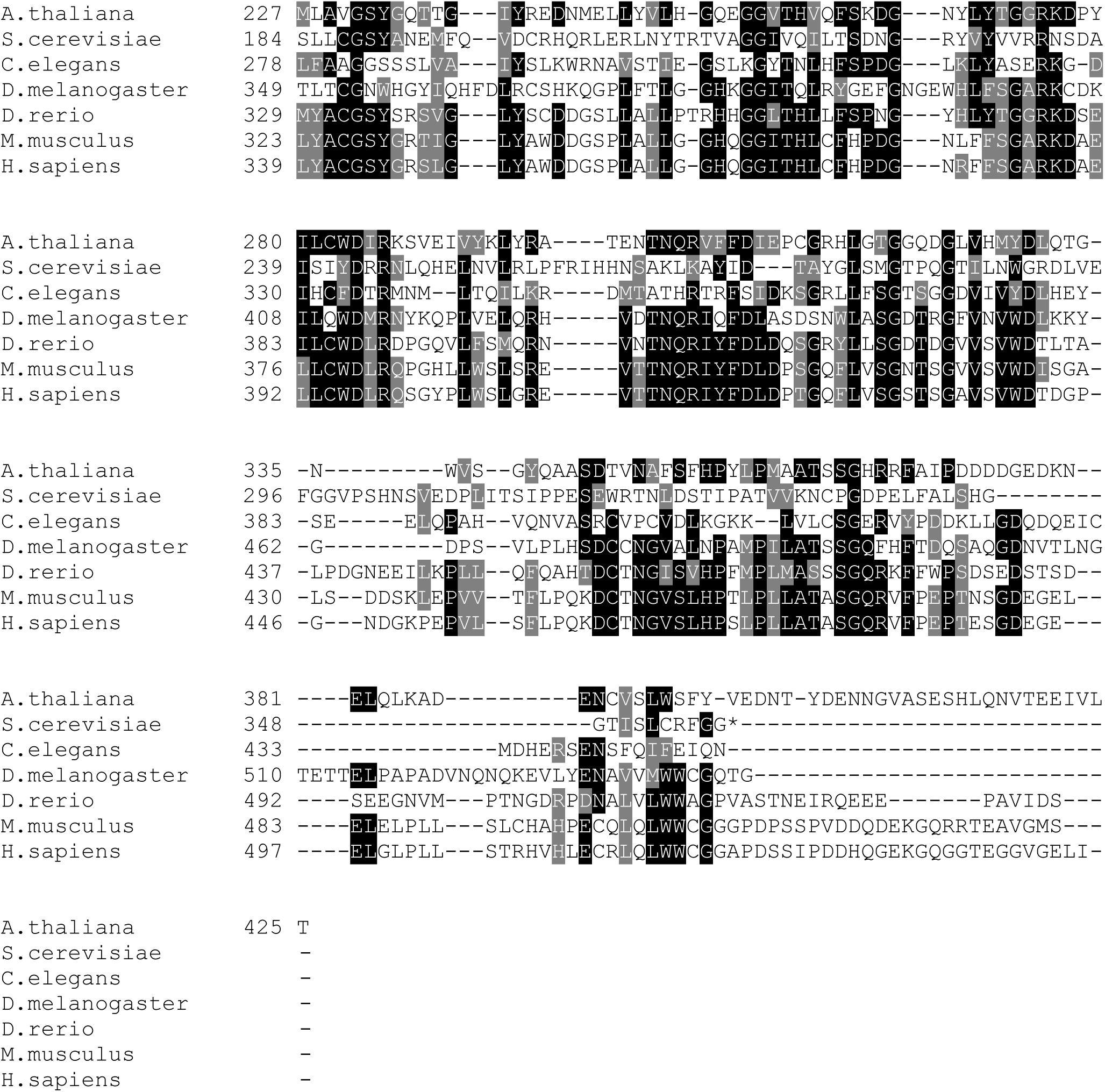
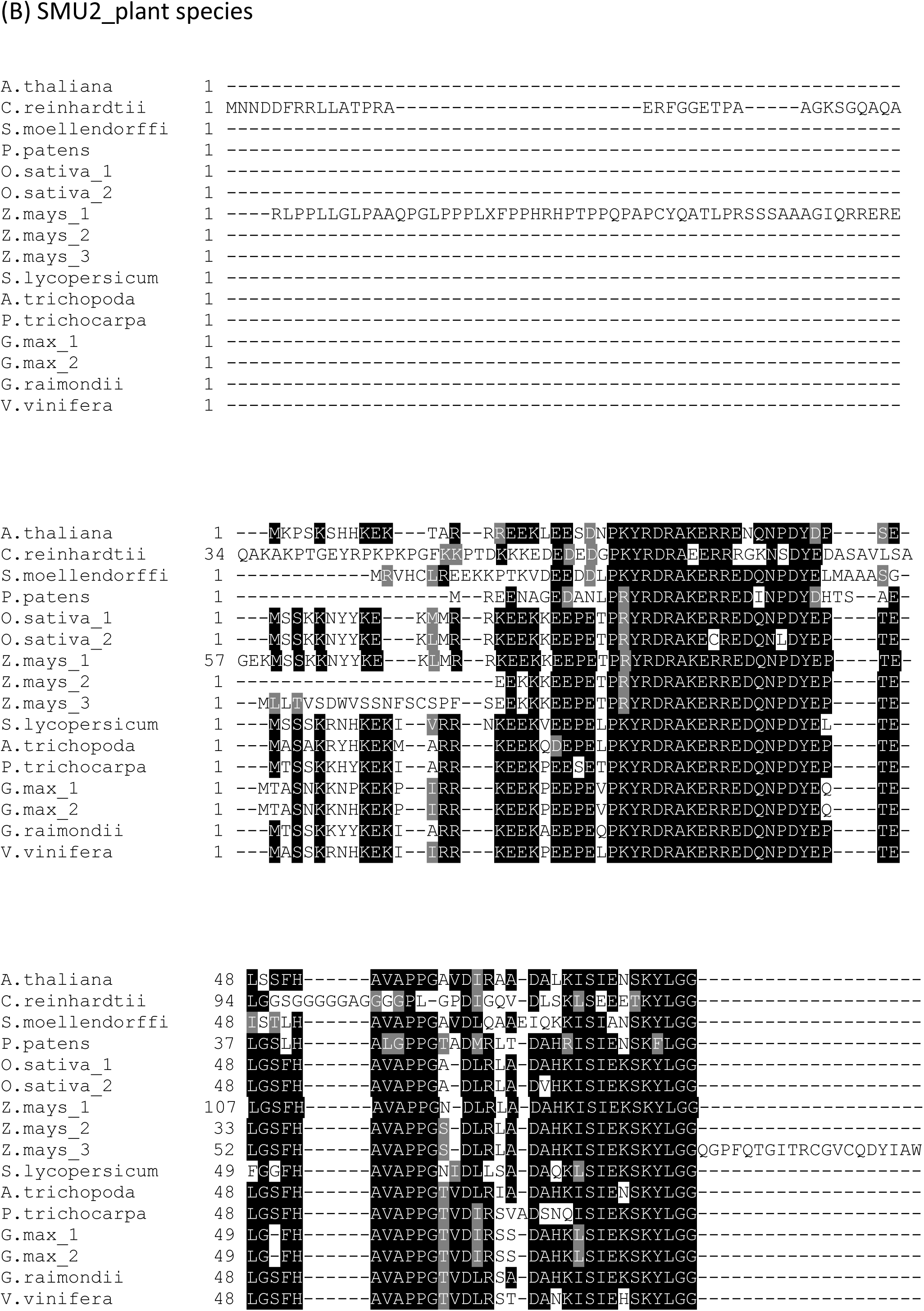
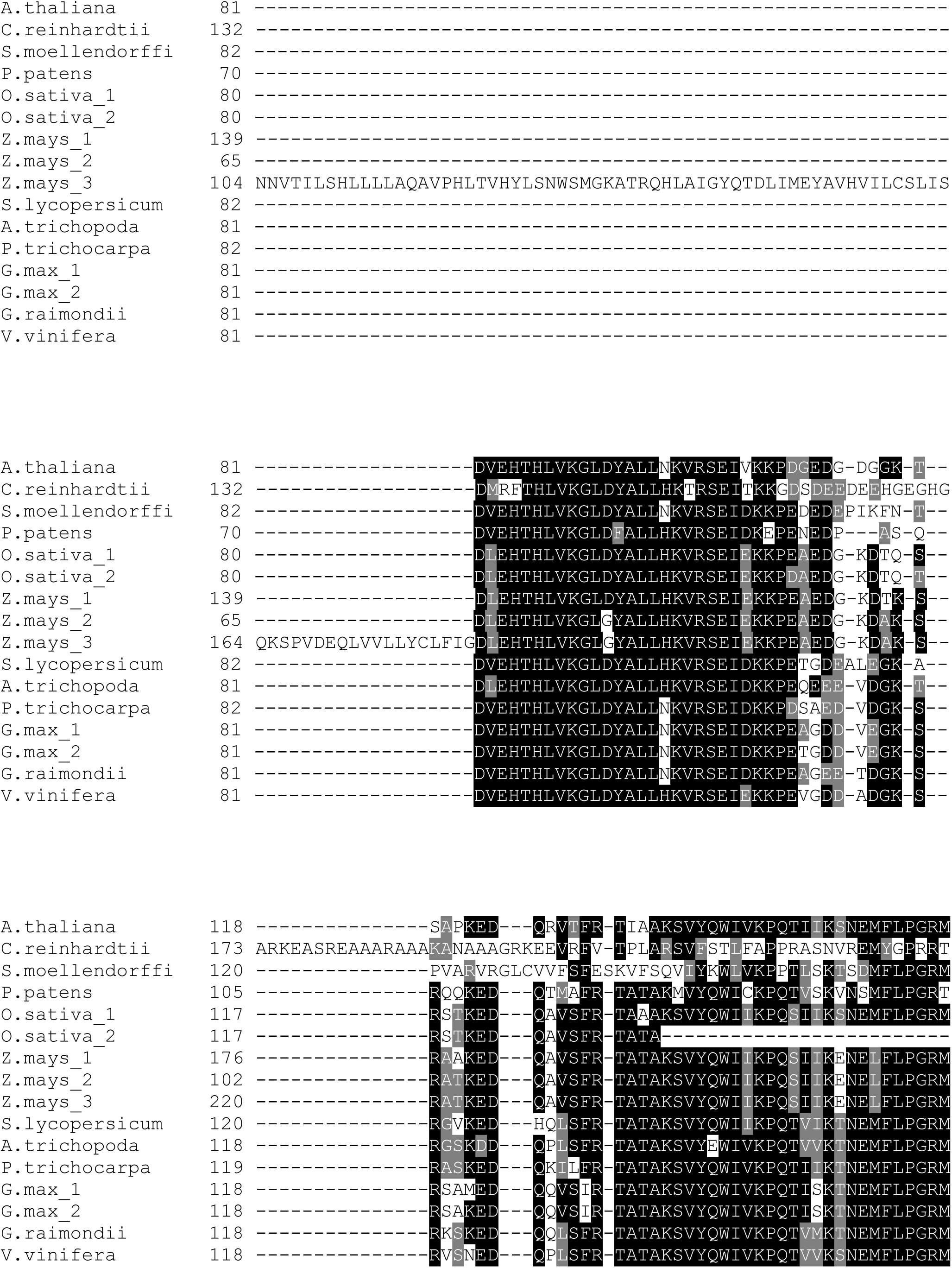
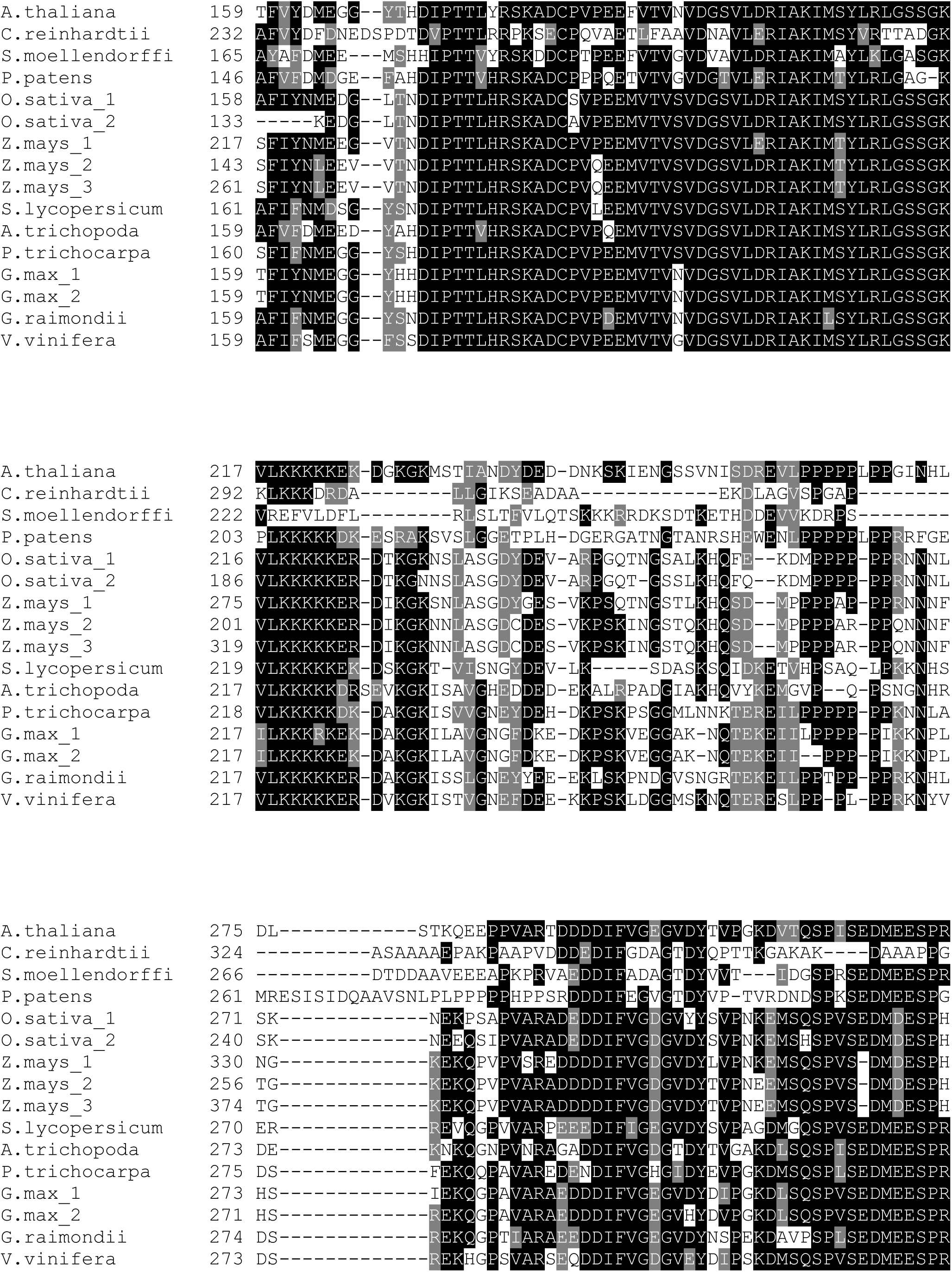
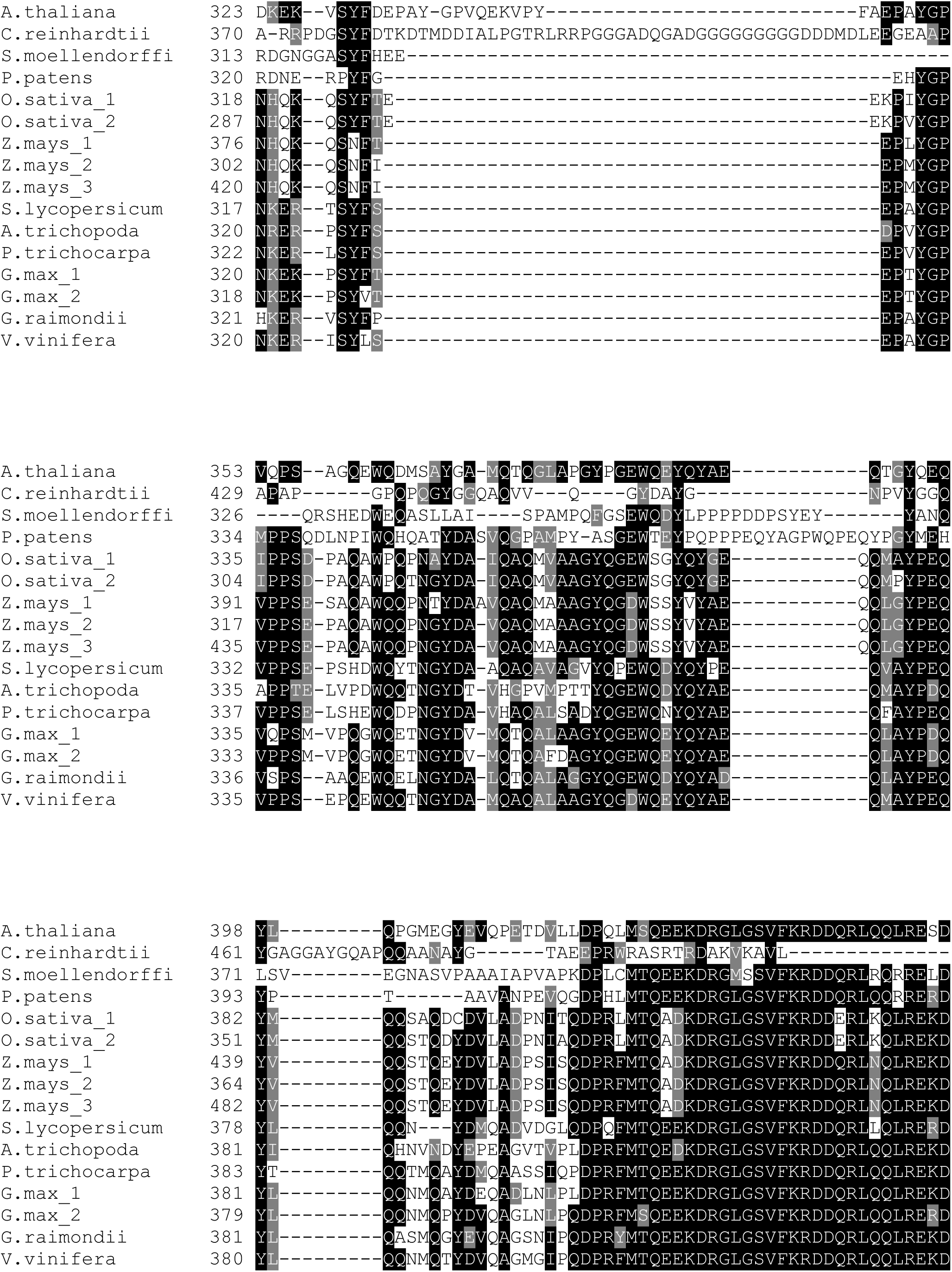
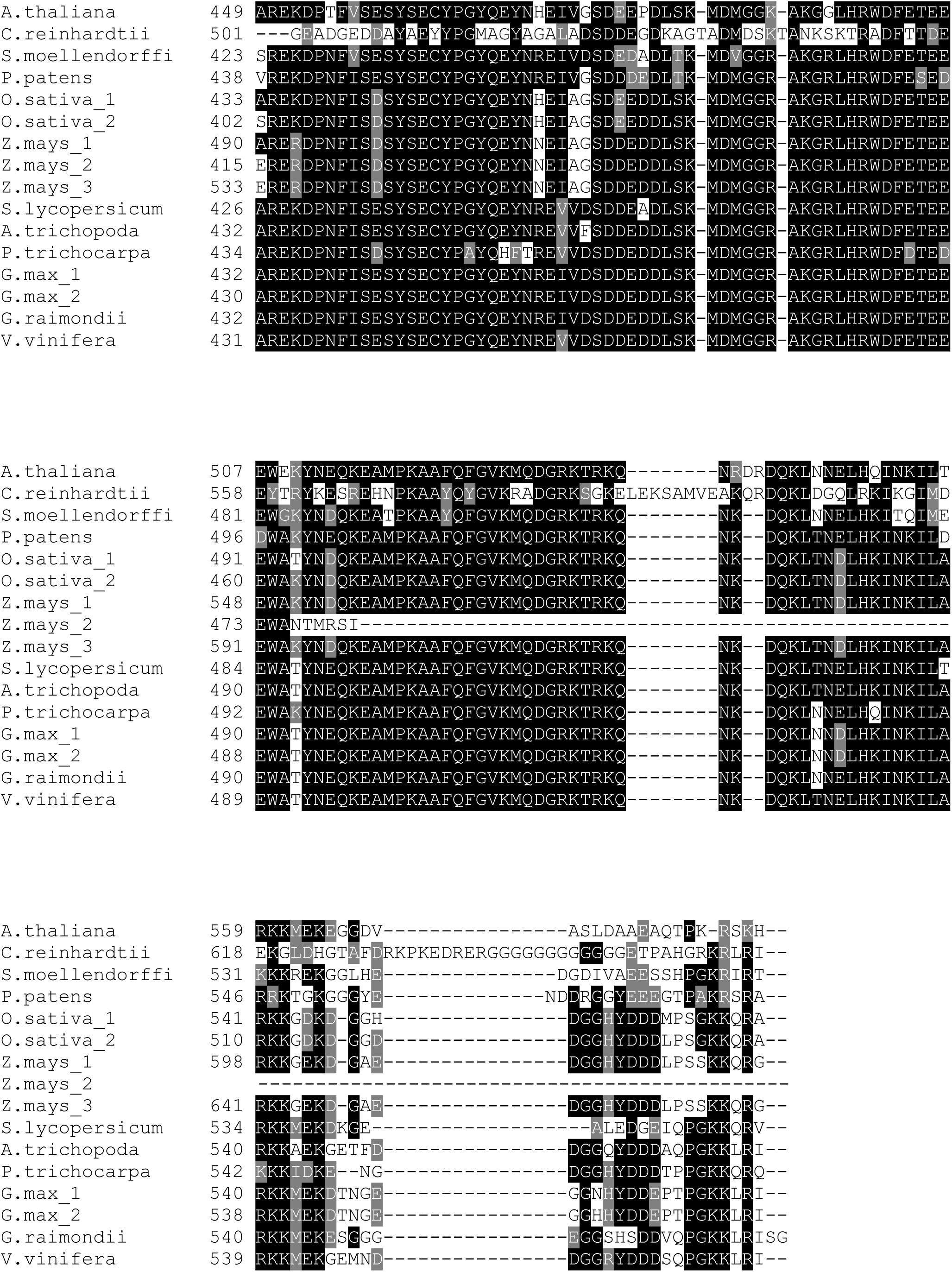
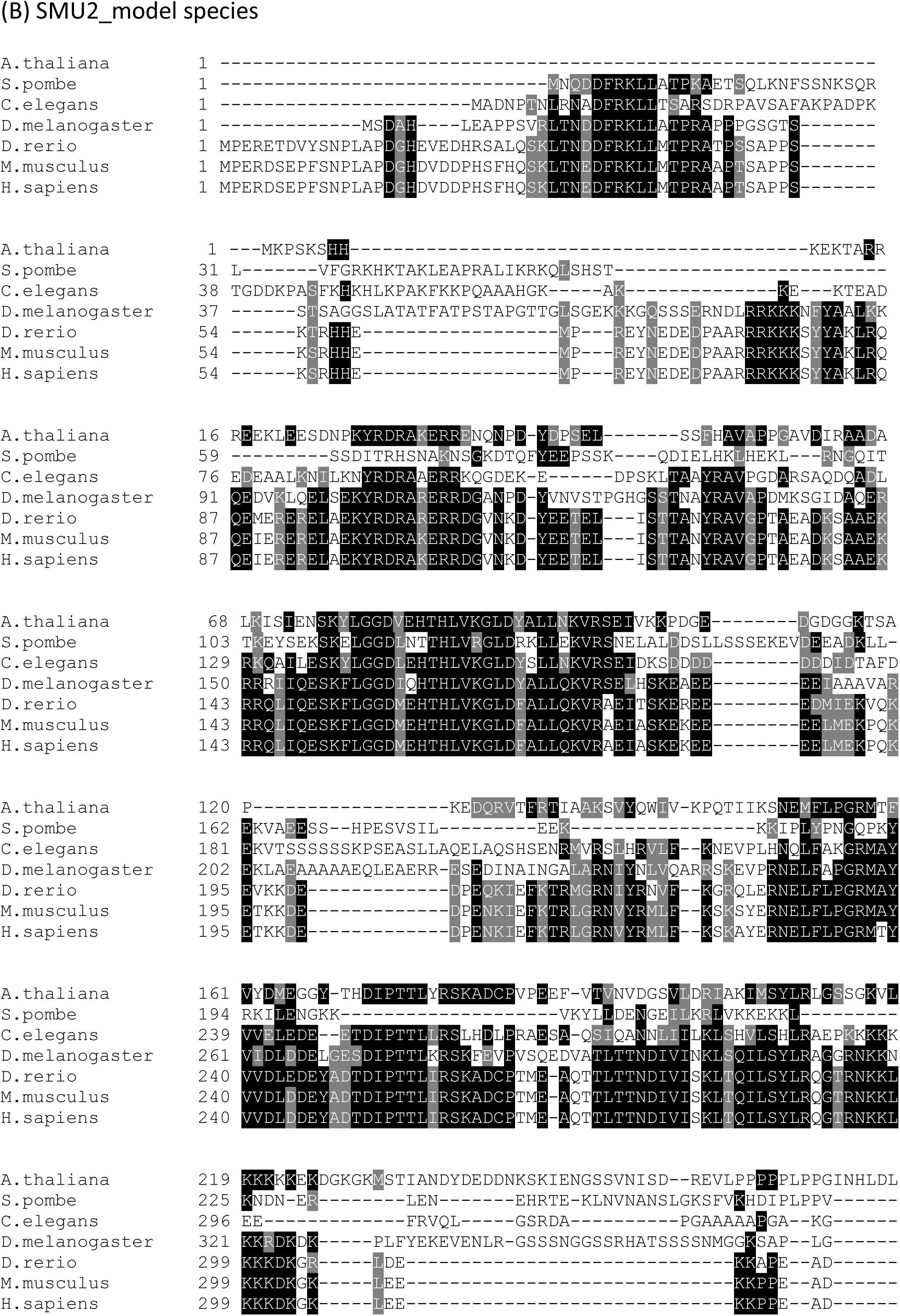
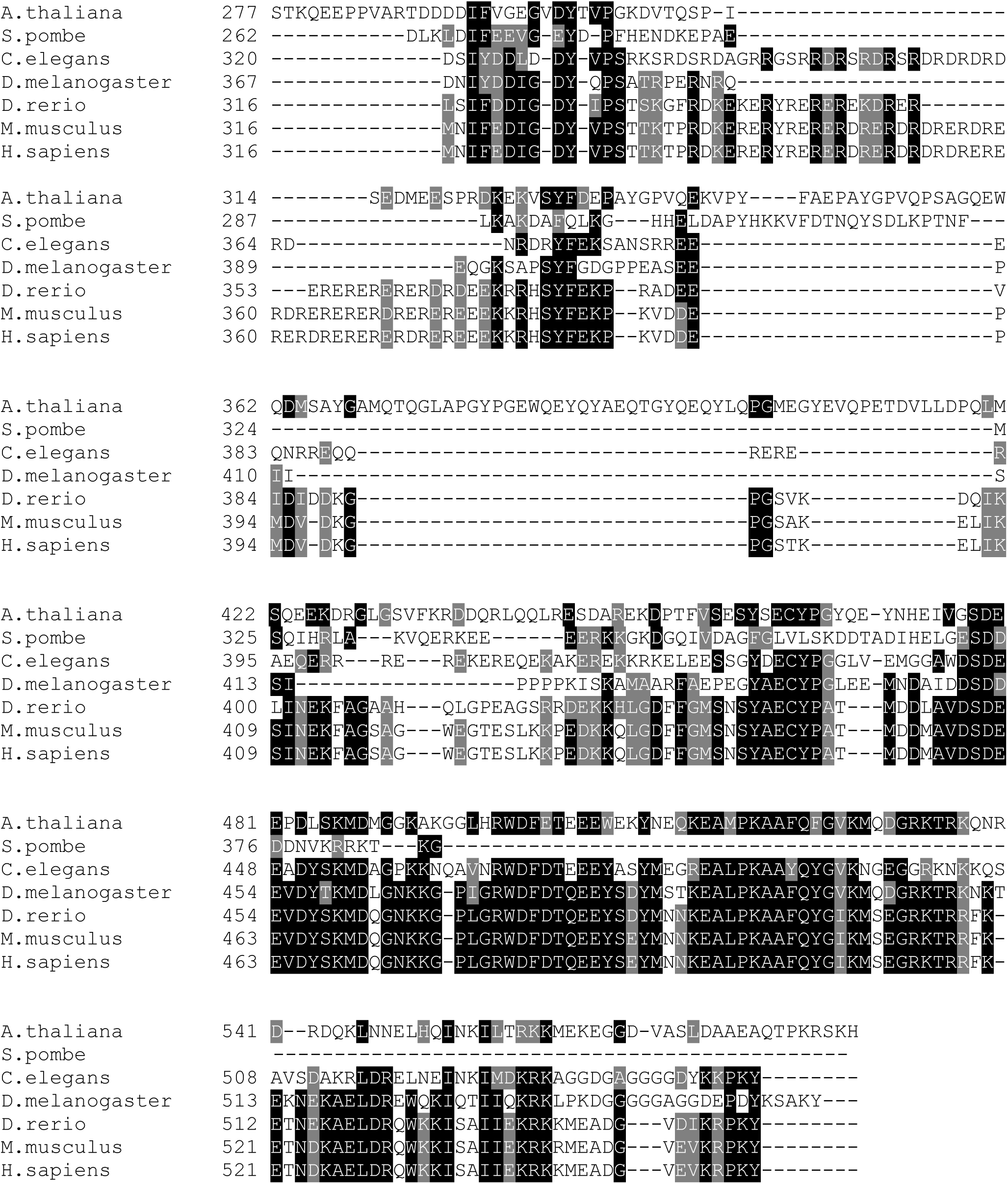
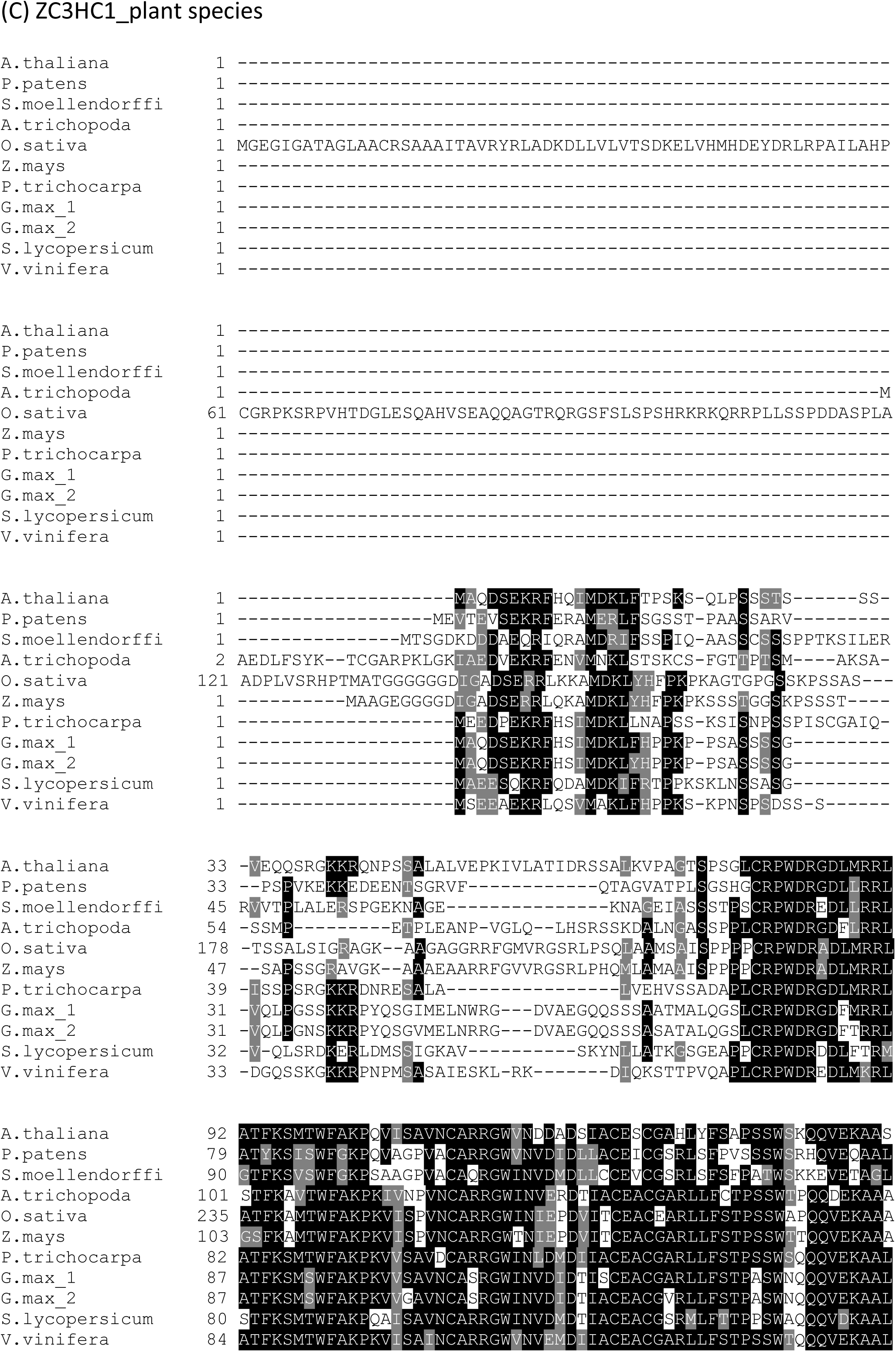
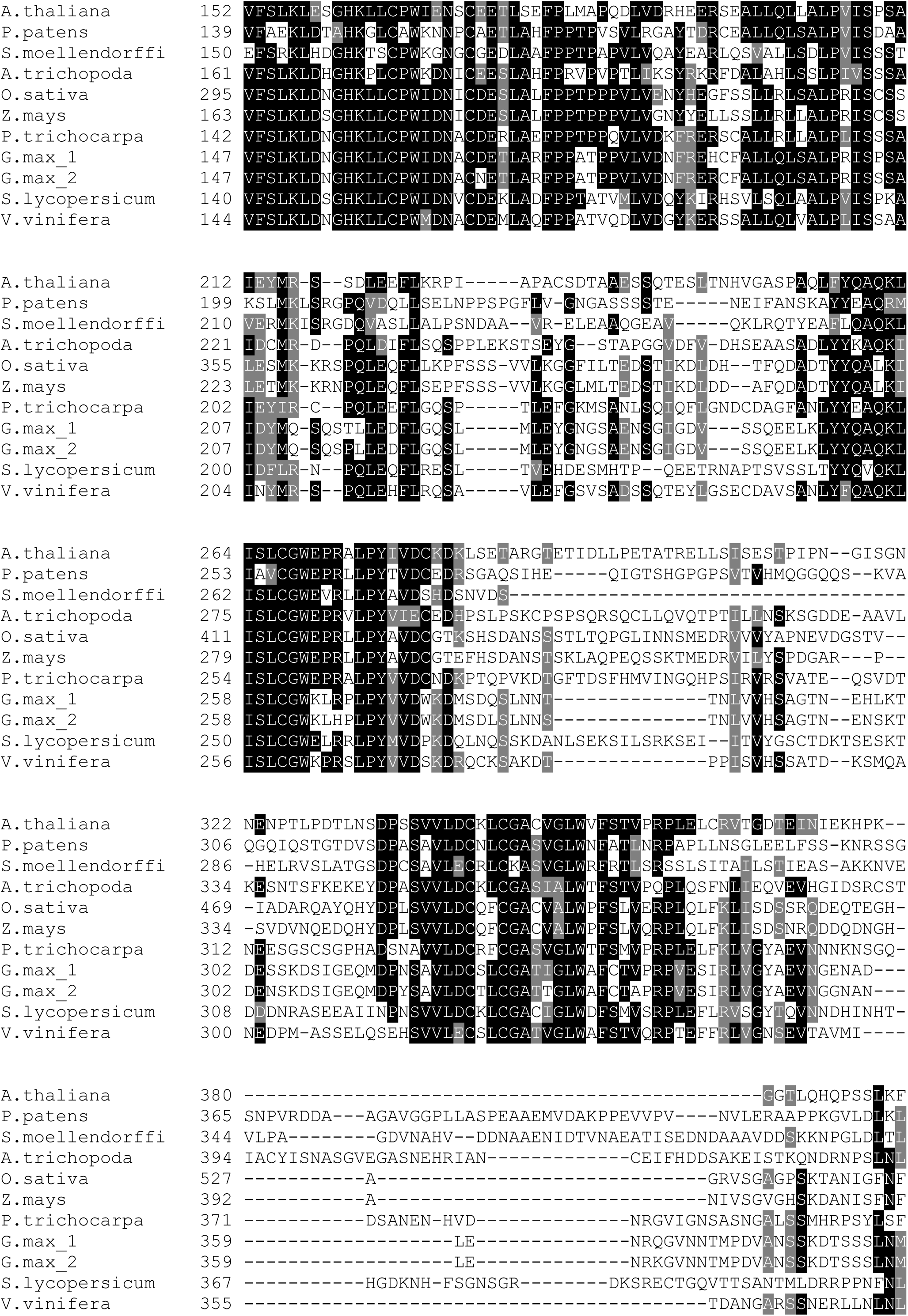
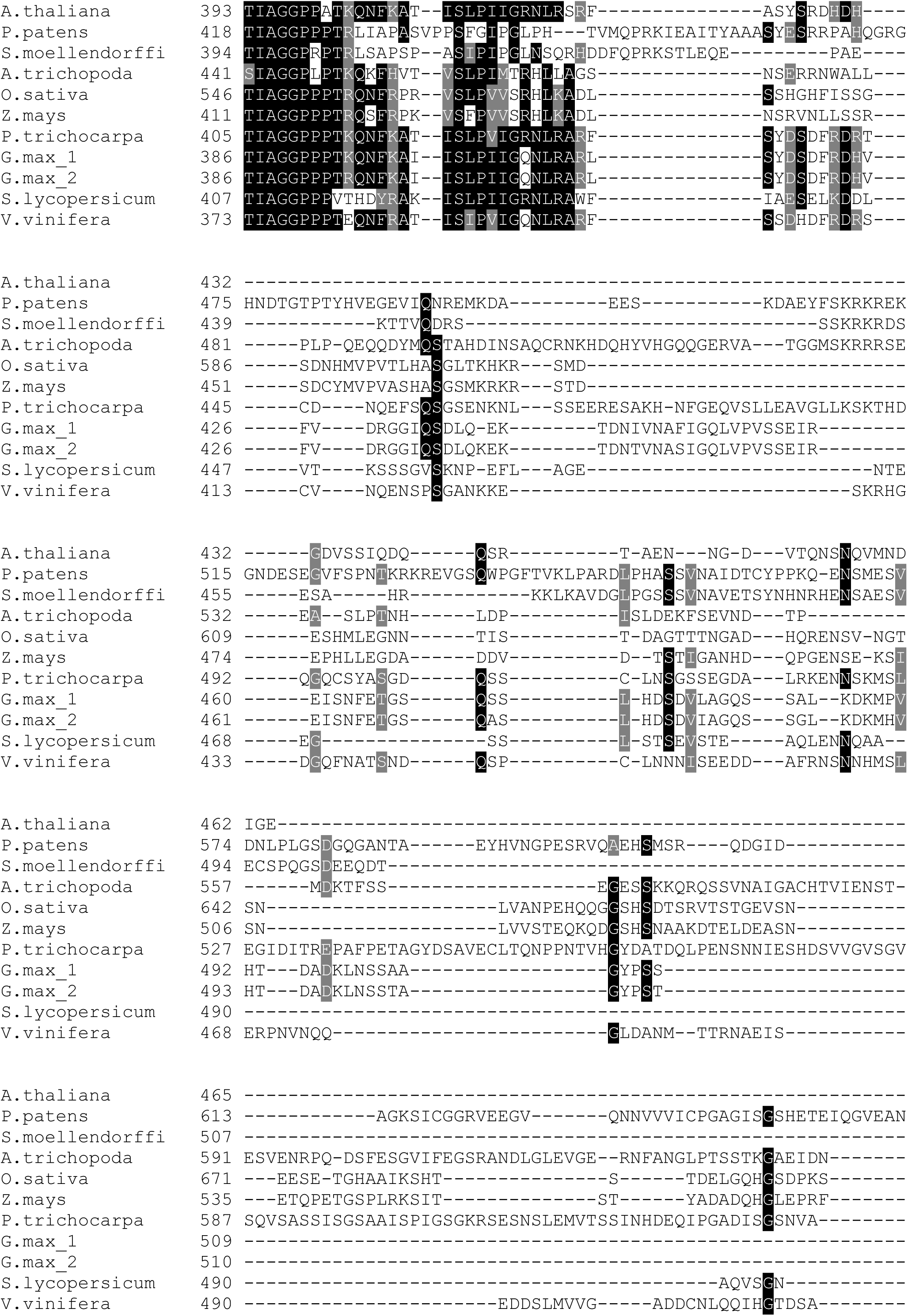
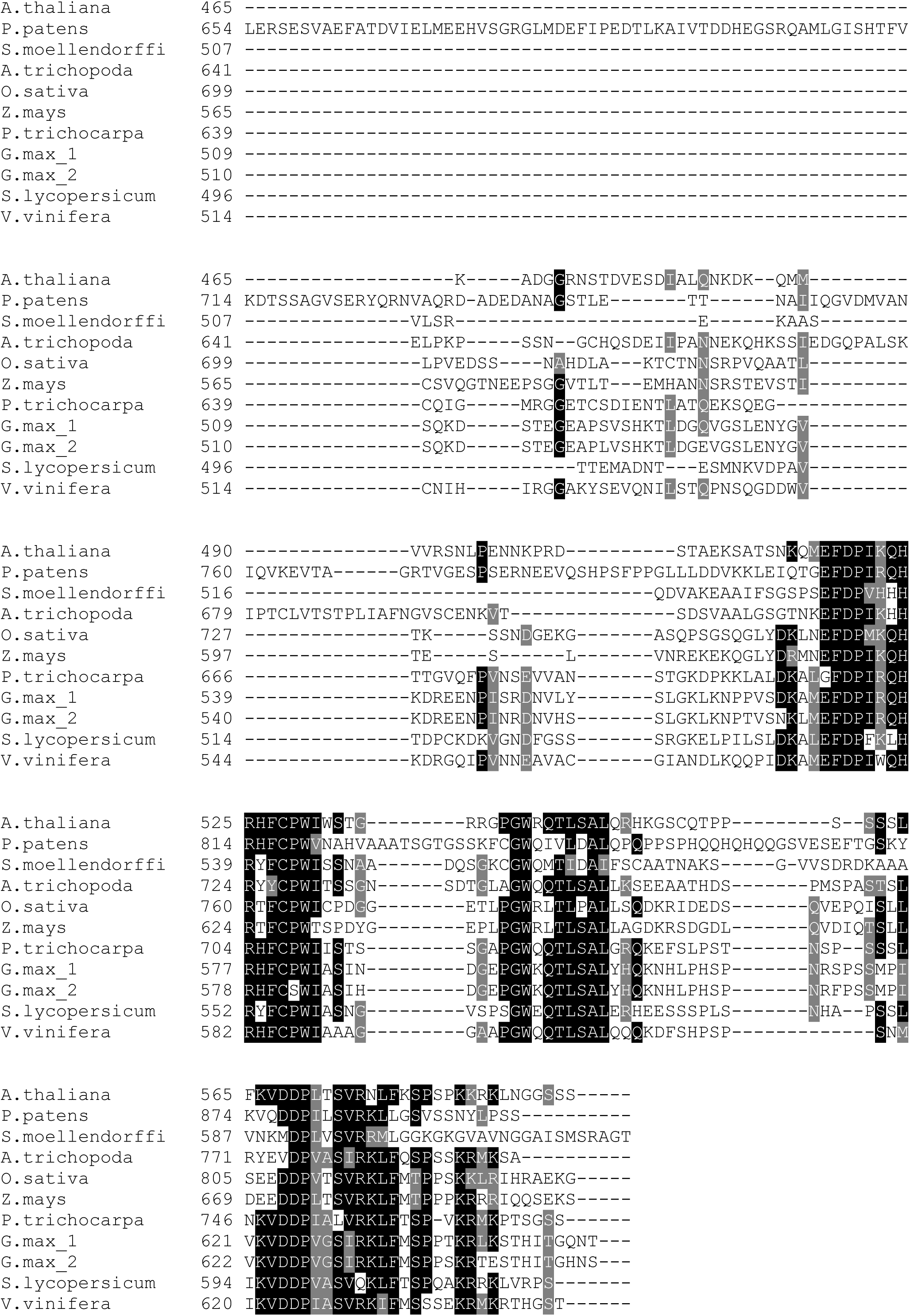
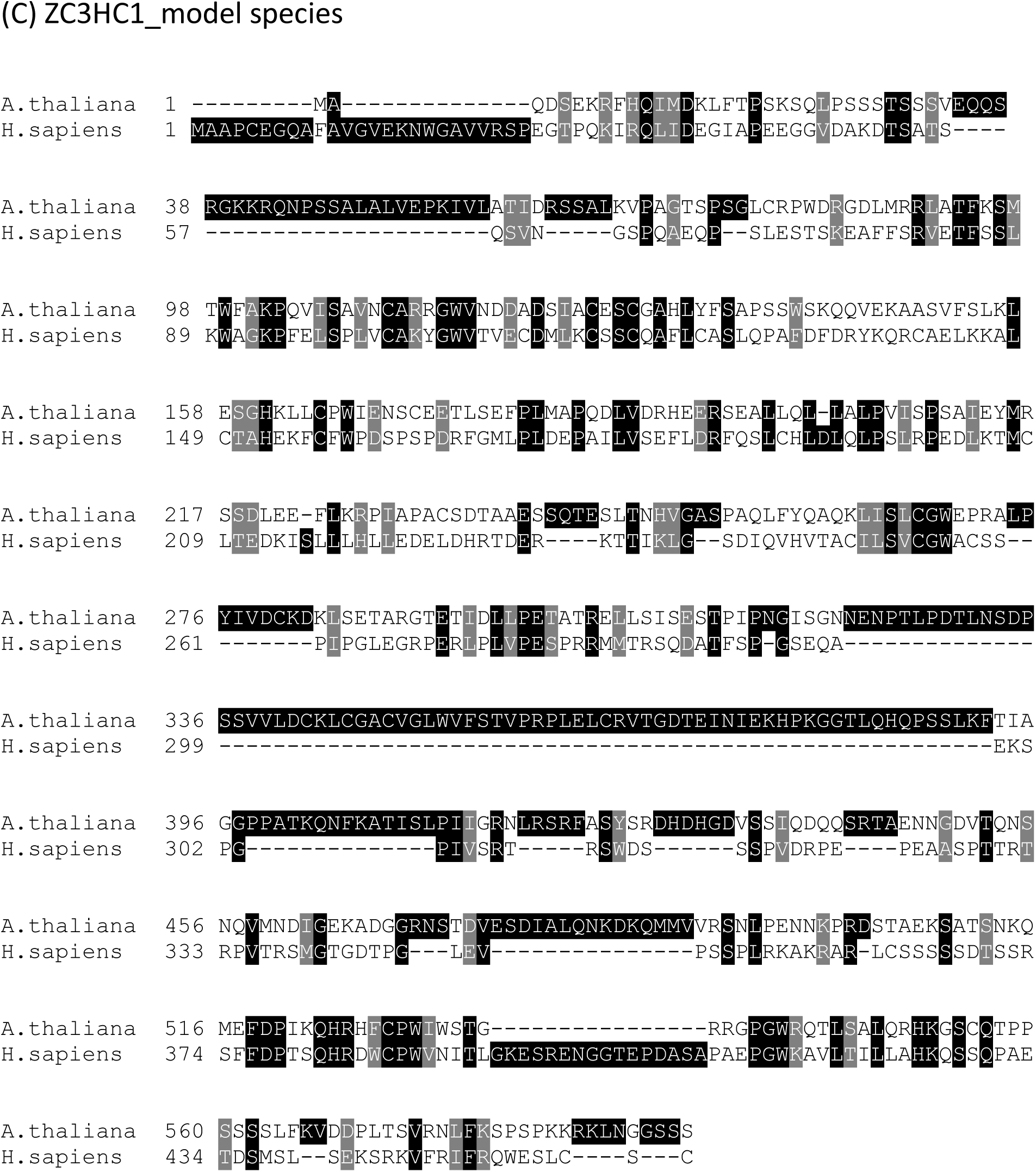
Amino acid sequence alignments. Amino acid sequence alignments of the indicated Arabidopsis proteins (identified in the coilin suppressor screen) with orthologs in other plant species and model species. (A) WRAP53; (B) SMU2; (C) ZC3HC1

**Figure S8:**
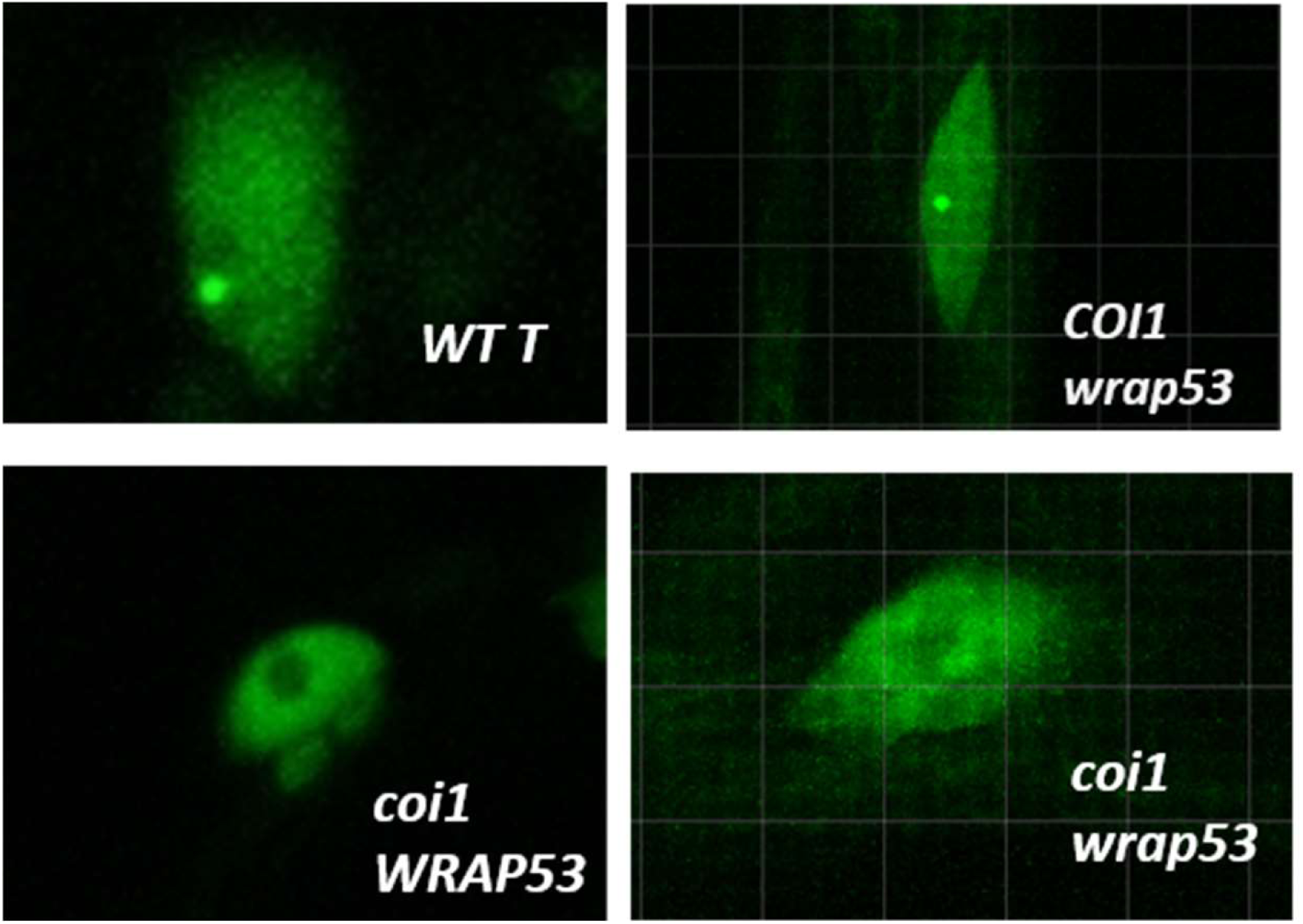
A *wrap53* mutation does not disrupt Cajal bodies in Arabidopsis. To examine whether *wrap53* mutations are able to disrupt CB structure, we used a fluorescence detection procedure described previously (Kanno et al., 2016). Briefly, a *coi1* homozygous mutant was crossed to a transgenic line expressing the CB-specific fluorescence marker U2B”: GFP (Collier *et al*., 2008). The resulting F1 plants were allowed to self-fertilize to produce F2 seeds, which were germinated on solid MS medium containing phosphinothricin (PPT). F2 seedlings were selected for PPT resistance (contributed by the U2B”: GFP line) and a GFP-negative phenotype, indicating absence of the *WT T* locus. Seedlings selected in this way were transferred to soil and later genotyped for the homozygous *coi1* mutation. The presence of fluorescent CBs in trichome nuclei of *WT T* and homozygous *coi1* plants containing the U2B”: GFP gene was assessed using a TCS LSI-III Confocal Microscope System. At least 20 leaf nuclei were examined for each genotype. **Top-left;** *WT T*, fluorescent CBs are visible. **Bottom left**: By contrast, CBs lose fluorescence in a *coi1* mutant that is wild-type for WRAP53. **Top right:** Notably, fluorescent CBs are also visible in the *wrap53* mutant that is wild-type for coilin (COI), indicating that the *wrap53* mutation alone does not disrupt CB integrity. **Bottom right**: CBs are not fluorescent in a *coi1 wrap 53* double mutant, demonstrating that a *wrap53* mutation cannot rescue the loss of CB fluorescence caused by the *coi1* mutation.

### Supplementary Tables

**Table S1:**
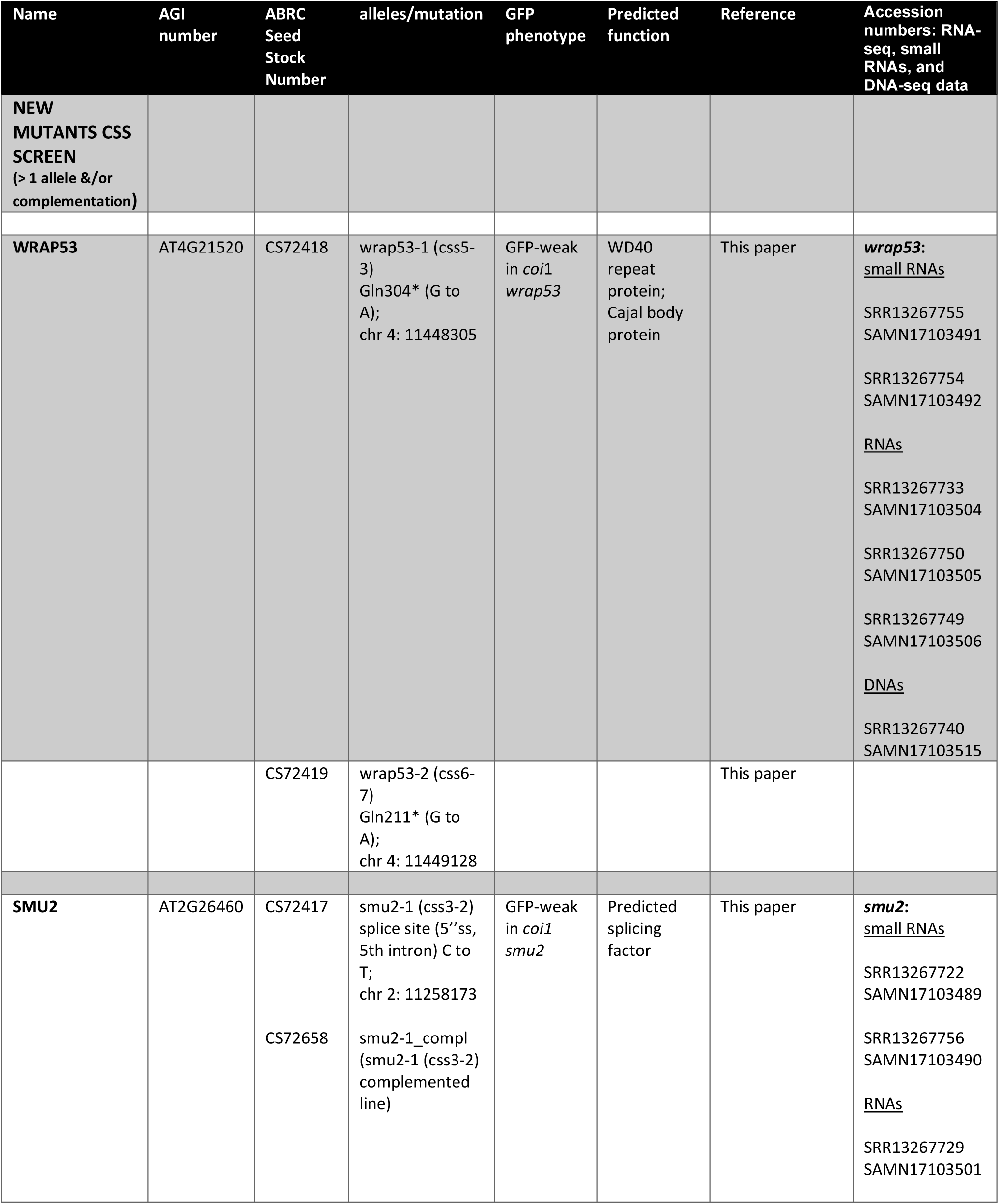
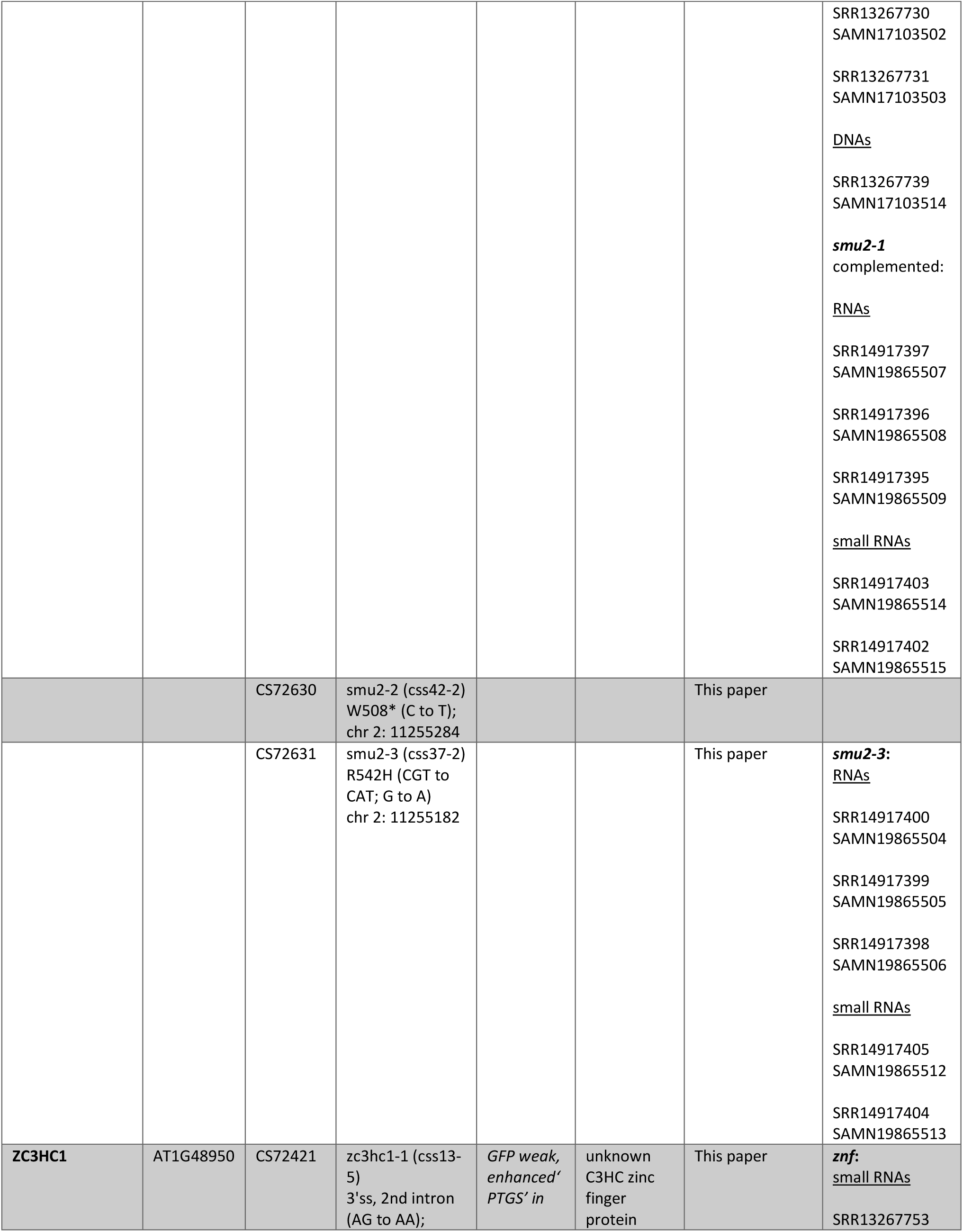
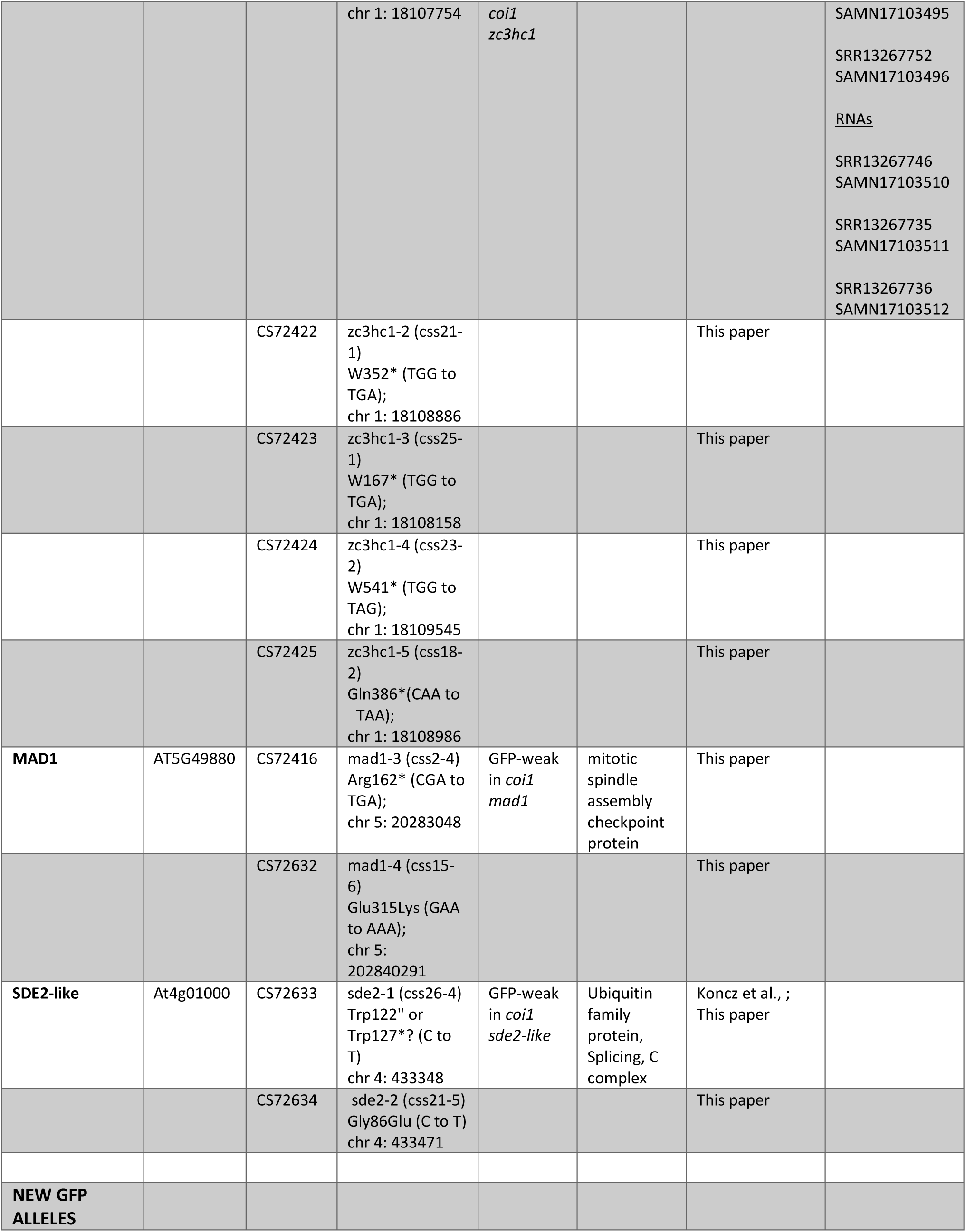
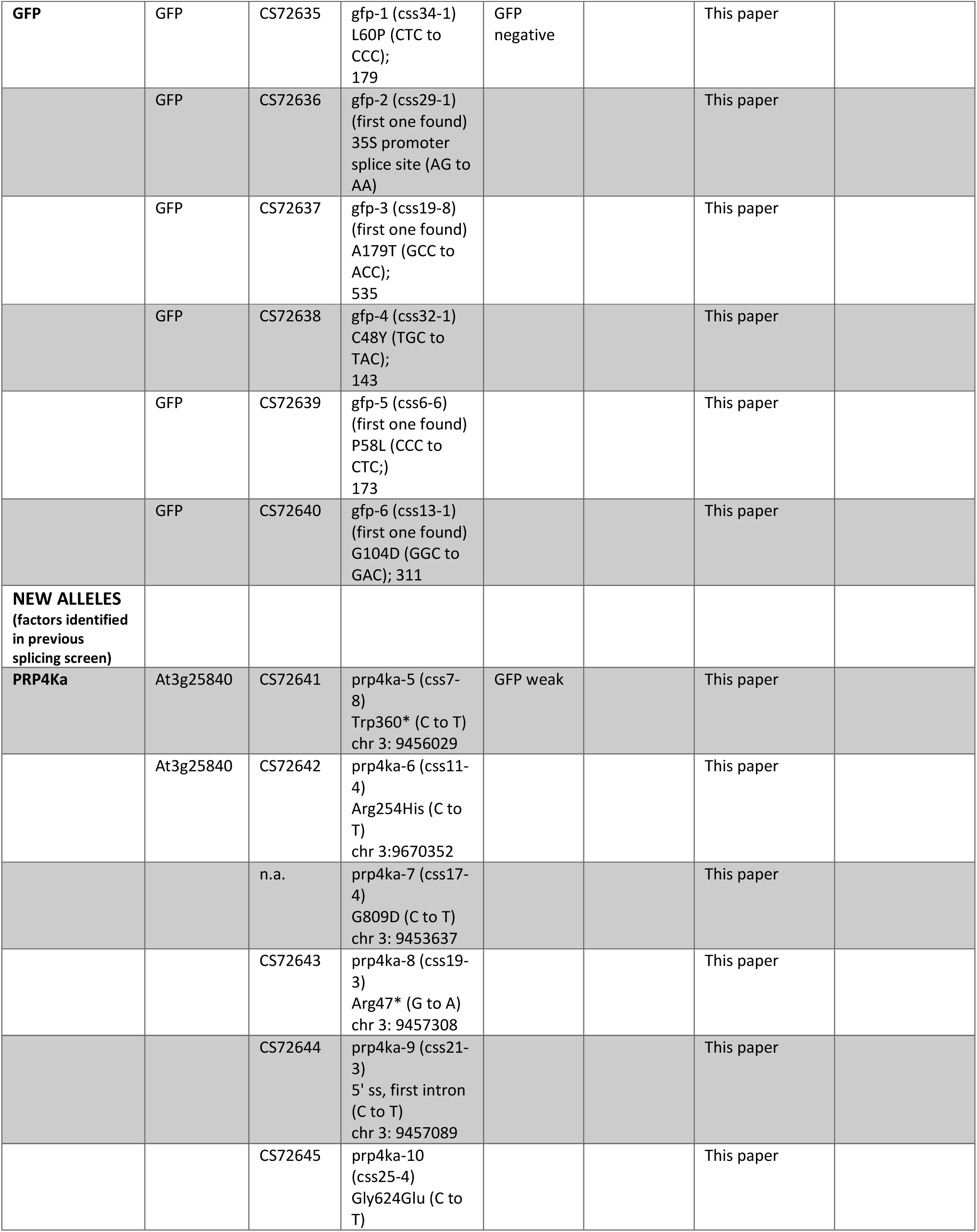
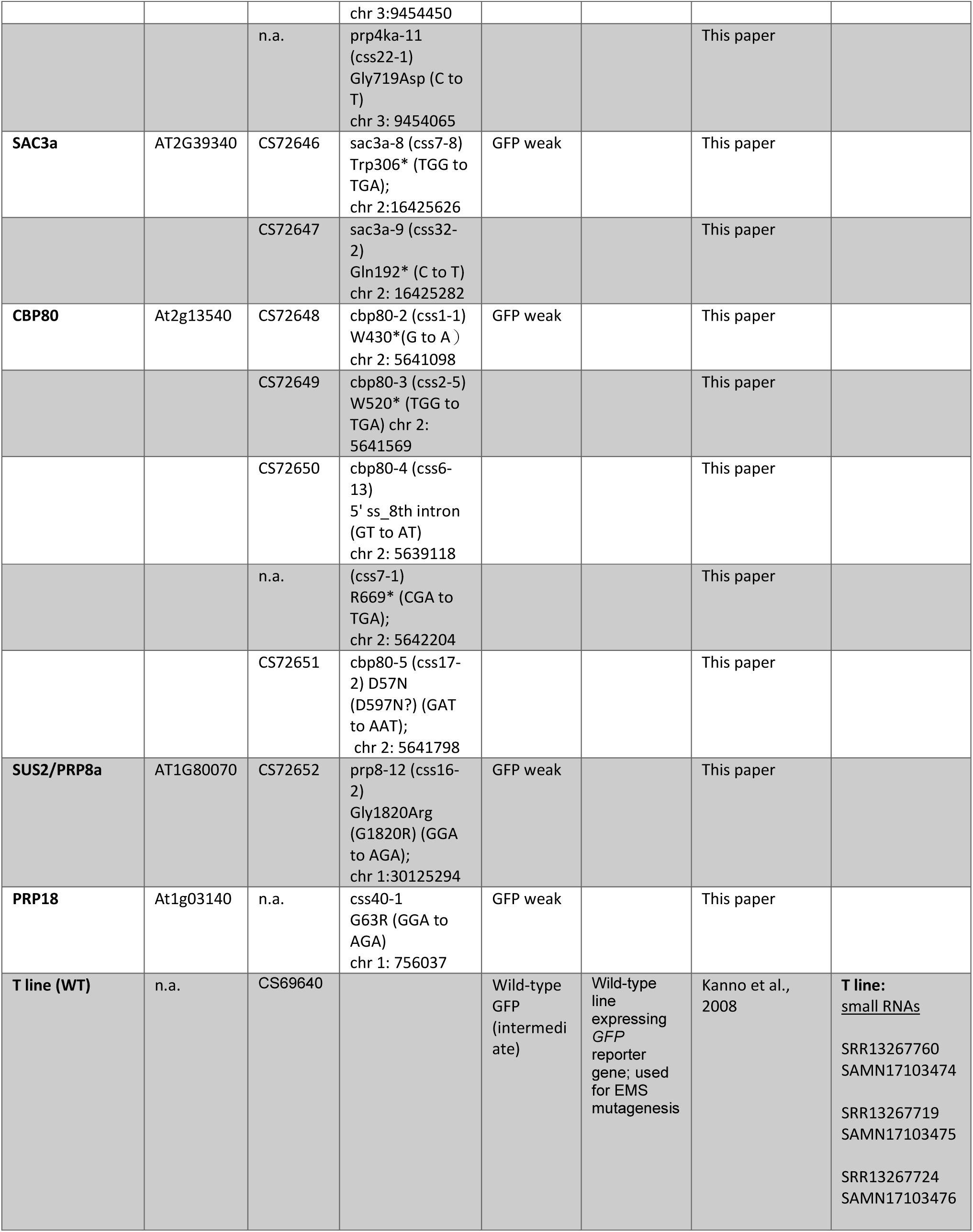
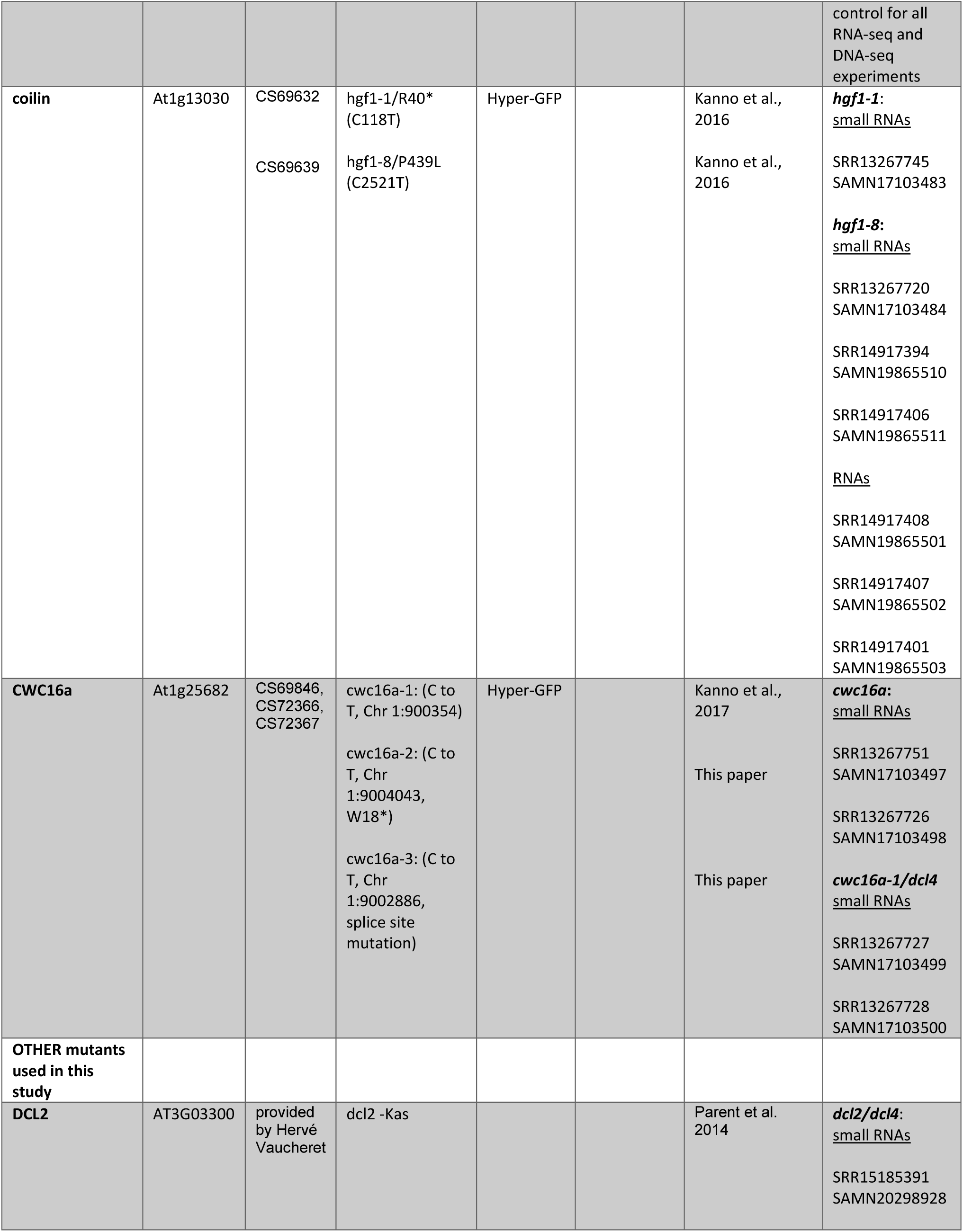
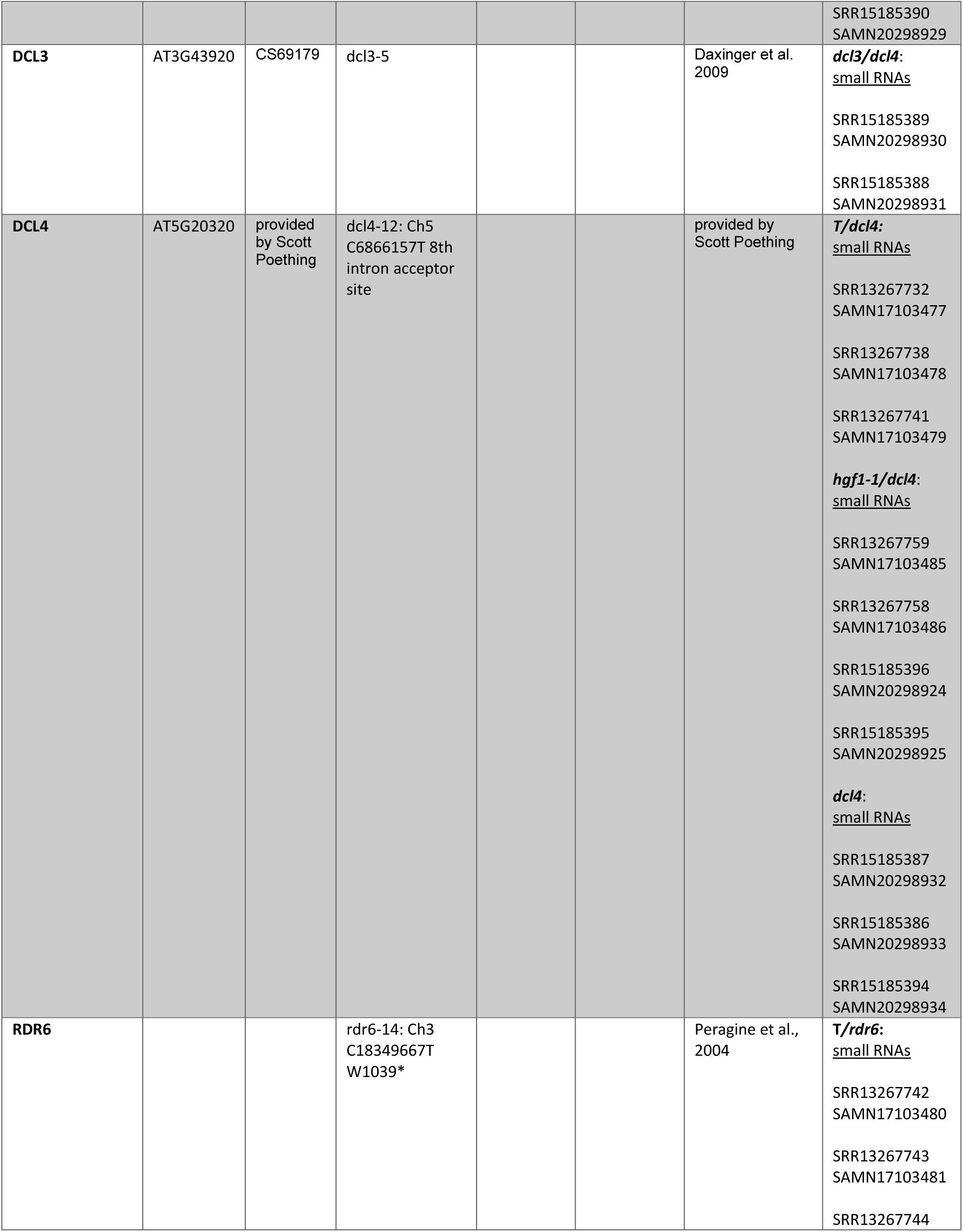
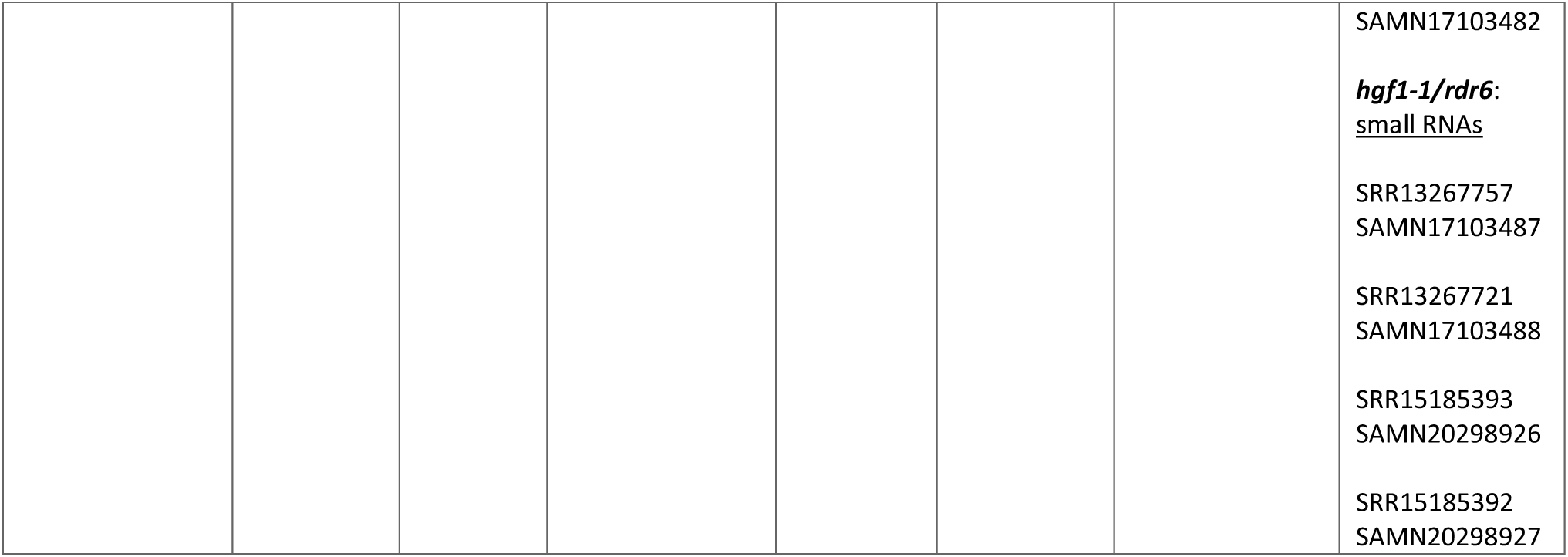
Mutants identified in the coilin suppressor screen and other mutants used in this study.

| Name | AGI number | ABRC Seed Stock Number | alleles/mutation | GFP phenotype | Predicted function | Reference | Accession numbers: RNA-seq, small RNAs, and DNA-seq data |
| --- | --- | --- | --- | --- | --- | --- | --- |
| **NEW MUTANTS CSS SCREEN**<br>(> 1 allele &/or complementation) |  |  |  |  |  |  |  |
| **WRAP53** | AT4G21520 | CS72418 | wrap53-1 (css5-3)<br>Gln304* (G to A);<br>chr 4: 11448305 | GFP-weak in *coi1*<br>*wrap53* | WD40 repeat protein; Cajal body protein | This paper | ***wrap53*:**<br><u>small RNAs</u><br>SRR13267755<br>SAMN17103491<br>SRR13267754<br>SAMN17103492<br><u>RNAs</u><br>SRR13267733<br>SAMN17103504<br>SRR13267750<br>SAMN17103505<br>SRR13267749<br>SAMN17103506<br><u>DNAs</u><br>SRR13267740<br>SAMN17103515 |
|  |  | CS72419 | wrap53-2 (css6-7)<br>Gln211* (G to A);<br>chr 4: 11449128 |  |  | This paper |  |
| **SMU2** | AT2G26460 | CS72417<br>CS72658 | smu2-1 (css3-2) splice site (5''ss, 5th intron) C to T;<br>chr 2: 11258173<br>smu2-1_compl (smu2-1 (css3-2) complemented line) | GFP-weak in *coi1*<br>*smu2* | Predicted splicing factor | This paper | ***smu2*:**<br><u>small RNAs</u><br>SRR13267722<br>SAMN17103489<br>SRR13267756<br>SAMN17103490<br><u>RNAs</u><br>SRR13267729<br>SAMN17103501 |
|  |  |  |  |  |  |  | SRR13267730<br>SAMN17103502<br>SRR13267731<br>SAMN17103503<br><u>DNA</u>s<br>SRR13267739<br>SAMN17103514<br>***smu2-1***<br>complemented:<br><u>RNA</u>s<br>SRR14917397<br>SAMN19865507<br>SRR14917396<br>SAMN19865508<br>SRR14917395<br>SAMN19865509<br><u>small RNA</u>s<br>SRR14917403<br>SAMN19865514<br>SRR14917402<br>SAMN19865515 |
|  |  | CS72630 | smu2-2 (css42-2)<br>W508* (C to T);<br>chr 2: 11255284 |  |  | This paper |  |
|  |  | CS72631 | smu2-3 (css37-2)<br>R542H (CGT to<br>CAT; G to A)<br>chr 2: 11255182 |  |  | This paper | ***smu2-3:***<br><u>RNA</u>s<br>SRR14917400<br>SAMN19865504<br>SRR14917399<br>SAMN19865505<br>SRR14917398<br>SAMN19865506<br><u>small RNA</u>s<br>SRR14917405<br>SAMN19865512<br>SRR14917404<br>SAMN19865513 |
| **ZC3HC1** | AT1G48950 | CS72421 | zc3hc1-1 (css13-5)<br>3'ss, 2nd intron<br>(AG to AA); | *GFP weak, enhanced'*<br>*PTGS' in* | unknown<br>C3HC zinc<br>finger<br>protein | This paper | ***znf:***<br><u>small RNA</u>s<br>SRR13267753 |
|  |  |  | chr 1: 18107754 | *coi1*<br>*zc3hc1* |  |  | SAMN17103495<br>SRR13267752<br>SAMN17103496<br><u>RNAs</u><br>SRR13267746<br>SAMN17103510<br>SRR13267735<br>SAMN17103511<br>SRR13267736<br>SAMN17103512 |
|  |  | CS72422 | zc3hc1-2 (css21-1)<br>W352* (TGG to TGA);<br>chr 1: 18108886 |  |  | This paper |  |
|  |  | CS72423 | zc3hc1-3 (css25-1)<br>W167* (TGG to TGA);<br>chr 1: 18108158 |  |  | This paper |  |
|  |  | CS72424 | zc3hc1-4 (css23-2)<br>W541* (TGG to TAG);<br>chr 1: 18109545 |  |  | This paper |  |
|  |  | CS72425 | zc3hc1-5 (css18-2)<br>Gln386* (CAA to TAA);<br>chr 1: 18108986 |  |  | This paper |  |
| **MAD1** | AT5G49880 | CS72416 | mad1-3 (css2-4)<br>Arg162* (CGA to TGA);<br>chr 5: 20283048 | GFP-weak in *coi1*<br>*mad1* | mitotic spindle assembly checkpoint protein | This paper |  |
|  |  | CS72632 | mad1-4 (css15-6)<br>Glu315Lys (GAA to AAA);<br>chr 5: 202840291 |  |  | This paper |  |
| **SDE2-like** | At4g01000 | CS72633 | sde2-1 (css26-4)<br>Trp122" or Trp127*? (C to T)<br>chr 4: 433348 | GFP-weak in *coi1*<br>*sde2-like* | Ubiquitin family protein, Splicing, C complex | Koncz et al., ;<br>This paper |  |
|  |  | CS72634 | sde2-2 (css21-5)<br>Gly86Glu (C to T)<br>chr 4: 433471 |  |  | This paper |  |
| **NEW GFP ALLELES** |  |  |  |  |  |  |  |
| **GFP** | GFP | CS72635 | gfp-1 (css34-1)<br>L60P (CTC to CCC);<br>179 | GFP negative |  | This paper |  |
|  | GFP | CS72636 | gfp-2 (css29-1)<br>(first one found)<br>35S promoter<br>splice site (AG to AA) |  |  | This paper |  |
|  | GFP | CS72637 | gfp-3 (css19-8)<br>(first one found)<br>A179T (GCC to ACC);<br>535 |  |  | This paper |  |
|  | GFP | CS72638 | gfp-4 (css32-1)<br>C48Y (TGC to TAC);<br>143 |  |  | This paper |  |
|  | GFP | CS72639 | gfp-5 (css6-6)<br>(first one found)<br>P58L (CCC to CTC);<br>173 |  |  | This paper |  |
|  | GFP | CS72640 | gfp-6 (css13-1)<br>(first one found)<br>G104D (GGC to GAC); 311 |  |  | This paper |  |
| **NEW ALLELES**<br>(factors identified in previous splicing screen) |  |  |  |  |  |  |  |
| **PRP4Ka** | At3g25840 | CS72641 | prp4ka-5 (css7-8)<br>Trp360* (C to T)<br>chr 3: 9456029 | GFP weak |  | This paper |  |
|  | At3g25840 | CS72642 | prp4ka-6 (css11-4)<br>Arg254His (C to T)<br>chr 3:9670352 |  |  | This paper |  |
|  |  | n.a. | prp4ka-7 (css17-4)<br>G809D (C to T)<br>chr 3: 9453637 |  |  | This paper |  |
|  |  | CS72643 | prp4ka-8 (css19-3)<br>Arg47* (G to A)<br>chr 3: 9457308 |  |  | This paper |  |
|  |  | CS72644 | prp4ka-9 (css21-3)<br>5' ss, first intron<br>(C to T)<br>chr 3: 9457089 |  |  | This paper |  |
|  |  | CS72645 | prp4ka-10<br>(css25-4)<br>Gly624Glu (C to T) |  |  | This paper |  |
|  |  |  | chr 3:9454450 |  |  |  |  |
|  |  | n.a. | prp4ka-11<br>(css22-1)<br>Gly719Asp (C to T)<br>chr 3: 9454065 |  |  | This paper |  |
| **SAC3a** | AT2G39340 | CS72646 | sac3a-8 (css7-8)<br>Trp306* (TGG to TGA);<br>chr 2:16425626 | GFP weak |  | This paper |  |
|  |  | CS72647 | sac3a-9 (css32-2)<br>Gln192* (C to T)<br>chr 2: 16425282 |  |  | This paper |  |
| **CBP80** | At2g13540 | CS72648 | cbp80-2 (css1-1)<br>W430*(G to A )<br>chr 2: 5641098 | GFP weak |  | This paper |  |
|  |  | CS72649 | cbp80-3 (css2-5)<br>W520* (TGG to TGA) chr 2:<br>5641569 |  |  | This paper |  |
|  |  | CS72650 | cbp80-4 (css6-13)<br>5' ss_8th intron (GT to AT)<br>chr 2: 5639118 |  |  | This paper |  |
|  |  | n.a. | (css7-1)<br>R669* (CGA to TGA);<br>chr 2: 5642204 |  |  | This paper |  |
|  |  | CS72651 | cbp80-5 (css17-2) D57N<br>(D597N?) (GAT to AAT);<br>chr 2: 5641798 |  |  | This paper |  |
| **SUS2/PRP8a** | AT1G80070 | CS72652 | prp8-12 (css16-2)<br>Gly1820Arg (G1820R) (GGA to AGA);<br>chr 1:30125294 | GFP weak |  | This paper |  |
| **PRP18** | At1g03140 | n.a. | css40-1<br>G63R (GGA to AGA)<br>chr 1: 756037 | GFP weak |  | This paper |  |
| **T line (WT)** | n.a. | CS69640 |  | Wild-type GFP (intermediate) | Wild-type line expressing GFP reporter gene; used for EMS mutagenesis | Kanno et al., 2008 | **T line:**<br><u>small RNAs</u><br>SRR13267760<br>SAMN17103474<br>SRR13267719<br>SAMN17103475<br>SRR13267724<br>SAMN17103476 |
|  |  |  |  |  |  |  | control for all RNA-seq and DNA-seq experiments |
| **coilin** | At1g13030 | CS69632<br>CS69639 | hgf1-1/R40* (C118T)<br>hgf1-8/P439L (C2521T) | Hyper-GFP |  | Kanno et al., 2016<br>Kanno et al., 2016 | ***hgf1-1***:<br><u>small RNAs</u><br>SRR13267745<br>SAMN17103483<br>***hgf1-8***:<br><u>small RNAs</u><br>SRR13267720<br>SAMN17103484<br>SRR14917394<br>SAMN19865510<br>SRR14917406<br>SAMN19865511<br><u>RNAs</u><br>SRR14917408<br>SAMN19865501<br>SRR14917407<br>SAMN19865502<br>SRR14917401<br>SAMN19865503 |
| **CWC16a** | At1g25682 | CS69846, CS72366, CS72367 | cwc16a-1: (C to T, Chr 1:900354)<br>cwc16a-2: (C to T, Chr 1:9004043, W18*)<br>cwc16a-3: (C to T, Chr 1:9002886, splice site mutation) | Hyper-GFP |  | Kanno et al., 2017<br>This paper<br>This paper | ***cwc16a***:<br><u>small RNAs</u><br>SRR13267751<br>SAMN17103497<br>SRR13267726<br>SAMN17103498<br>***cwc16a-1/dcl4***<br><u>small RNAs</u><br>SRR13267727<br>SAMN17103499<br>SRR13267728<br>SAMN17103500 |
| **OTHER mutants used in this study** |  |  |  |  |  |  |  |
| **DCL2** | AT3G03300 | provided by Hervé Vaucheret | dcl2 -Kas |  |  | Parent et al. 2014 | ***dcl2/dcl4***:<br><u>small RNAs</u><br>SRR15185391<br>SAMN20298928 |
|  |  |  |  |  |  |  | SRR15185390<br>SAMN20298929 |
| **DCL3** | AT3G43920 | CS69179 | dcl3-5 |  |  | Daxinger et al.<br>2009 | ***dcl3/dcl4***:<br><u>small RNAs</u><br>SRR15185389<br>SAMN20298930<br>SRR15185388<br>SAMN20298931 |
| **DCL4** | AT5G20320 | provided<br>by Scott<br>Poething | dcl4-12: Ch5<br>C6866157T 8th<br>intron acceptor<br>site |  |  | provided by<br>Scott Poething | ***T/dcl4***:<br><u>small RNAs</u><br>SRR13267732<br>SAMN17103477<br>SRR13267738<br>SAMN17103478<br>SRR13267741<br>SAMN17103479<br>***hgf1-1/dcl4***:<br><u>small RNAs</u><br>SRR13267759<br>SAMN17103485<br>SRR13267758<br>SAMN17103486<br>SRR15185396<br>SAMN20298924<br>SRR15185395<br>SAMN20298925<br>***dcl4***:<br><u>small RNAs</u><br>SRR15185387<br>SAMN20298932<br>SRR15185386<br>SAMN20298933<br>SRR15185394<br>SAMN20298934 |
| **RDR6** |  |  | rdr6-14: Ch3<br>C18349667T<br>W1039* |  |  | Peragine et al.,<br>2004 | ***T/rdr6***:<br><u>small RNAs</u><br>SRR13267742<br>SAMN17103480<br>SRR13267743<br>SAMN17103481<br>SRR13267744 |
|  |  |  |  |  |  |  | SAMN17103482 |
|  |  |  |  |  |  |  | ***hgf1-1/rdr6:***<br>**<u>small RNAs</u>** |
|  |  |  |  |  |  |  | SRR13267757<br>SAMN17103487 |
|  |  |  |  |  |  |  | SRR13267721<br>SAMN17103488 |
|  |  |  |  |  |  |  | SRR15185393<br>SAMN20298926 |
|  |  |  |  |  |  |  | SRR15185392<br>SAMN20298927 |

**Study: SRP298335**

**Bioproject accession: PRJNA686012**

<u>References</u>

**Daxinger**, L., Kanno, T., Bucher, E., van der Winden, J., Naumann, U., Matzke, A. J., & Matzke, M. (2009). A stepwise pathway for biogenesis of 24-nt secondary siRNAs and spreading of DNA methylation. The EMBO journal, 28(1), 48–57. https://doi.org/10.1038/emboj.2008.260

**Kanno, T.**, Lin, W. D., Fu, J. L., Wu, M. T., Yang, H. W., Lin, S. S., Matzke, A. J., & Matzke, M. (2016). Identification of Coilin Mutants in a Screen for Enhanced Expression of an Alternatively Spliced GFP Reporter Gene in Arabidopsis thaliana. Genetics, 203(4), 1709–1720. https://doi.org/10.1534/genetics.116.190751

**Kanno, T.**, Lin, W. D., Fu, J. L., Matzke, A., & Matzke, M. (2017). A genetic screen implicates a CWC16/Yju2/CCDC130 protein and SMU1 in alternative splicing in Arabidopsis thaliana. RNA (New York, N.Y.), 23(7), 1068–1079. https://doi.org/10.1261/rna.060517.116

**Kanno, T.**, Bucher, E., Daxinger, L., Huettel, B., Böhmdorfer, G., Gregor, W., Kreil, D. P., Matzke, M., & Matzke, A. J. (2008). A structural-maintenance-of-chromosomes hinge domain-containing protein is required for RNA-directed DNA methylation. Nature genetics, 40(5), 670–675. https://doi.org/10.1038/ng.119

**Parent**, J.-S., Bouteiller, N., Elmayan, T. and Vaucheret, H. (2015), Respective contributions of Arabidopsis DCL2 and DCL4 to RNA silencing. Plant J, 81: 223-232. https://doi.org/10.1111/tpj.12720

**Peragine** A, Yoshikawa M, Wu G, Albrecht HL, Poethig RS. SGS3 and SGS2/SDE1/RDR6 are required for juvenile development and the production of trans-acting siRNAs in Arabidopsis. Genes Dev. 2004;18(19):2368-2379. doi:10.1101/gad.1231804

**Table S2:**
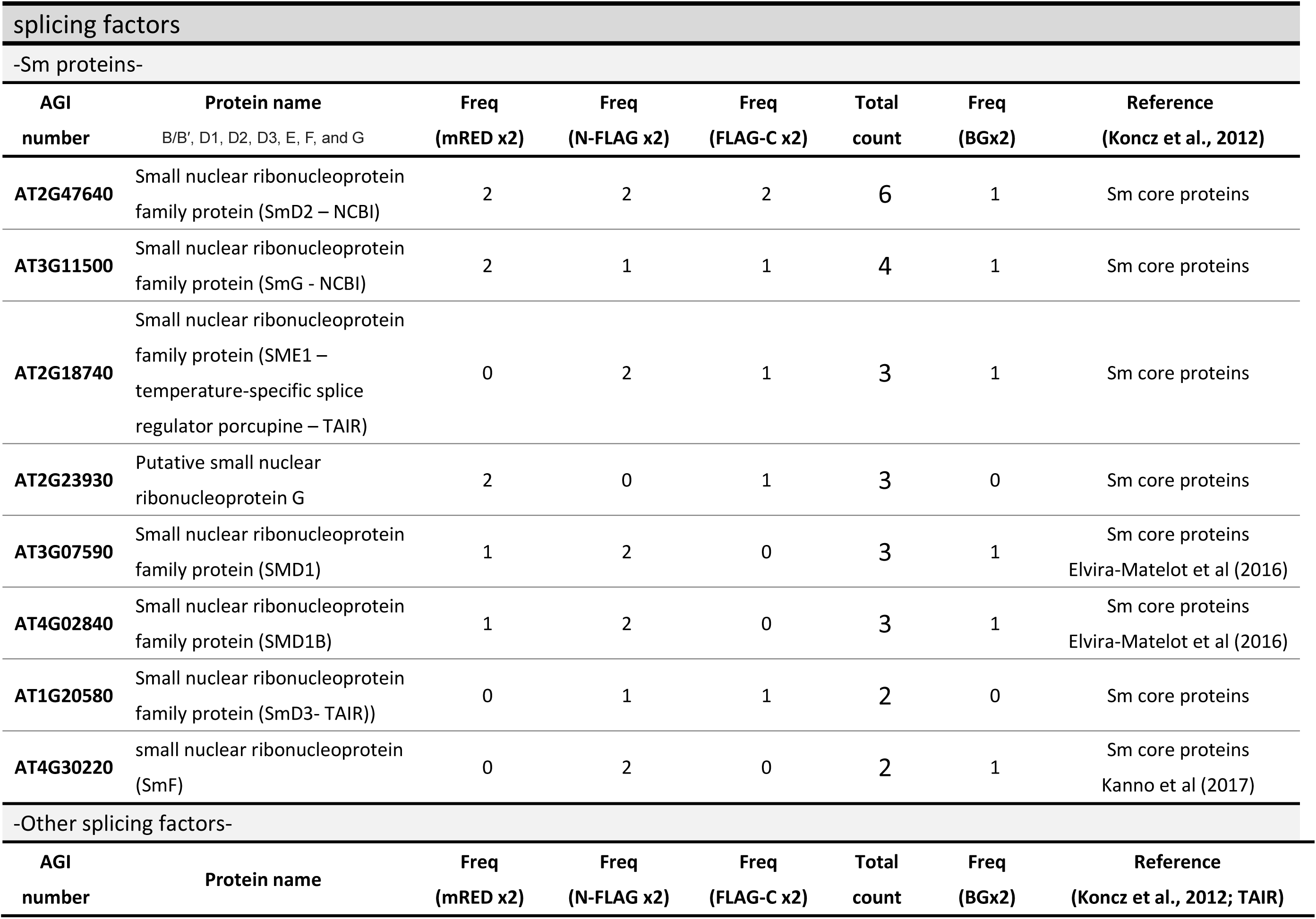
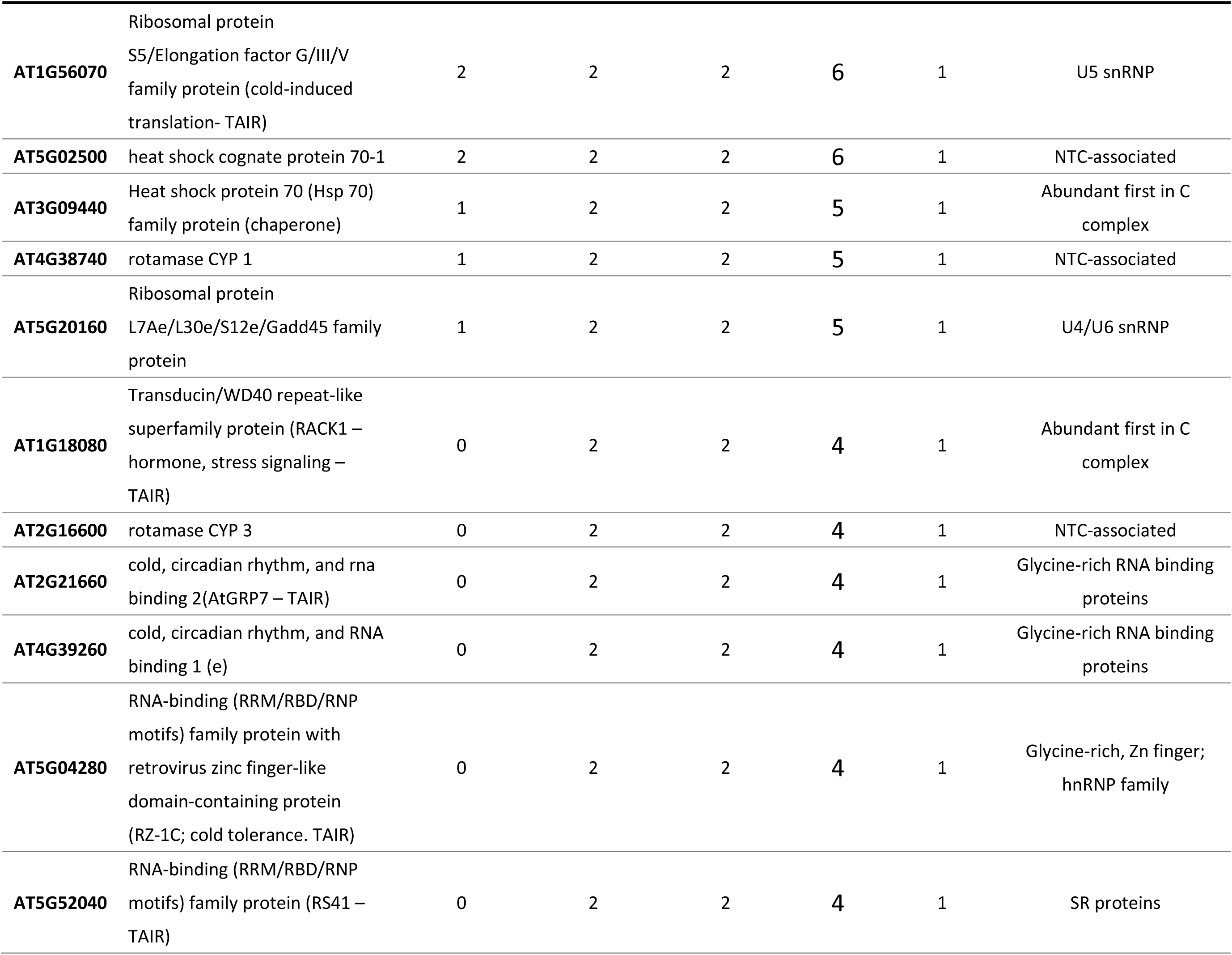
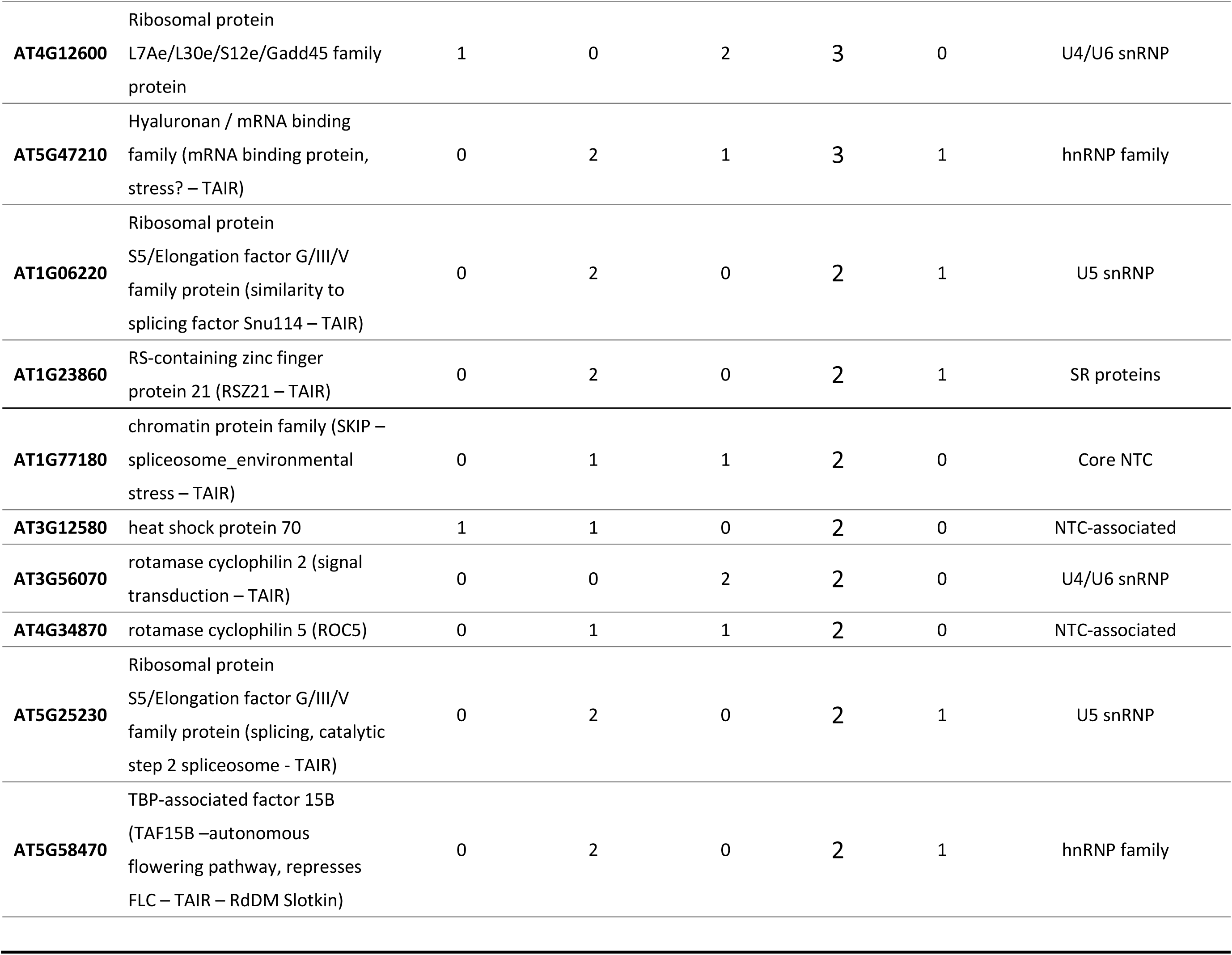
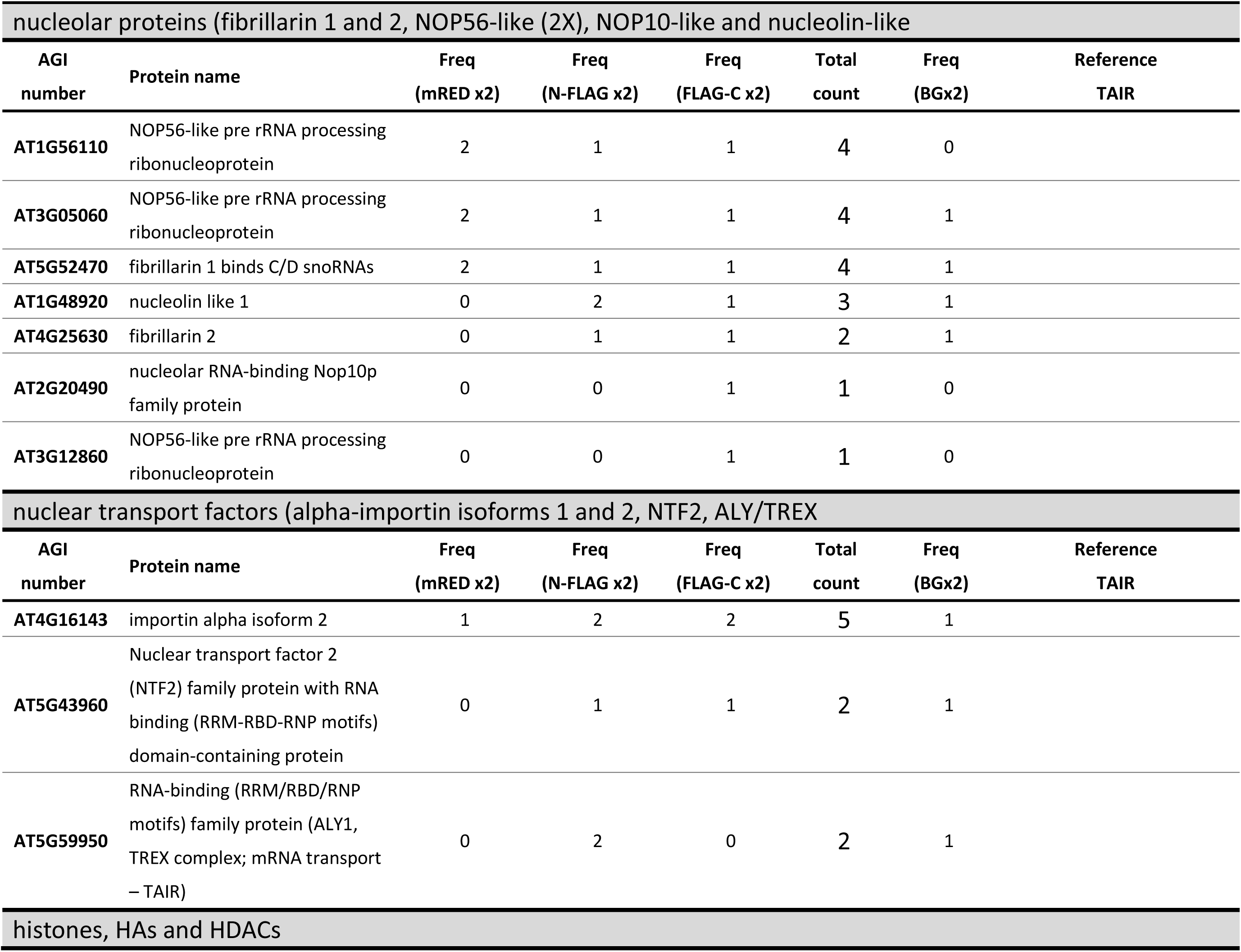
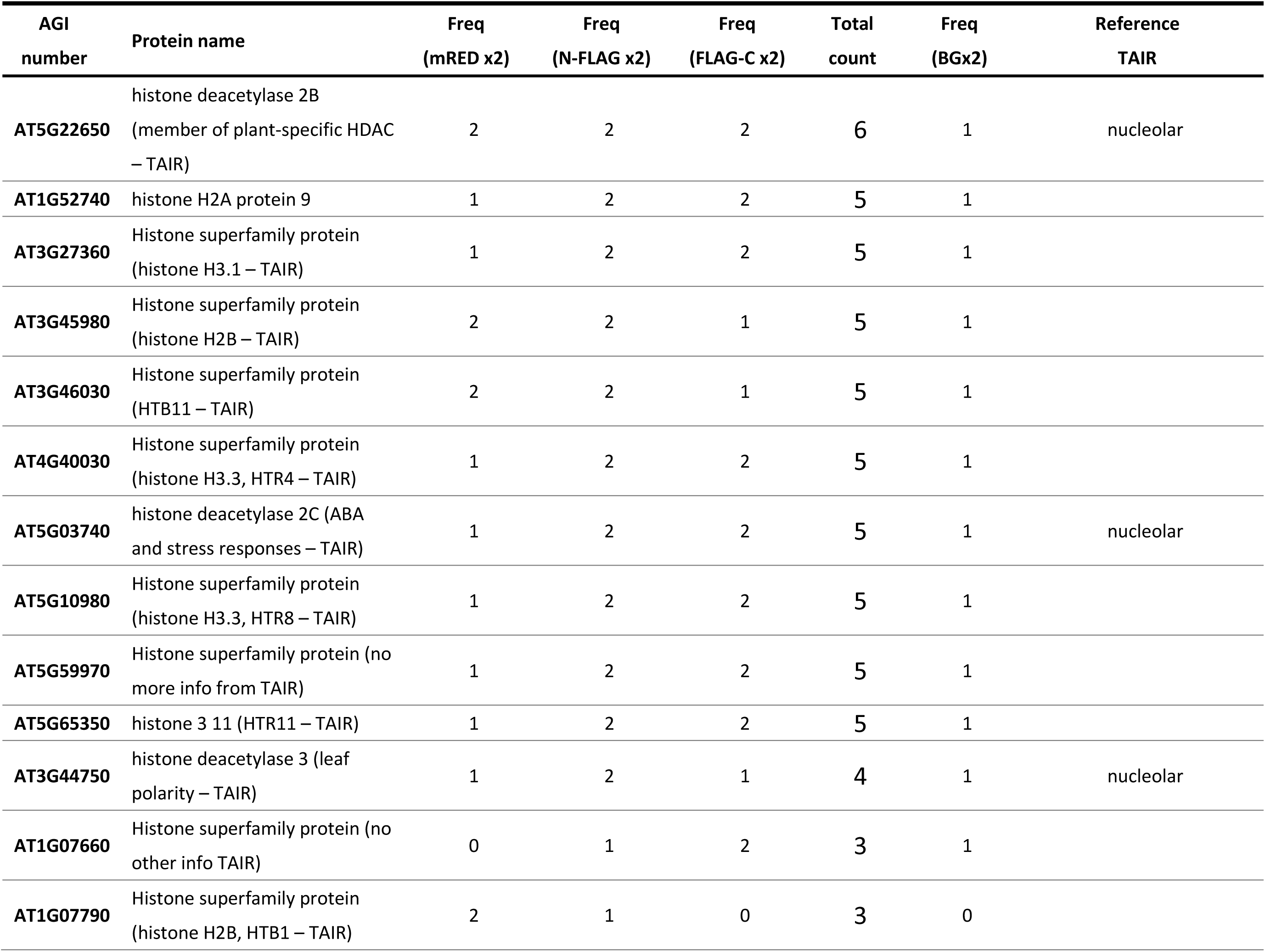
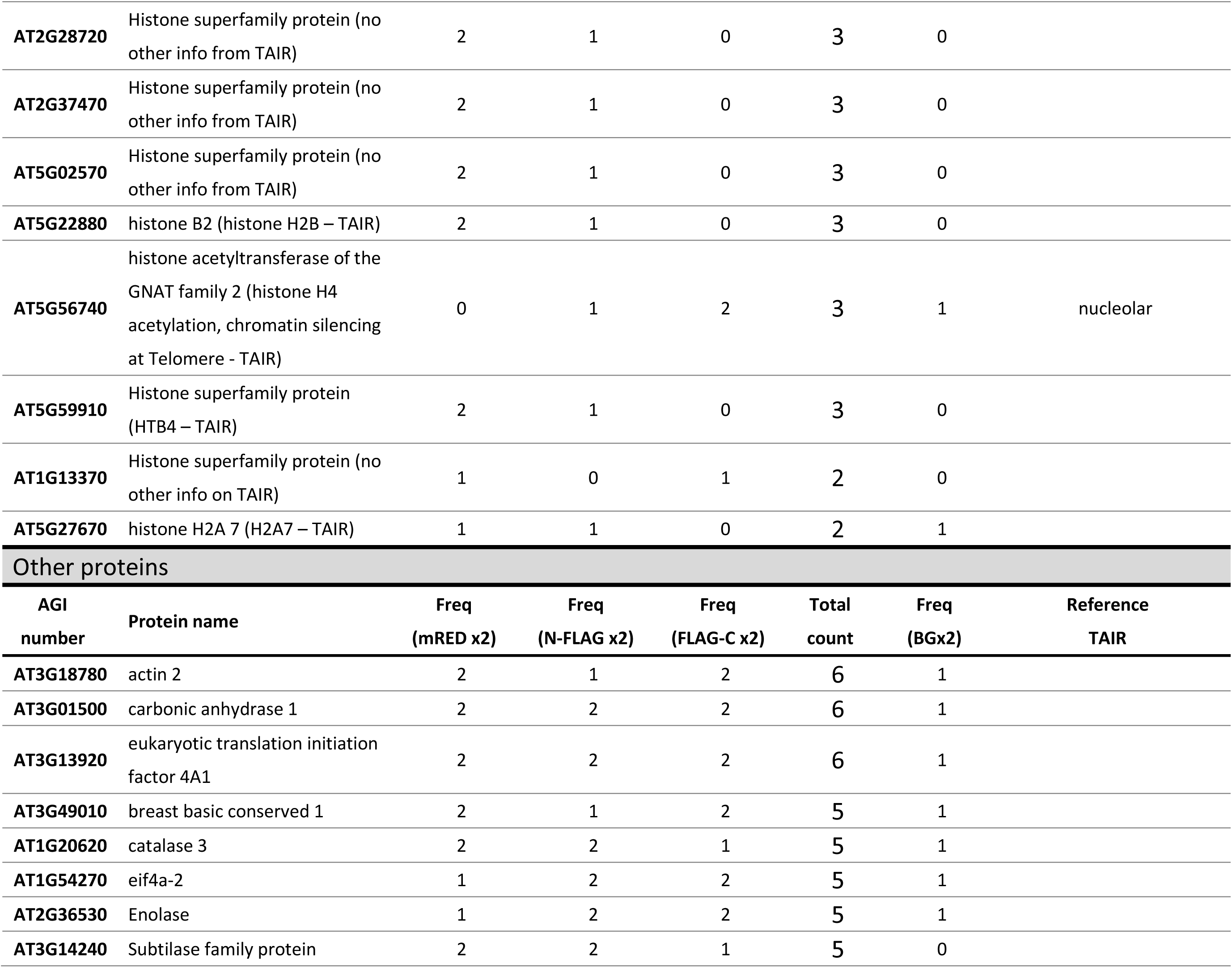
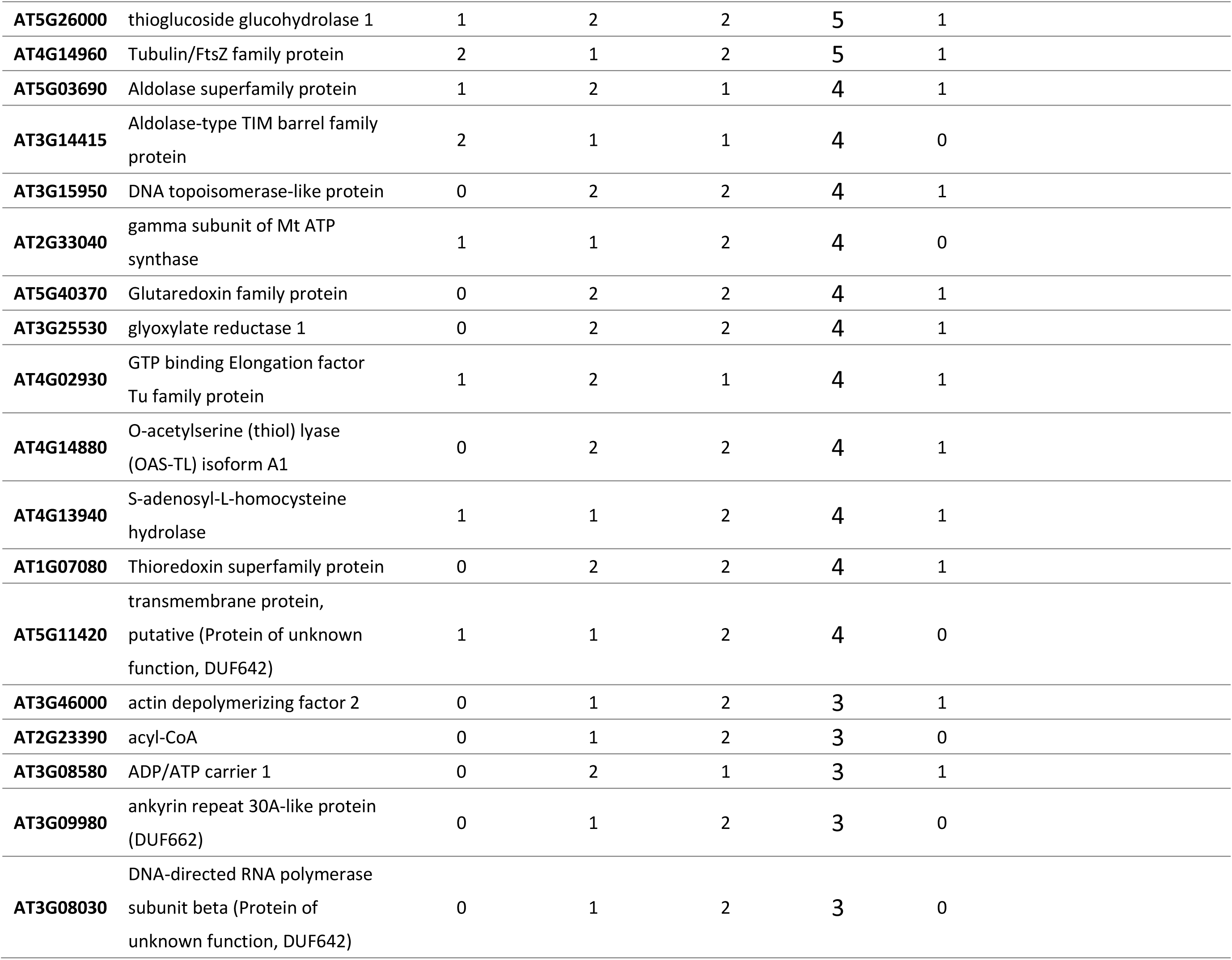
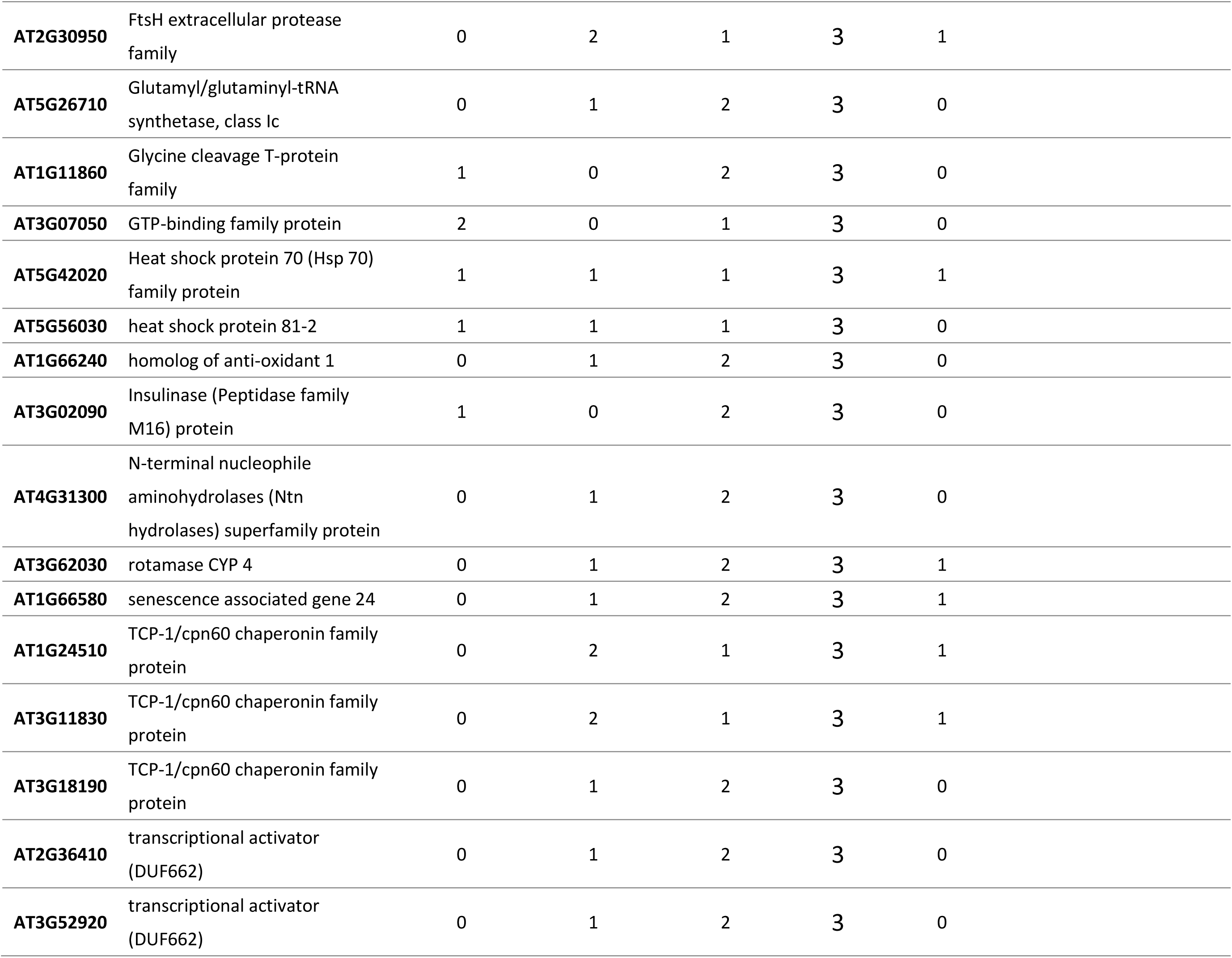
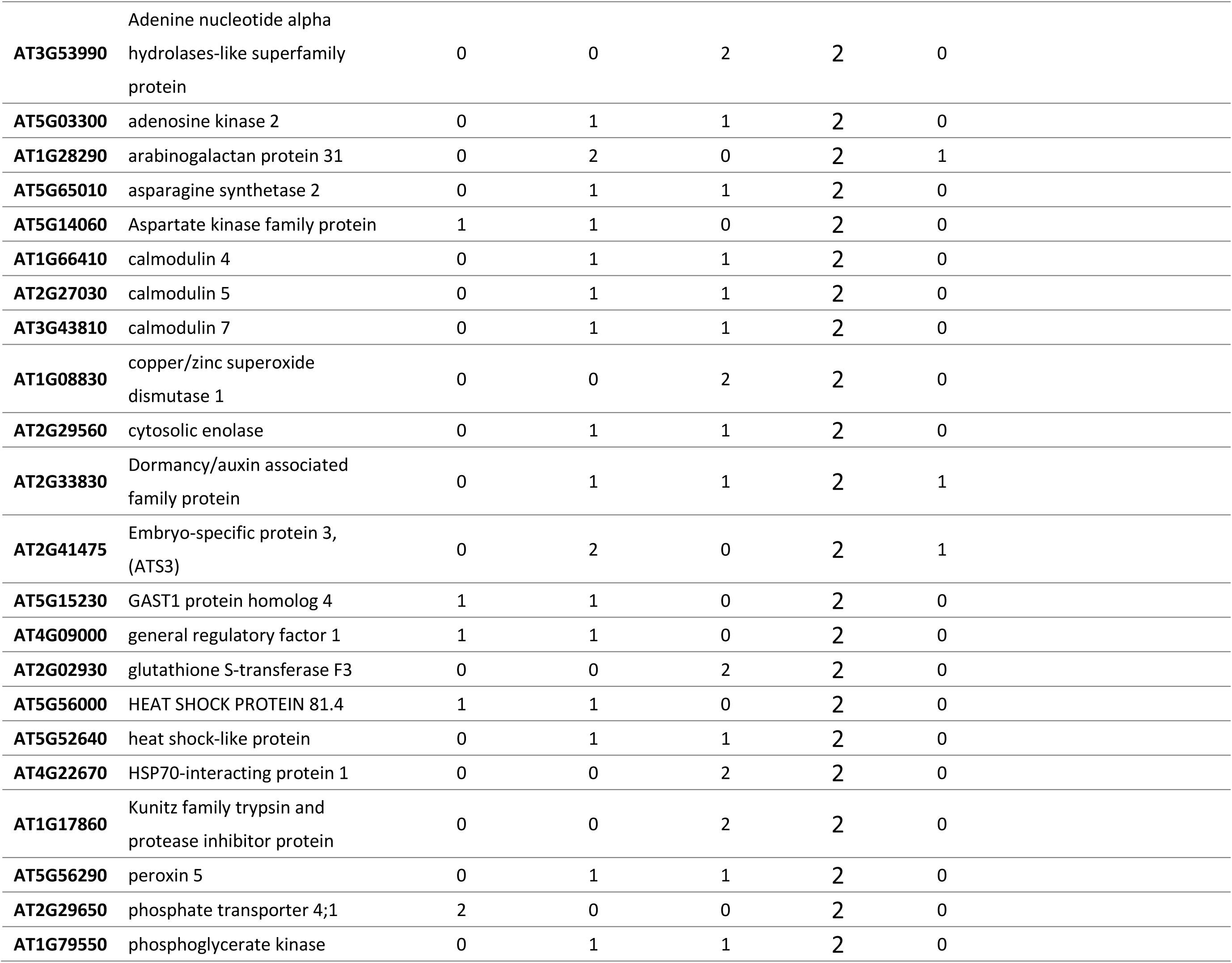
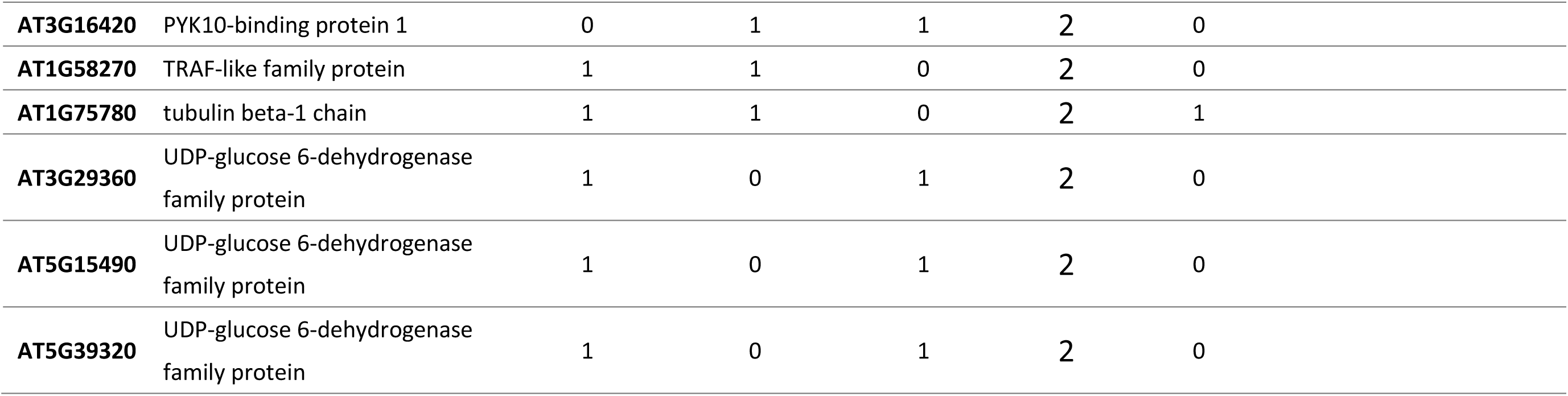
Summary of coilin IP-MS experiments.

Two independent experiments were performed for each of the three epitope-tagged coilin proteins (six experiments total). Frequency columns indicate the number of times a coilin interaction was detected for the indicated protein in the six experiments. The Freq BG indicates the number of times a protein was detected in two independent experiments (one for mRED, one for FLAG) without IP, i.e., background noise.

These finding are in accord with earlier findings showing that metazoan Sm proteins can interact with interact with the coilin CTD and several nucleolar proteins (Machnya et al., 2015; Xu et al., 2005). Abbreviation: TAIR – The Arabidopsis Information Resource (https://www.arabidopsis.org/), NTC – Nine Teen complex

**References**

Elvira-Matelot E, Bardou F, Ariel F, Jauvion V, Bouteiller N, Le Masson I, Cao J, Crespi MD, Vaucheret H (2016) The nuclear ribonucleoprotein SmD1 interplays with splicing, RNA quality control, and posttranscriptional gene silencing in Arabidopsis. Plant Cell 28:426-38. doi: 10.1105/tpc.15.01045.

Kanno T, Lin WD, Fu JL, Matzke AJM, Matzke M (2017) A genetic screen implicates a CWC16/Yju2/CCDC130 protein and SMU1 in alternative splicing in Arabidopsis thaliana. RNA 23:1068-1079.

Koncz, C., F. Dejong, N. Villacorta, D. Szakonyi, and Z. Koncz, 2012, The spliceosome-activating complex: molecular mechanisms underlying the function of a pleiotropic regulator. Front. Plant Sci. 3: 9. https://doi.org/10.3389/fpls.2012.00009

Machnya M, Neugebauer KM, Staněk D (2015) Coilin: the first 25 years. RNA Biology: 12: 590-596

Xu H, Pillai RS, Azzouz TN, Shpargel KB, Kambach C, Hebert MD, Schümperli D, Matera AG (2005) The C-terminal domain of coilin interacts with Sm proteins and U snRNPs. Chromosoma 114:155-166

**Table S3:** Summary of bisufite sequencing experiments to detect DNA methylation in the *GFP* upstream region.

**(A) *GFP***
| Genotype | methyated CG | unmethyated CG | methyated CHG | unmethyated CHG | methyated CHH | unmethyated CHH | # of clones |
| --- | --- | --- | --- | --- | --- | --- | --- |
| WT T | 11/110 (10.00) | 99/110 (90.00) | 0/120 (0.00) | 120/120 (100.00) | 29/461 (6.29) | 432/461 (93.71) | 10 |
| *coi1-1* | 11/110 (10.00) | 99/110 (90.00) | 0/120 (0.00) | 120/120 (100.00) | 29/461 (6.29) | 432/461 (93.71) | 10 |
| *coi1-1*<br>*zc3hc1* | 11/110 (10.00) | 99/110 (90.00) | 0/120 (0.00) | 120/120 (100.00) | 25/461 (5.42) | 436/461 (94.58) | 10 |
| *coi1-1*<br>*zc3hc1-4* | 11/110 (10.00) | 99/110 (90.00) | 3/120 (2.5) | 117/120 (97.5) | 28/467 (6.00) | 439/467 (94.00) | 10 |

**(B) *PHAVOLUTA***
| Genotype | methyated CG | unmethyated CG | methyated CHG | unmethyated CHG | methyated CHH | unmethyated CHH | # of clones |
| --- | --- | --- | --- | --- | --- | --- | --- |
| WT T | 4/117 (3.42) | 113/117 (96.58) | 1/162 (0.62) | 161/162 (99.38) | 8/456 (1.75) | 448/456 (98.25) | 9 |
| *coi1-1* | 0/130 (0.00) | 130/130 (100.00) | 0/180 (0.00) | 180/180 (100.00) | 12/508 (2.36) | 496/508 (97.64) | 10 |
| *coi1-1*<br>*zc3hc1-1* | 0/103 (0.00) | 103/103 (100.00) | 0/144 (0.00) | 144/144 (100.00) | 14/405 (3.46) | 391/405 (96.54) | 8 |
| *coi1-1*<br>*zc3hc11-4* | 0/117 (0.00) | 117/117 (100.00) | 0/162 (0.00) | 162/162 (100.00) | 13/458 (2.84) | 445/458 (97.16) | 9 |

**Legend:** The table shows the numbers of methylated and unmethylated cytosines and percentage of total in each of the indicated sequence contexts (CG. CHG, CHH) in *coi1* and *zc3hc1* mutants. The region tested for methylation in the *GFP* upstream region is indicated by the opposing blue arrows in Fig. 1, top. Previous work has shown that siRNAs direct to this region, which contains a short tandem repeat, can trigger TGS of the *GFP* reporter gene (Kanno et al., 2008). The *WT T* line is wild-type for *ZC3HC1* and *COI*. PHAVOLUTA is used as an unmethylated control (Bao et al, 2004; Reinders et al. (2008). Bisulfite sequencing as carried out as described in Daxinger et al (2009) using the primers listed below.

For GFP (top strand):

Top2F GCG GTG TYA TYT ATG TTA YTA GAT

Top2R CTT CTT RAT RTT CCA TAR CTT TCC

For PHV (this was performed by nested PCR)

Primary PCR

PHAV_P-F GTG YAG ATY TGT TTG GAG YTG ATT Y

PHAV_P-R TTT AAT ATC TAA CAT AAC CAA CCT TT

Secondary PCR

PHAV_S-F2 GGA YYA TAG TGA TGY YAT ATT GTG

PHAV_S-R TAT CAT CAA CAA CTT TCC ACA CC

**REFERENCES**

Bao N, Lye KW, Barton MK (2004) MicroRNA binding sites in Arabidopsis class III HD-ZIP mRNAs are required for methylation of the template chromosome. Dev Cell 7: 653–662

Daxinger L, Kanno T, Bucher E, van der Winden J, Naumann U, Matzke AJ, Matzke M (2009) A stepwise pathway for biogenesis of 24-nt secondary siRNAs and spreading of DNA methylation. EMBO J 28:48-57.

Kanno T, Bucher E, Daxinger L, Huettel B, Böhmdorfer G, Gregor W, Kreil DP, Matzke M, Matzke AJ (2008) A structural-maintenance-of-chromosomes hinge domain-containing protein is required for RNA-directed DNA methylation. Nat Genet 40:670-675.

Reinders J, Delucinge Vivier C, Theiler G, Chollet D, Descombes P, Paszkowski J (2008) Genome-wide, high-resolution DNA methylation profiling using bisulfite-mediated cytosine conversion. Genome Res 18: 469–476

**Table S4:**
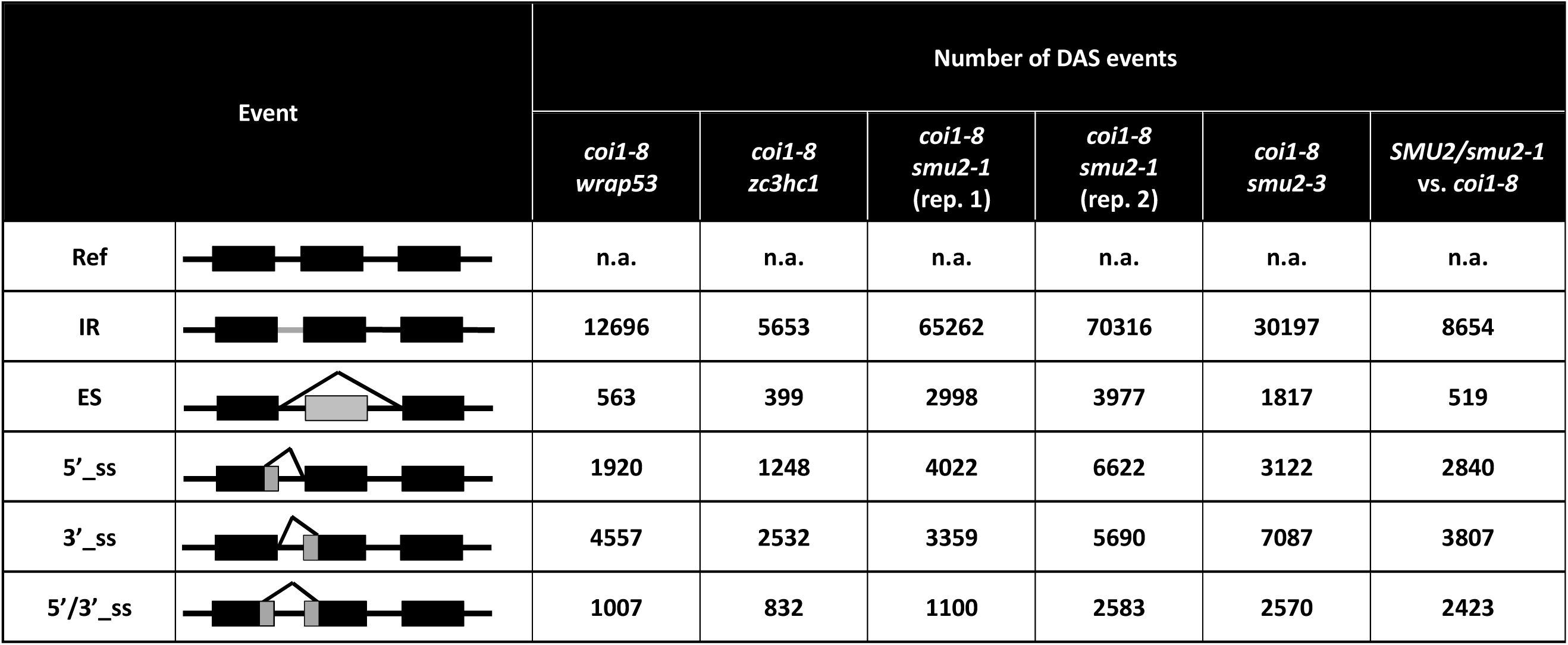
Summary of DAS events in smu2 mutants. For comparison to *smu2*, the DAS results for two other mutants identified in the *coi1-8* suppressor screen (*wrap53* and *zc3hc1*) are shown. The most striking changes in *smu2* mutants concern the number of IRs (roughly 10X more), ES (roughly 6X more) and 5’ss (roughly 3-4X more) compared to the other two mutants shown (*wrap53* and *zc3hc1*). The high number of IR events was observed in two independent experiments (reps) of *smu2-1* as well in a second allele, *smu2-3*. The high numbers of IRs return to a more ‘normal’ value when compared to *coi1-8* that is heterozygous for the *smu2-1* allele. **Abbreviations:** IR, intron retention; ES, exon skipping; 5’_ss, alternative 5’ splice site; 3’_ss, alternative 3’ splice site; 5’/3’_ss, alternative 5’ and 3’ splice sites

| Event |  | Number of DAS events |  |  |  |  |  |
| --- | --- | --- | --- | --- | --- | --- | --- |
|  |  | *coi1-8*<br>*wrap53* | *coi1-8*<br>*zc3hc1* | *coi1-8*<br>*smu2-1*<br>(rep. 1) | *coi1-8*<br>*smu2-1*<br>(rep. 2) | *coi1-8*<br>*smu2-3* | *SMU2/smu2-1*<br>vs. *coi1-8* |
| Ref |  | n.a. | n.a. | n.a. | n.a. | n.a. | n.a. |
| IR |  | 12696 | 5653 | 65262 | 70316 | 30197 | 8654 |
| ES |  | 563 | 399 | 2998 | 3977 | 1817 | 519 |
| 5'_ss |  | 1920 | 1248 | 4022 | 6622 | 3122 | 2840 |
| 3'_ss |  | 4557 | 2532 | 3359 | 5690 | 7087 | 3807 |
| 5'/3'_ss |  | 1007 | 832 | 1100 | 2583 | 2570 | 2423 |

